# Intertidal Gastropods (Gastropoda: Mollusca): Insights on diversity and distribution in the Mumbai Metropolitan Region, India

**DOI:** 10.1101/2025.11.20.689409

**Authors:** Raniya Momin Ansari, Pradip Patade, Shaunak Modi

## Abstract

Marine biodiversity documentation from the Mumbai Metropolitan Region (MMR) remains neglected despite the region having diversity of marine coastal habitats. The regions intertidal is one such habitat where species documentation remained heavily deficient due to lack of assessments and general apathy towards the habitat. This study addresses the issue of data deficiency of one of the largest taxa, Gastropoda through a decade long citizen science project, Marine Life of Mumbai. There exist large gaps in taxonomic research that have led to inconsistencies in species identification and inadequate ecosystem representation. This study addresses these issues by focusing on one of the largest taxa, the Molluscan class Gastropoda within the MMR. We present the spatial distribution of gastropod assemblages from 28 rocky, sandy and muddy intertidal sites within the Mumbai Metropolitan Region, on India’s west coast. A total of 163 species were recorded from 2164 observations of marine gastropods. Among these, 29 species, 34 genera and one family Limapontiidae are new records for the region. Additionally, this study reports rediscoveries of 7 species from their type locality, with 5 species of Heterobranchs recorded after 78 years: one species from Neogastropoda, *Lataxiena bombayana*, after 131 years and one from Siphonariida, *Siphonaria bassiensis* after 31 years, from their type locality. These species are herein illustrated with detailed morphological descriptions and their local distribution on 28 sites in the Mumbai Metropolitan Region. Through this study we elucidate that the citizen science efforts and the subsequent taxonomic analysis provide an effective and low-cost method for filling data gaps from large, understudied geographical areas.

## Introduction

Mollusca is the second-largest phylum of Kingdom Animalia, comprising 1,41,052 recognised species. Within it, Gastropoda is the largest and most heterogeneous class with 75,401 extant species of which 40,895 species are marine (Mollusca base, 2025; WoRMS, 2025). Marine gastropods are found across all marine interfaces, from the deep-sea trenches of the hadal region to the upper intertidal margins.

Intertidal gastropods represent a diverse taxonomic group that inhabit a dynamic ecotone where terrestrial and marine systems mutually influence each other. This has resulted in the intertidal zone being one of the most productive and biodiverse ecosystems in the world (Dorey, N. et. al., 2023). Gastropod assemblages in the intertidal are distributed vertically across upper, middle and lower zones; each characterised by varying degrees of exposure to air, water and selective environmental pressures from tidal fluctuations. This interplay between shore topography and tidal cycles creates complex assemblages adapted to diverse niches, responding to environmental gradients such as temperature, salinity, subaerial exposure and pH variation (Pawar et al., a, b, 2017; Sukumaran et al., 2021).

Despite their taxonomic diversity, ecological ubiquity and significance as bioindicators of environmental change, comprehensive baseline documentation of gastropod assemblages remains deficient; particularly in regions experiencing rapid urbanisation and anthropogenic disturbance. The Mumbai Metropolitan Region, on India’s west coast epitomises this problem (Pawar et al., a, b, 2017; Sukumaran et al., 2021).

Originally an archipelago of seven islands, Mumbai (erstwhile Bombay) was transformed into a single mainland city following extensive land reclamation beginning in the eighteenth century (Riding, 2018). Today, it ranks amongst the most densely populated metropolises in the world. The rapid growth in population and economic activity has placed significant anthropogenic stresses on the surrounding marine environment, intensifying the need for baseline biodiversity documentation (Vijay et al., 2016; Kamble & Vijay, 2016)

The history of malacological research from the region reveals a substantial knowledge deficit despite nearly two centuries of scientific attention. Most existing studies comprise classical natural history accounts (Leith, 1853; Abercrombie, 1893; Melvill, 1893; Melvill & Abercrombie, 1893; Melvill, 1894; Melvill, 1896), lacking morphological information necessary for reliable species identification (Melvill & Standen, 1901; Comber, 1906; Rai, 1933; Subrahmanyam et al., 1951, 1952; Apte, 1993) or represent discontinuous accounts across localised sites within the city (Winckworth, 1946a, b; Awati & Karandikar, 1948; Hornell, 1949; Bhatt & Bal, 1973; Kasinathan et al., 1975; Govindan et al., 1976; Sreeramamurthy, 1980; Kulkarni & Jaiswar, 2000; Kulkarni & Jaiswar, 2001; Jaiswar & Kulkarni, 2002; Datta et al., 2008; Prabhu & Kulkarni, 2012; Gavas et al., 2016; Kulkarni et al., 2017; Kantharajan et al., 2017; Gupta et al., 2019; Salvankar & Jadhav, 2021; Fatema et al., 2021; Mithbavkar et al., 2023; Shetty et al., 2024).

Lack of visual records, morphological descriptions and fine-scale spatial coverage in existing studies have generated incomplete and geographically scattered data. Temporal discontinuity in data collection has hindered quantification of long-term biodiversity shifts in response to environmental changes or anthropogenic pressures. Together, these have resulted in insufficient data for evidence-based conservation planning or a robust baseline for predicting future ecological changes in this highly anthropogenic ecosystem.

The present work addresses these gaps by presenting the first comprehensive checklist of gastropod diversity documented through a decade-long citizen science project, Marine Life of Mumbai. The project sought to explore and document the intertidal marine biodiversity along the MMR through community participation, generating an exhaustive dataset of georeferenced observations.

This work achieves two objectives: 1) consolidating photographic documentation of gastropod diversity compiled through citizen-led opportunistic intertidal surveys over a decade, 2) providing local distribution and photographic representation of the recorded gastropod diversity from the MMR.

The study also demonstrates citizen science as a viable methodology for a sustained, scalable landscape-scale biodiversity monitoring framework replicable in data-deficient ecosystems (Rosa et al., 2022; Callaghan et al., 2022).

## Methods

### Study Area

The Mumbai Metropolitan Region (19° 30’ N to 19°18’ N latitude and 72°42’ E to 72°57’ E longitude) situated on the west coast of India, covers an area of 6,328 km^2^ and consists of 9 municipal corporations, 9 municipal councils, and more than 1,000 villages in Thane, Raigad, and Palghar districts. The western seaboard is characterised by rocky shores, vast sandy shores and mangrove patches along creek mouths of four rivers draining into the Arabian sea. The eastern coastline is characterised by extensive mudflats and mangroves, along the Thane Creek. (MMRDA, 2023). The coastline is under the influence of the annual cycle of monsoon winds (May-September) and is subject to semi-diurnal tides with a period of 12 hours and 40 minutes. Data presented in this study was collected from 28 intertidal sites between Vasai and Colaba on the western seaboard and from 2 sites on the eastern coastline. These sites have been divided into 11 zones for ease of data presentation (Cárdenas Calle et al., 2020). The zones are elaborated and listed in Table 1 and visualised in Figure 1.

**Figure 1:**
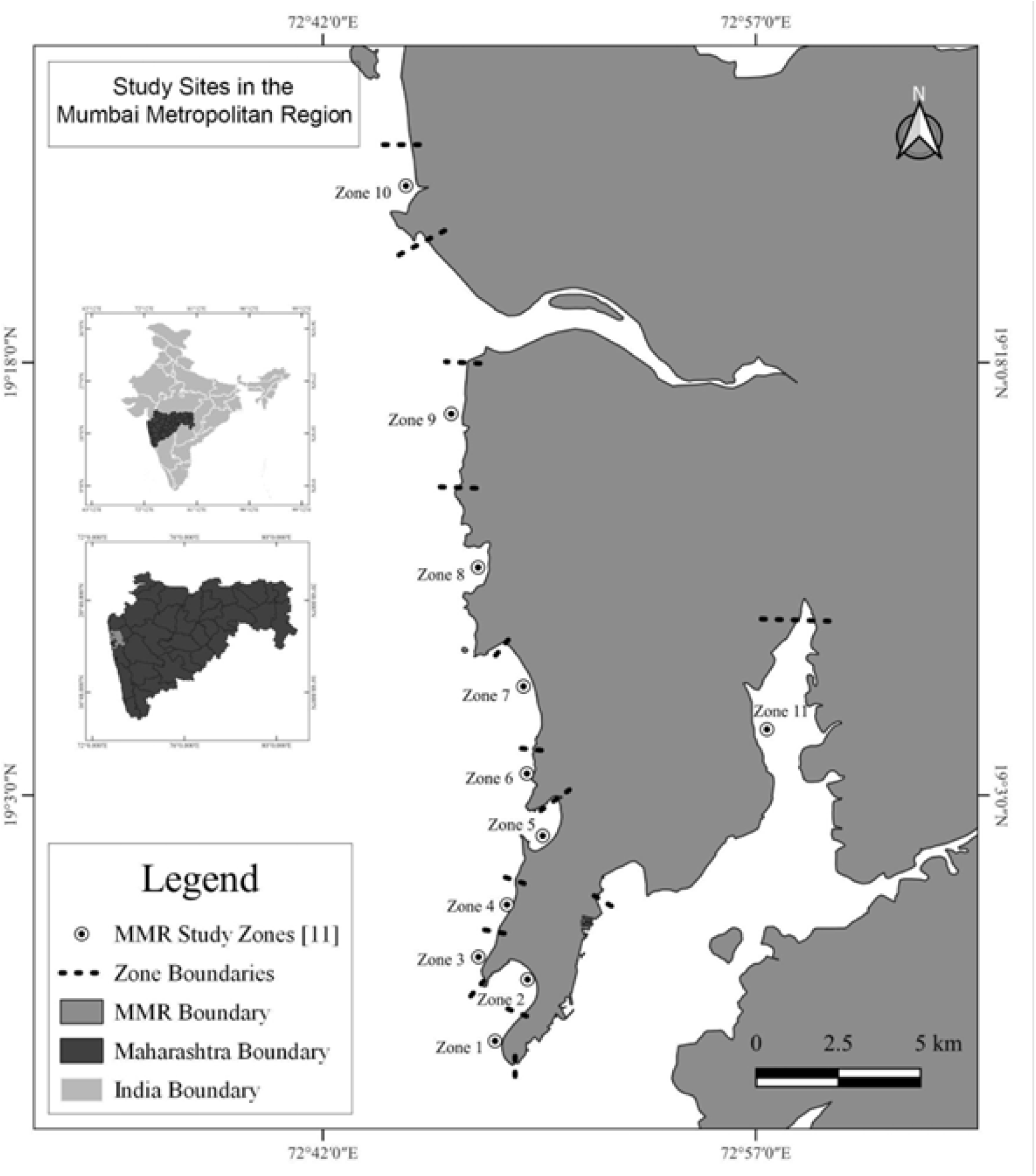
Map of MMR region showing the Study sites (Source: Authors).

**Table 1:** Study sites divided into Zones (Source: Authors).

| **Zone** | **Study Sites** | **District** |
| --- | --- | --- |
| Zone 1 | 1. Navy Nagar | Mumbai City |
|  | 2. Geeta Nagar | Mumbai City |
|  | 3. Cuffe Parade | Mumbai City |
| Zone 2 | 4. Marine Drive | Mumbai City |
|  | 5. Girgaon Chowpatty | Mumbai City |
| Zone 3 | 6. Walkeshwar | Mumbai City |
|  | 7. Nepean sea road | Mumbai City |
|  | 8. Priyadarshini Park | Mumbai City |
| Zone 4 | 9. Mahalaxmi | Mumbai City |
|  | 10. Haji Ali | Mumbai City |
| Zone 5 | 11. Worli | Mumbai City |
|  | 12. Dadar | Mumbai City |
|  | 13. Mahim | Mumbai City |
|  | 14. Mithi estuary | Mumbai City |
| Zone 6 | 15. Bandstand | Mumbai Suburban |
|  | 16. Chimbai | Mumbai Suburban |
|  | 17. Carter Road | Mumbai Suburban |
|  | 18. Khar Danda | Mumbai Suburban |
| Zone 7 | 19. Juhu Beach | Mumbai Suburban |
|  | 20. Versova | Mumbai Suburban |
| Zone 8 | 21. Madh | Mumbai Suburban |
|  | 22. Danapani | Mumbai Suburban |
|  | 23. Marve | Mumbai Suburban |
| Zone 9 | 24. Gorai Beach | Mumbai Suburban |
|  | 25. Gorai Jetty | Mumbai Suburban |
|  | 26. Uttan | Thane |
| Zone 10 | 27. Bhuigaon | Palghar |
| Zone 11 | 28. Sewri, Airoli | Mumbai City & Thane |

### Data Collection

Data were collected between February 2014 and January 2024 by 13 participants during citizen-led opportunistic intertidal surveys conducted at low tides below 1 metre. All intertidal marine biodiversity encountered during these surveys was documented.

Gastropods were photographed *in situ* using DSLR cameras equipped with macro lenses, submersible point-and-shoot cameras, and smartphones. For shelled gastropods, dorsal, ventral, and lateral views were photographed prior to returning specimens to their original or adjacent microhabitats. Shell-less and reduced-shell species were photographed without disturbance. For heterobranchs, photographs emphasising diagnostic morphological features including rhinophores, gills, cerata, and colour patterns were also captured to facilitate taxonomic identification.

Photographs were uploaded to iNaturalist.org and added into the ‘Marine Life of Mumbai’ project.

### Data Analysis

#### Literature Review

A bibliographic search was conducted using Google Scholar and Biodiversity Heritage Library. Eligibility criteria included any published papers about marine gastropods from the Mumbai Metropolitan Region published between 1800 and 30th January 2024. Grey literature and publications in regional languages were excluded from this review. The historical publications are enlisted in Table 2.

**Table 2:** List of Historical Molluscan Publications from the city.

| Research Publication | Year of Publication | Taxa | Type of study |
| --- | --- | --- | --- |
| 1. Leith | 1853 | Gastropoda | Classical taxonomy |
| 2. Abercrombie | 1893 | Gastropoda | Classical taxonomy, Diversity & Brief natural history accounts |
| 3. Melvill | 1893 | Gastropoda | Classical taxonomy |
| 4. Melvill & Abercrombie | 1893 | Mollusca | Classical taxonomy |
| 5. Melvill | 1894 | Gastropoda | Classical taxonomy |
| 6. Melvill | 1896 | Gastropoda | Classical taxonomy, Diversity & Distribution |
| 7. Melvill & Standen | 1901 | Mollusca | Classical taxonomy, Diversity & Distribution |
| 8. Comber | 1906 | Mollusca | Diversity & Distribution |
| 9. Rai | 1933 | Mollusca | Fisheries |
| 10. Winckworth | 1946 a | Heterobranchia | Classical taxonomy, Diversity & Distribution |
| 11. Winckworth | 1946 b | Heterobranchia | Classical taxonomy, Diversity & Distribution |
| 12. Awati & Karandikar | 1948 | Heterobranchia | Classical taxonomy, Diversity & Distribution |
| 13. Hornell | 1949 | Mollusca | Malacology |
| 14. Subrahmanyam et al. | 1951 | Gastropoda | Diversity & Distribution |
| 15. Subrahmanyam et al. | 1952 | Gastropoda | Diversity & Distribution |
| 16. Bhatt & Bal | 1973 | Animalia | Diversity & Distribution |
| 17. Kasinathan et al. | 1975 | Gastropoda | Reproductive biology |
| 18. Govindan et al. | 1976 | Animalia | Diversity & Abundance |
| 19. Sreeramamurthy | 1980 | Animalia | Ecology |
| 20. Apte | 1993 | Gastropoda | Diversity |
| 21. Kulkarni & Jaiswar | 2000 | Gastropoda | Diversity, Distribution & Ecology |
| 22. Kulkarni & Jaiswar | 2001 | Mollusca | Diversity & Distribution |
| 23. Jaiswar & Kulkarni | 2002 | Mollusca | Ecology |
| 24. Jaiswar & Kulkarni | 2005 | Mollusca | Diversity, Distribution & Conservation ecology |
| 25. Datta et al. | 2008 | Animalia | Diversity & Distribution |
| 26. Datta et al. | 2010 | Animalia | Diversity & Distribution |
| 27. Prabhu & Kulkarni | 2012 | Animalia & Plantae | Diversity & Local Distribution |
| 28. Sukumaran et al. | 2014 | Animalia | Marine Ecology & Anthropogenic Disturbance Impact |
| 29. Gavas et al. | 2016 | Heterobranchia | Classical taxonomy, Diversity & Distribution |
| 30. Kulkarni et al. | 2017 | Animalia | Diversity & Distribution |
| 31. Kantharajan et al. | 2017 | Mollusca | Diversity & Distribution |
| 32. Gupta et al. | 2019 | Mollusca | Diversity & Local Distribution |
| 33. Takar et al. | 2020 | Animalia | Diversity & Distribution |
| 34. Salvankar & Jadhav | 2021 | Mollusca | Diversity & Local Distribution |
| 35. Fatema et al. | 2021 | Animalia | Diversity & Distribution |
| 36. Mithbavkar et al. | 2023 | Mollusca | Diversity & Local Distribution |
| 37. Shetty et al. | 2024 | Animalia | Diversity & Local Distribution |

### Species Identification

Species were identified to the lowest attainable taxonomic level using morphological characters visible in the photographs and corroborated against published literature (Leith, 1853; Abercrombie, 1893; Melvill, 1893; Melvill & Abercrombie, 1893; Eliot, 1910; Rao, 1968; Bebbington, 1974; Gavas et al., 2016; Joshi & Mankodi, 2016; Apte & Desai, 2017; Gosliner et al., 2018; Venkitesan et al., 2019; Vakani et al., 2020). Identifications were subsequently validated by subject experts on the platform. For records with uncertain or provisional identifications, expert consultations were sought in online communities including Nudibase: Sharing Nudibranch Knowledge and Nudibranch Central or through taxonomic factsheets on the Sea Slug Forum. Reference websites, including www.keralamarinelife.in, www.wildsingapore.com, www.stromboidea.de were also used for verification.

Taxonomy and binomial nomenclature follow the World Register of Marine Species (WoRMS, 2025).

Identifications of all gastropod observations (n=2164) added to the ‘Marine Life of Mumbai’ project on iNaturalist.org (https://www.inaturalist.org/projects/marine-life-of-mumbai) were verified and IDs were corrected where necessary. The dataset was downloaded from iNaturalist.org and analysed for this study.

### Selection Criteria

For the final checklist, information was extracted from observations of live specimens photographed in their natural habitat. Information from observations of empty shells has been separately presented in Table 3. Observations with photos that lacked adequate diagnostic features were excluded. The compiled information included the taxonomic classification, IUCN status, accepted binomial name on WoRMS, original binomial name, and local distribution within the study area. Morphological diagnostic features, egg cases, and species ecology were noted down for each species.

**Table 3:** Record of Shelled Gastropods if they were observed live. (Source: Authors).

| Species Name | Encountered live | Encountered dead.<br>(only shell) | Earlier record from MMR |
| --- | --- | --- | --- |
| 1) *Aplysia cf. rudmani* | ✓ |  |  |
| 2) *Bursatella leachii* | ✓ |  |  |
| 3) *Bakawan rotundata* | ✓ |  |  |
| 4) *Haloa pemphis* | ✓ |  |  |
| 5) *Haloa wallisii* | ✓ |  |  |
| 6) *Smaragdinella sieboldi* | ✓ |  |  |
| 7) *Cylichna cylindracea* |  | ✓ | ✓ |
| 8) *Cassidula aurisfelis* | ✓ |  |  |
| 9) *Cassidula nucleus* | ✓ |  |  |
| 10) *Ellobium* sp. |  | ✓ | ✓ |
| 11) *Melampus sincaporensis* |  | ✓ | ✓ |
| 12) *Creseis* sp. | ✓ |  |  |
| 13) *Siphonaria basseinensis* | ✓ |  |  |
| 14) *Neripteron violaceum* | ✓ |  |  |
| 15) *Nerita albicilla* | ✓ |  |  |
| 16) *Nerita chamaeleon* | ✓ |  |  |
| 17) *Nerita oryzae* | ✓ |  |  |
| 18) *Clithon colombarium* | ✓ |  |  |
| 19) *Euchelus asper* | ✓ |  |  |
| 20) *Diodora lima* | ✓ |  |  |
| 21) *Diodora ruppellii* | ✓ |  |  |
| 22) *Scutus unguis* | ✓ |  |  |
| 23) *Trochus radiatus* | ✓ |  |  |
| 24) *Clanculus depictus* | ✓ |  |  |
| 25) *Umbonium vestiarium* | ✓ |  |  |
| 26) *Turbo intercostalis* | ✓ |  |  |
| 27) *Astracium semicostatum* | ✓ |  |  |
| 28) *Cellana radiata* | ✓ |  |  |
| 29) *Cellana rota* | ✓ |  |  |
| 30) *Patelloida lentiginosa* | ✓ |  |  |
| 31) *Planaxis sulcatus* | ✓ |  |  |
| 32) *Pirenella cingulata* | ✓ |  |  |
| 33) *Telescopium telescopium* | ✓ |  |  |
| 34) *Clypeomorus batillariaeformis* | ✓ |  |  |
| 35) *Melanoides tuberculata* | ✓ |  |  |
| 36) *Turritella duplicata* |  | ✓ | ✓ |
| 37) *Turritella* sp. |  | ✓ |  |
| 38) *Babylonia spirata* | ✓ |  |  |
| 39) Family: Muricidae - Unidentified | ✓ |  |  |
| 40) *Chicoreus brunneus* | ✓ |  |  |
| 41) *Chicoreus torrefactus* | ✓ |  |  |
| 42) *Murex* sp. | ✓ |  |  |
| 43) *Indothais lacera* | ✓ |  |  |
| 44) *Indothais sacellum* | ✓ |  |  |
| 45) *Indothais blanfordi* | ✓ |  |  |
| 46) *Purpura bufo* | ✓ |  |  |
| 47) *Purpura persica* | ✓ |  |  |
| 48) *Semiricinula tissoti* | ✓ |  |  |
| 49) *Reishia* cf. *bitubercularis* | ✓ |  |  |
| 50) *Ergalatax contracta* | ✓ |  |  |
| 51) *Lataxiena bombayana* |  | ✓ | ✓ |
| 52) *Arakawania granulata* | ✓ |  |  |
| 53) *Turricula javana* | ✓ |  |  |
| 54) *Conus achatinus* | ✓ |  |  |
| 55) *Conus biliosus* | ✓ |  |  |
| 56) *Conus hyaena* | ✓ |  |  |
| 57) *Conasprella dictator* |  | ✓ |  |
| 58) *Annulaturreis amicta* |  | ✓ | ✓ |
| 59) Family: Turridae-Unidentified |  | ✓ |  |
| 60) *Duplicaria spectabilis* | ✓ |  |  |
| 61) *Anachis terpsichore* | ✓ |  |  |
| 62) Family: Columbellidae-Unidentified | ✓ |  | - |
| 63) *Volegalea cochlidium* | ✓ |  |  |
| 64) *Nassarius stolatus* | ✓ |  |  |
| 65) *Nassarius jacksonianus* | ✓ |  |  |
| 66) *Nassarius foveolatus* | ✓ |  |  |
| 67) *Nassarius nodiferus* | ✓ |  |  |
| 68) *Bullia* sp. |  | ✓ | ✓ |
| 69) *Cantharus spiralis* | ✓ |  |  |
| 70) *Scalptia scalarina* | ✓ |  |  |
| 71) *Pseudonebularia*<br>*proscissa* | ✓ |  |  |
| 72) *Agaronia gibbosa* |  | ✓ | ✓ |
| 73) *Mauritia arabica* | ✓ |  |  |
| 74) *Monetaria moneta* |  | ✓ | ✓ |
| 75) *Naria turdus* | ✓ |  |  |
| 76) *Erronea pallida* | ✓ |  |  |
| 77) *Palmadusta lentiginosa* | ✓ |  |  |
| 78) *Cuspidolva* sp. | ✓ |  |  |
| 79) *Primovula* sp. | ✓ |  |  |
| 80) *Crenavolva striatula* | ✓ |  |  |
| 81) *Lamellaria* sp. | ✓ |  |  |
| 82) *Littoraria* sp. 1 | ✓ |  |  |
| 83) *Littoraria* sp. 2 | ✓ |  |  |
| 84) *Littoraria* sp. 3 | ✓ |  |  |
| 85) *Littoraria* sp. 4 | ✓ |  |  |
| 86) *Echinolittorina malaccana* | ✓ |  |  |
| 87) *Paratectonatica tigrina* | ✓ |  |  |
| 88) *Tanea lineata* | ✓ |  |  |
| 89) *Neverita didyma* | ✓ |  |  |
| 90) *Tibia curta* | ✓ |  |  |
| 91) *Bufonaria echinata* |  | ✓ | ✓ |
| 92) *Gyrineum natator* | ✓ |  |  |
| 93) *Assimineia marginata* | ✓ |  |  |
| 94) *Assimineia subconica* | ✓ |  |  |
| 95) *Iravadia bombayana* | ✓ |  |  |
| 96) *Thylacodes* sp.1 | ✓ |  |  |
| 97) *Thylacodes* sp.2 | ✓ |  |  |
| 98) *Thylacodes* sp. 3 | ✓ |  |  |
| 99) *Ergaea walshi* | ✓ |  |  |
| 100) *Janthina* sp. |  | ✓ | ✓ |

## Results

We report a total of 163 gastropod records belonging to 112 genera, 66 families and 5 subclasses from 28 locations within the Mumbai Metropolitan Region (Table 4). Additionally, 64 new records are reported from Mumbai, comprising 29 new species records, 34 new genus records, one new family Limapontiidae has been recorded (Table 5).

**Table 4:**
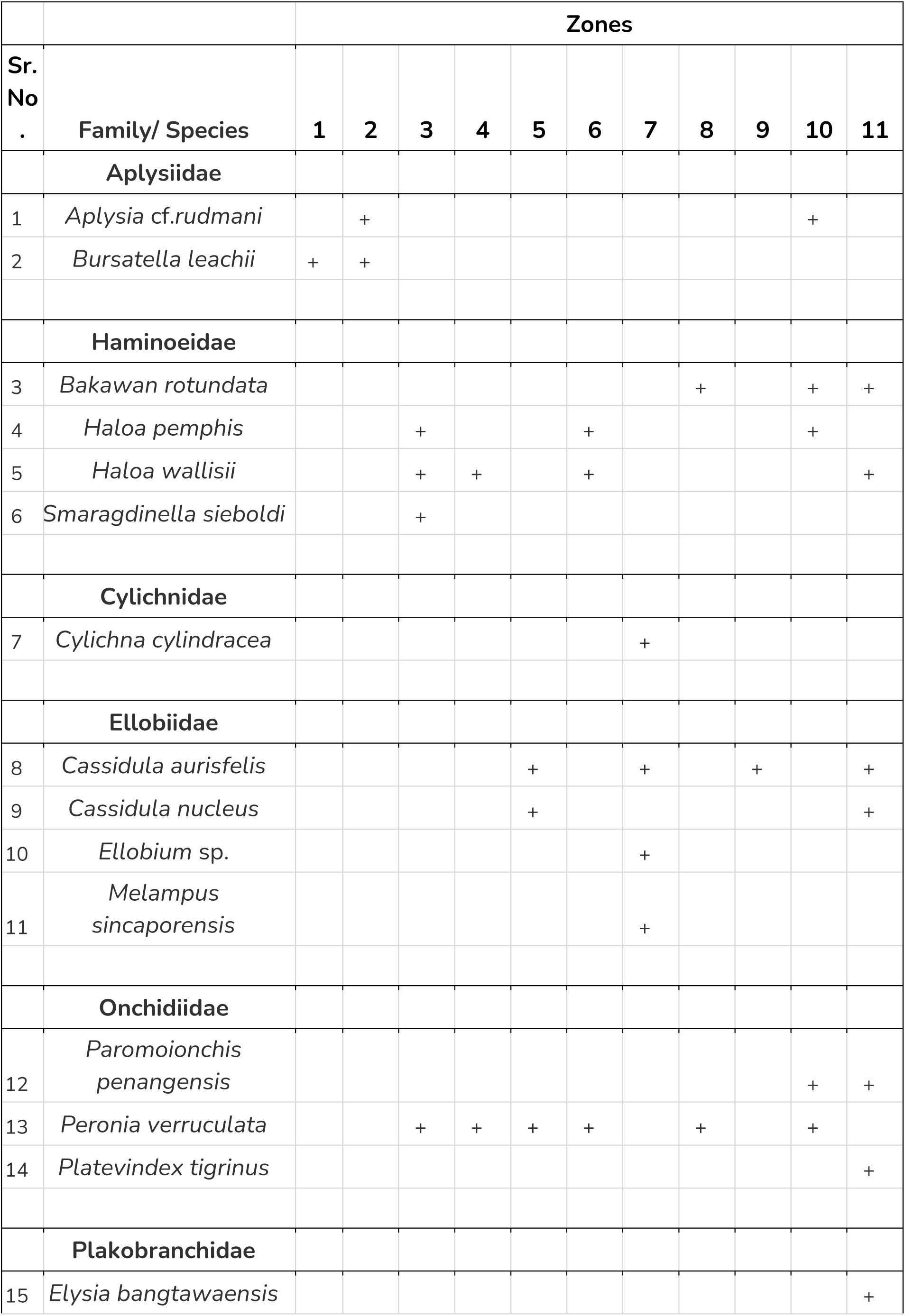
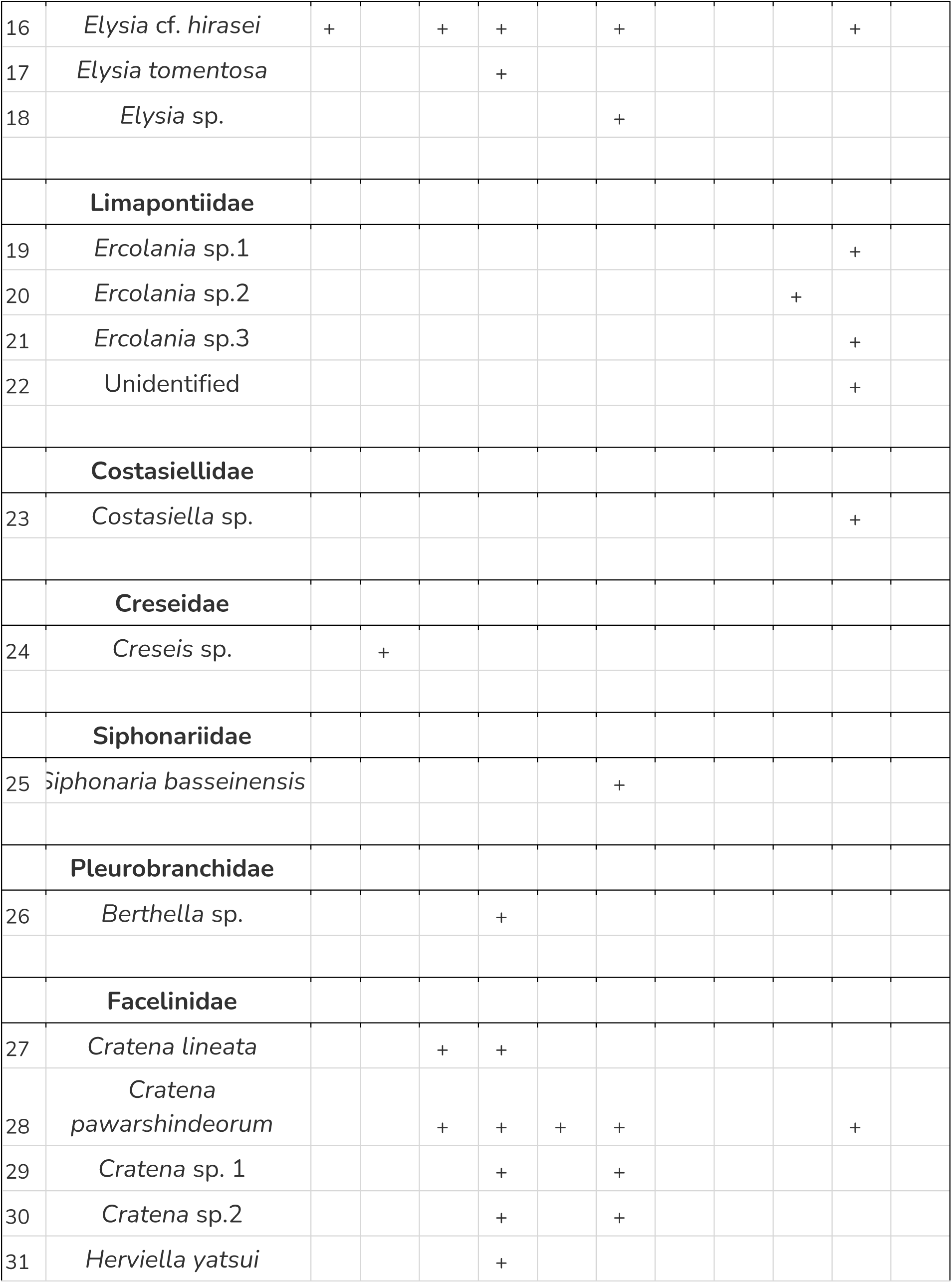
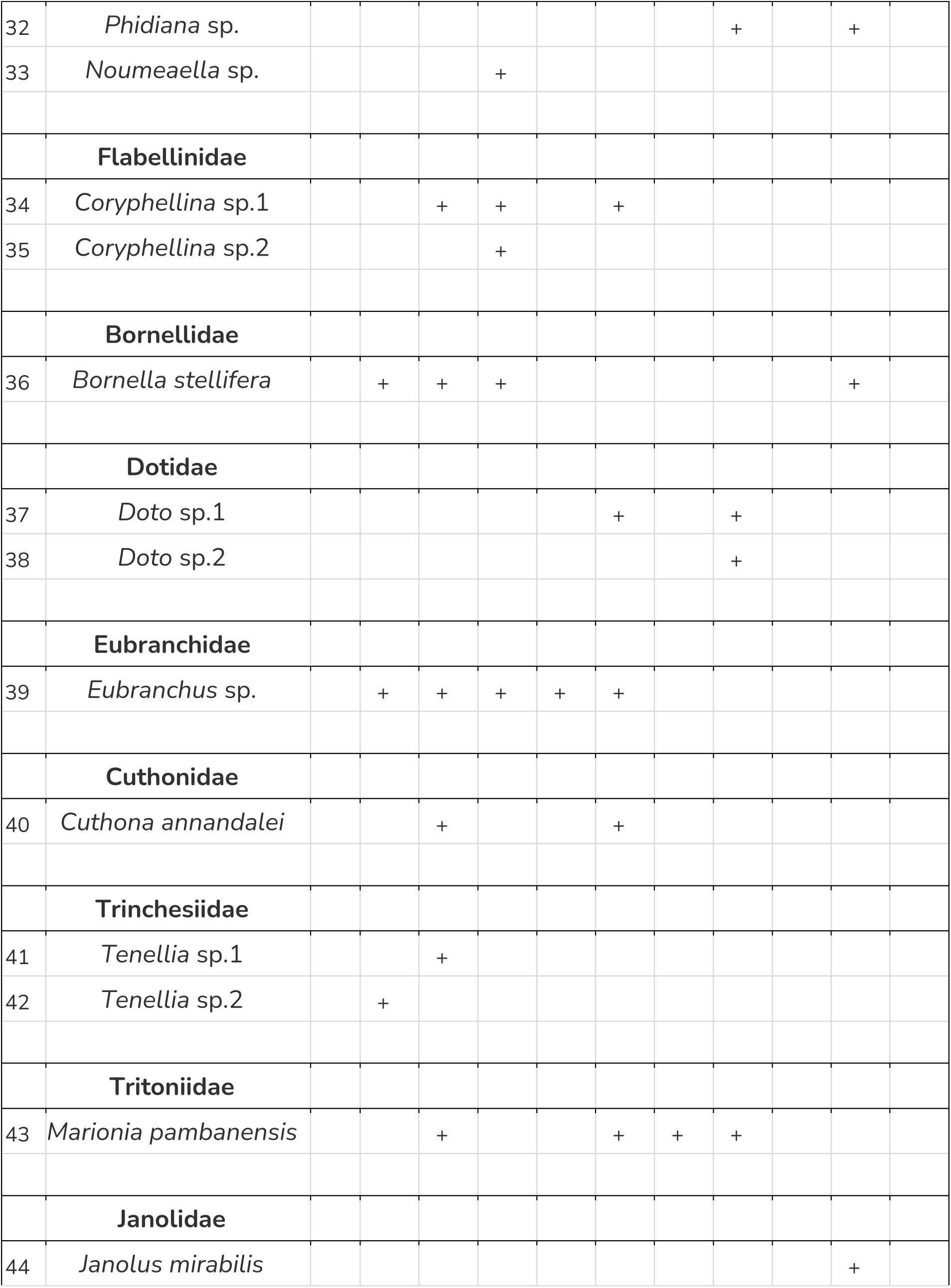
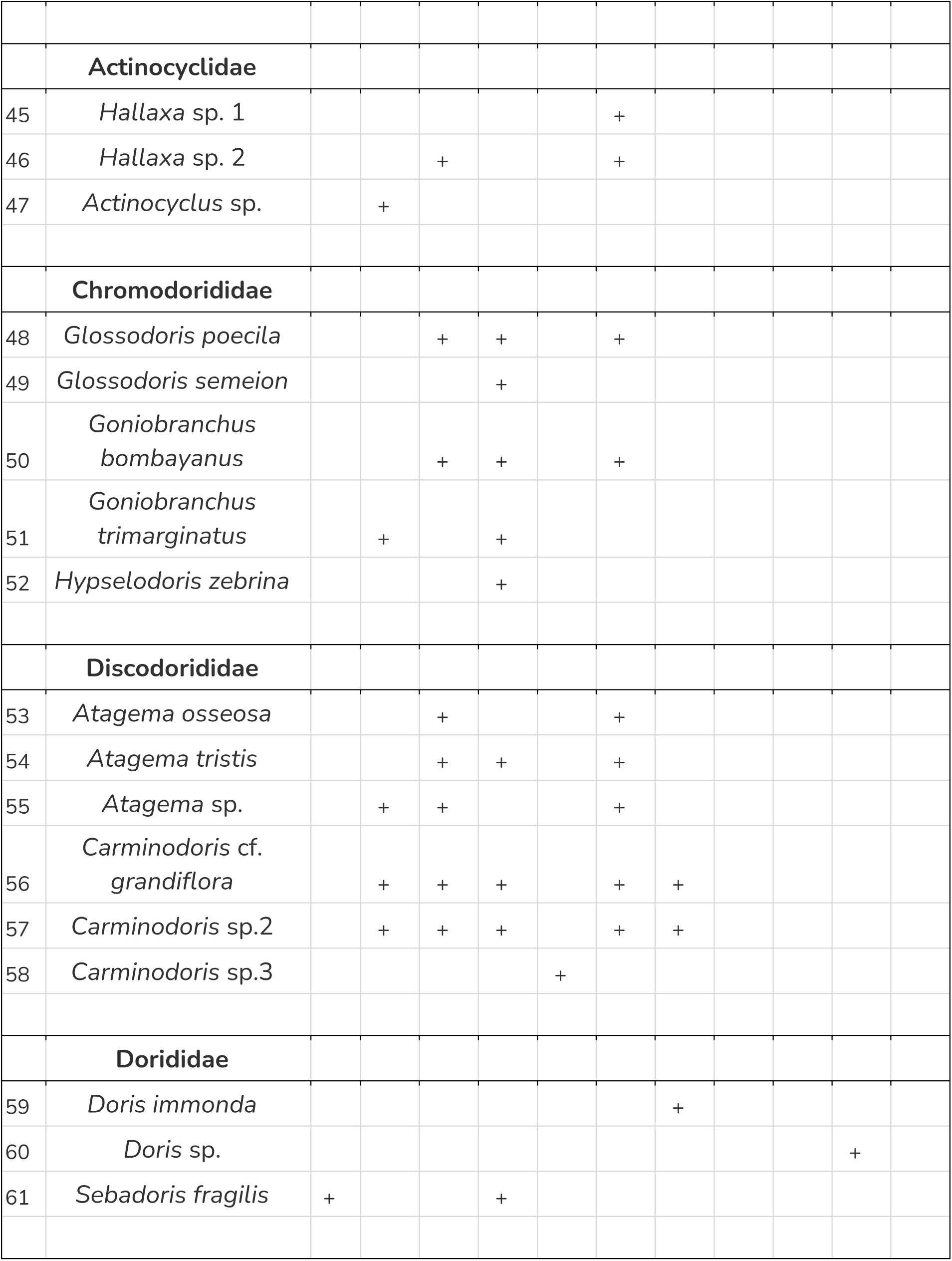
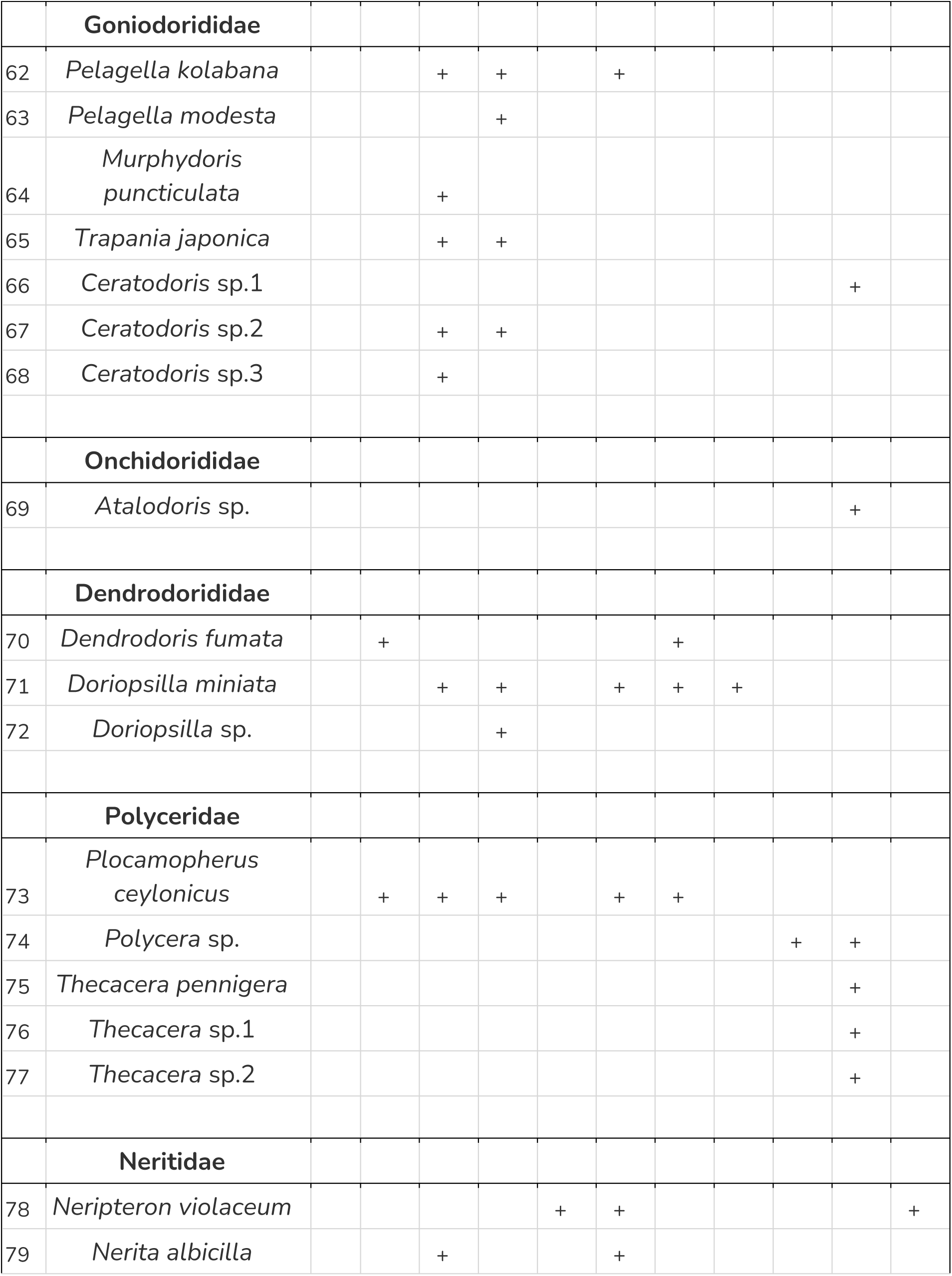
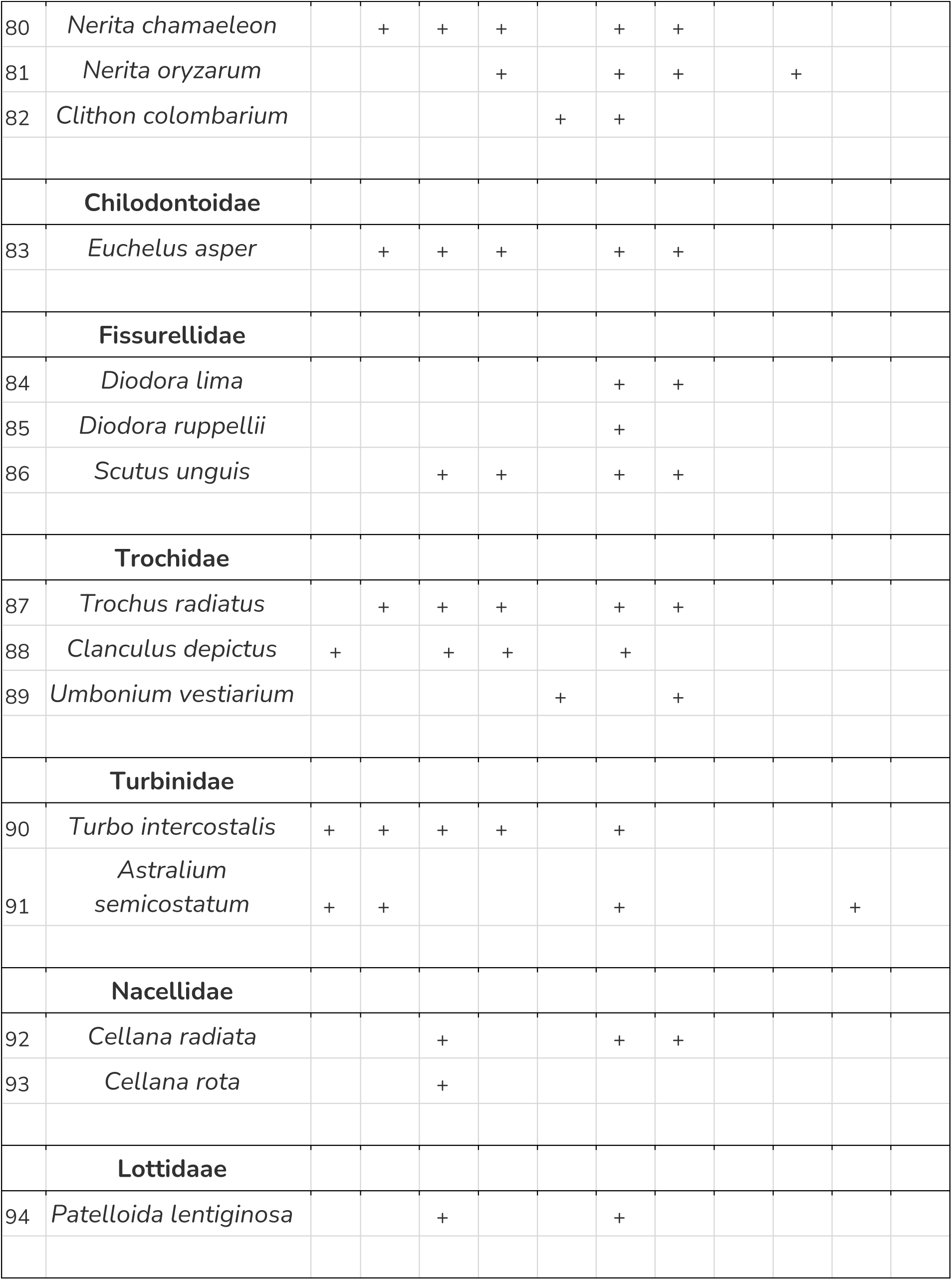
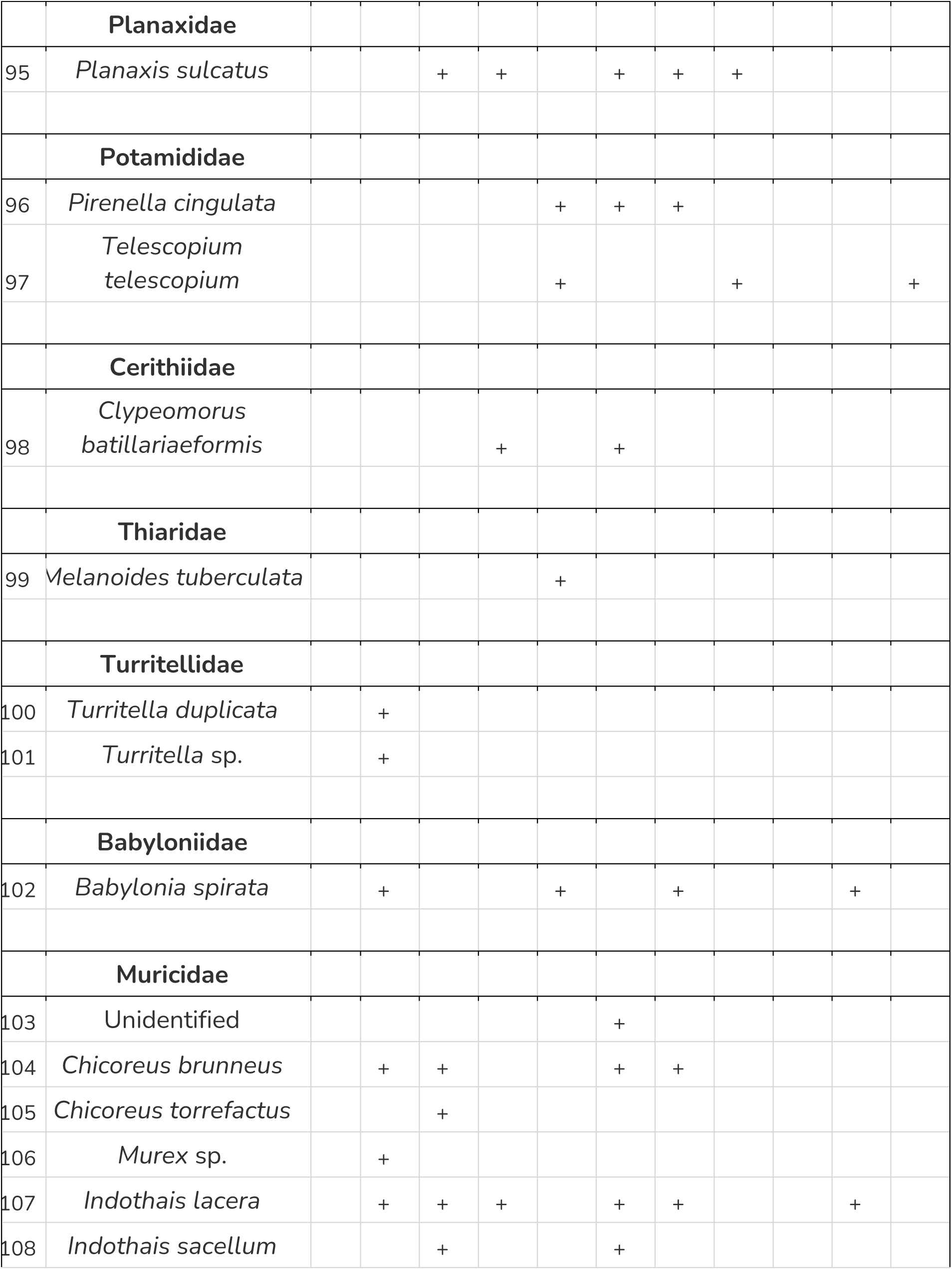
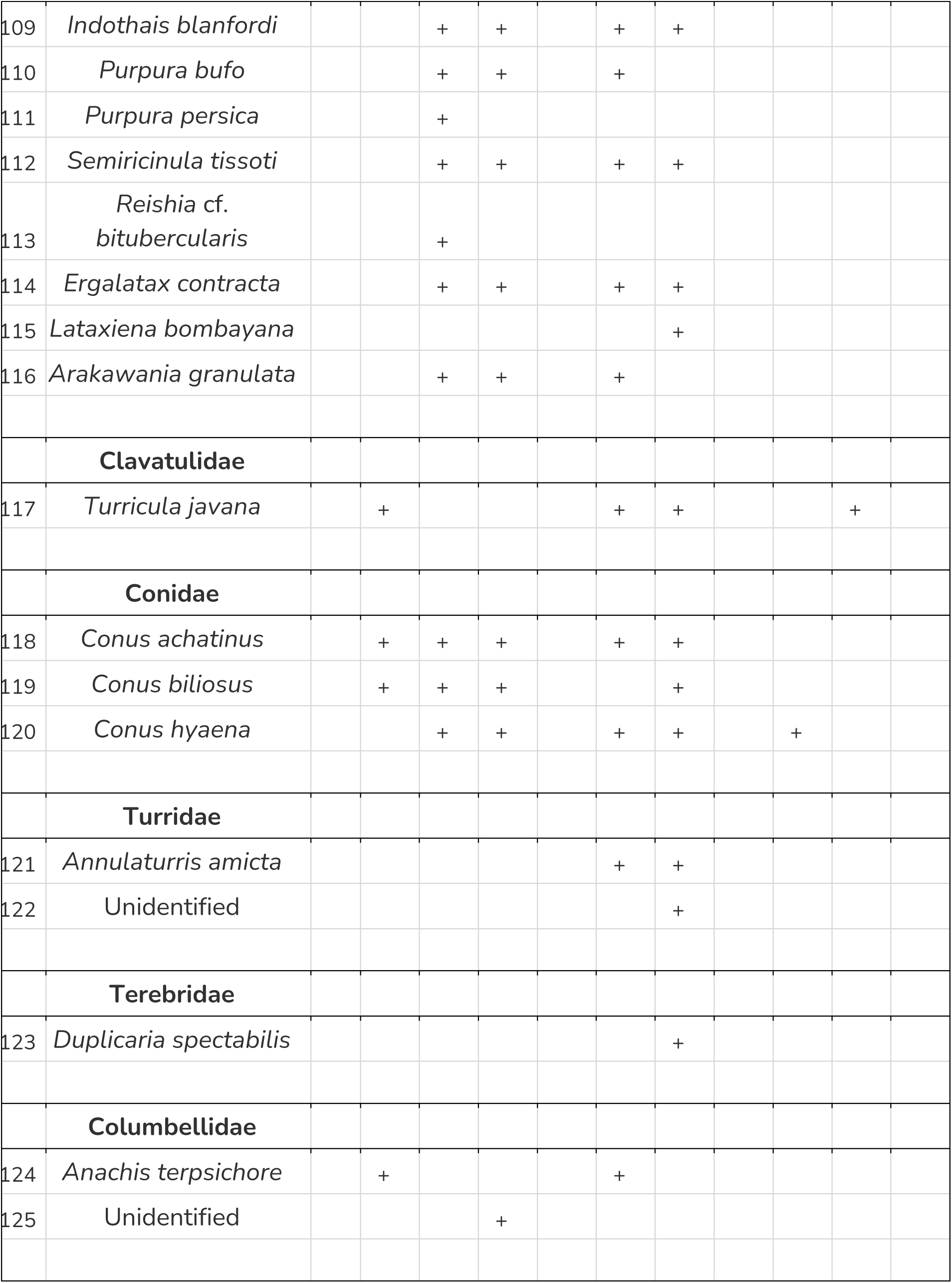
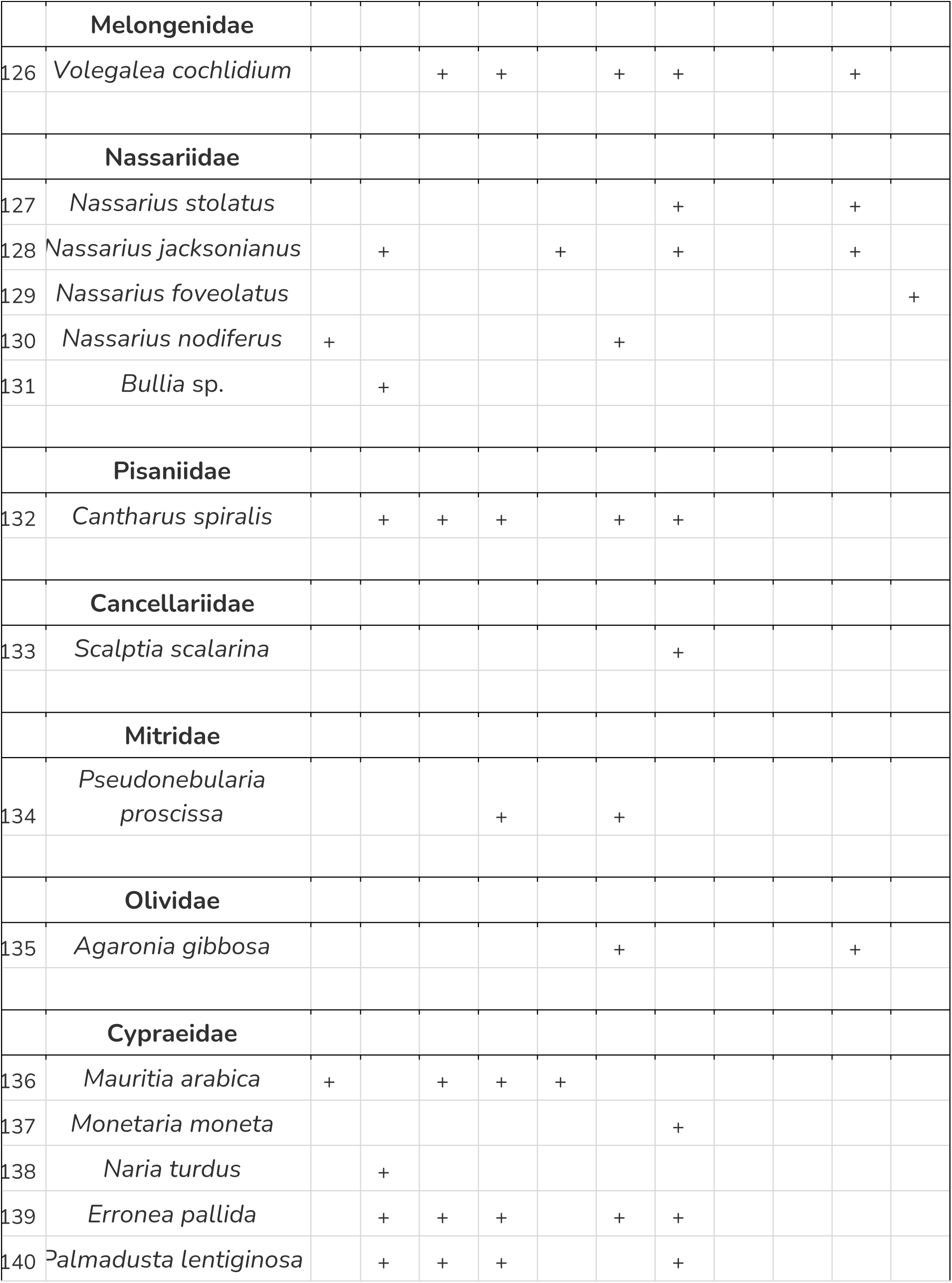
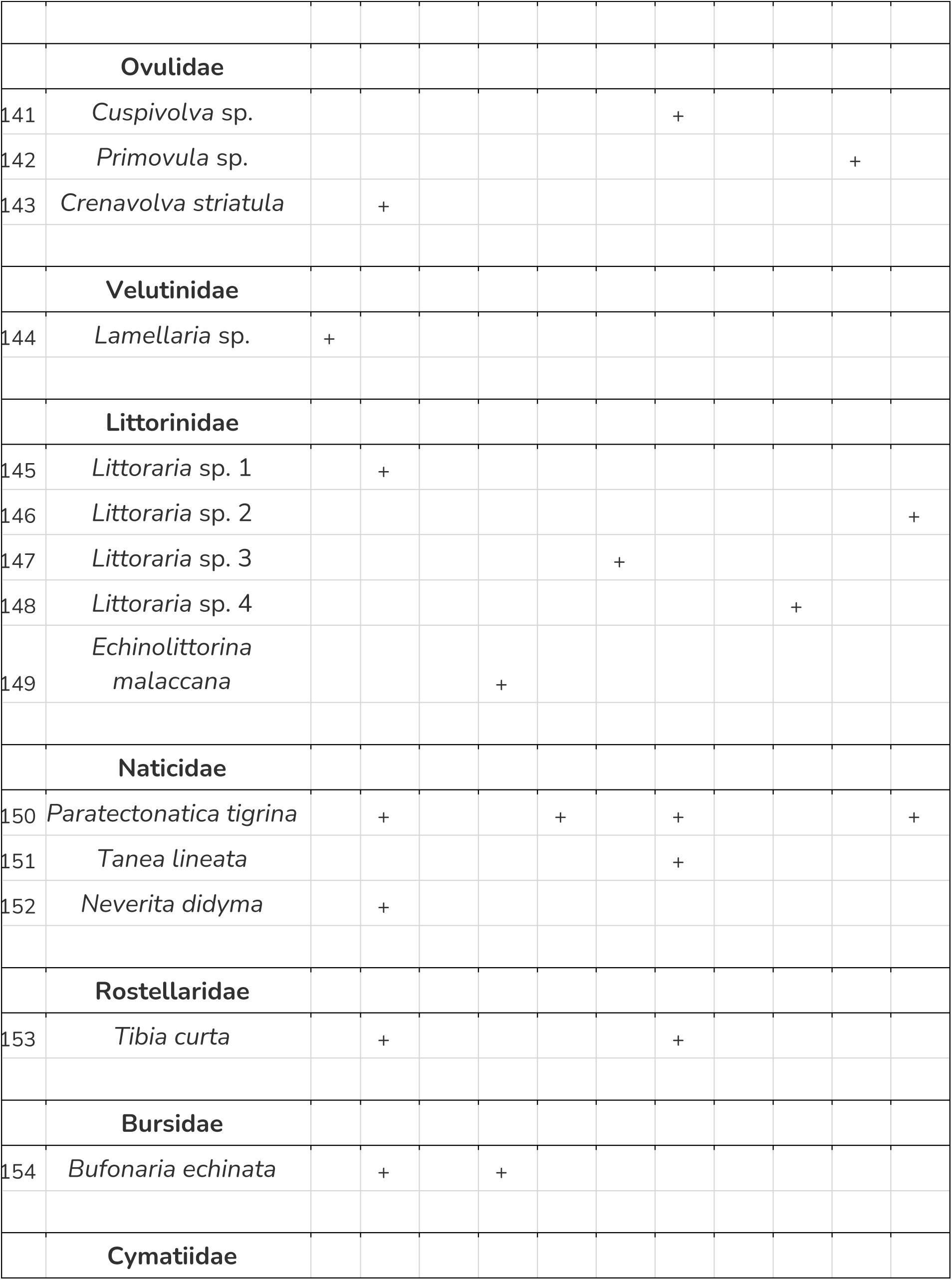
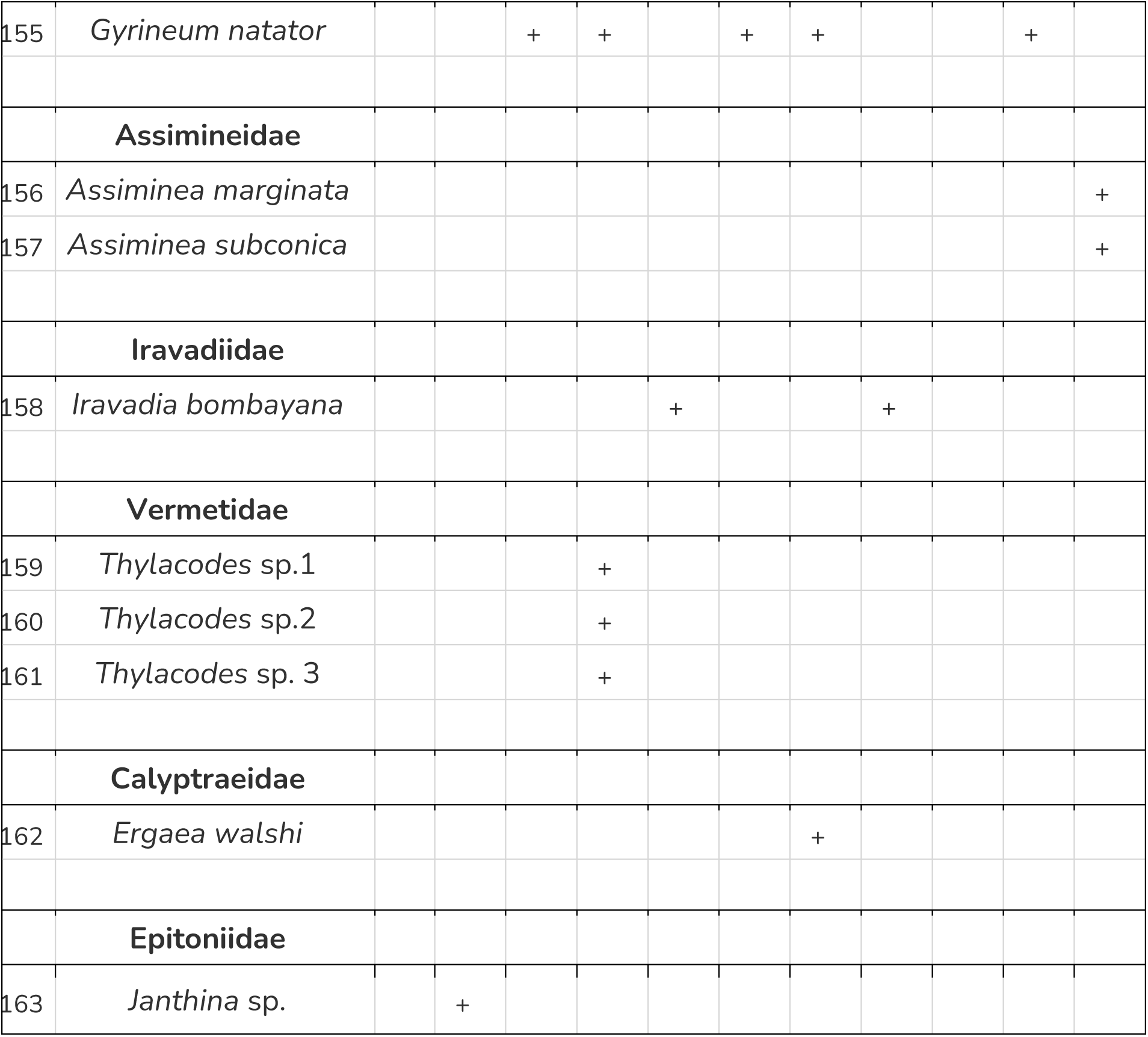
Zonewise distribution of Gastropods (Source: Authors).

**Table 5:** New Records of Gastropods from MMR (Source: Authors).

| New Records of Species from MMR |  |
| --- | --- |
| Family | Species |
| Aplysiidae | 1) *Aplysia* cf. *rudmani* |
| Haminoeidae | 2) *Smaragdinella sieboldi* |
| Onchidiidae | 3) *Paromoionchis penangensis* |
|  | 4) *Platevindex tigrinus* |
| Facelinidae | 5) *Herviella yatsui* |
|  | 6) *Cratena* sp.1 |
|  | 7) *Cratena* sp.2 |
| Bornellidae | 8) *Bornella stellifera* |
| Cuthonidae | 9) *Cuthona annandalei* |
| Tritoniidae | 10) *Marionia pambanensis* |
| Janolidae | 11) *Janolus mirabilis* |
| Chromodorididae | 12) *Hypselodoris zebrina* |
| Discodorididae | 13) *Atagema tristis* |
|  | 14) *Atagema* sp. |
|  | 15) *Carminodoris* sp.1 |
|  | 16) *Carminodoris* sp.2 |
| Dorididae | 17) *Doris immonda* |
|  | 18) *Sebadoris fragilis* |
| Goniodorididae | 19) *Goniodoris modesta* |
|  | 20) *Murphydoris puncticulata* |
|  | 21) *Trapania japonica* |
| Dendrodorididae | 22) *Dendrodoris fumata* |
|  | 23) *Doriopsilla miniata* |
| Polyceridae | 24) *Plocamopherus ceylonicus* |
|  | 25) *Thecacera pennigera* |
| Muricidae | 26) *Chicoreus torrefactus* |
| Terebridae | 27) *Duplicaria spectabilis* |
| Cypraeidae | 28) *Naria Turdus* |
| Ovulidae | 29) *Crenovolva striatula* |
| **New Records of Genera from MMR** |  |
| Limapontiidae | 1) *Ercolania* sp.1 |
|  | 2) *Ercolania* sp.2 |
|  | 3) *Ercolania* sp.3 |
| Costasiellidae | 4) *Costasiella* sp. |
| Pleurobranchidae | 5) *Berthella* sp. |
| Facelinidae | 6) *Phidiana* sp. |
|  | 7) *Noumeaella* sp. |
| Flabellinidae | 8) *Coryphellina* sp.1 |
|  | 9) *Coryphellina* sp.2 |
| Dotidae | 10) *Doto* sp.1 |
|  | 11) *Doto* sp.2 |
| Eubbranchidae | 12) *Eubbranchus* sp. |
| Trinchesiidae | 13) *Tenellia* sp.1 |
|  | 14) *Tenellia* sp.2 |
| Actinocyclusidae | 15) *Hallaxa* sp.1 |
|  | 16) *Hallaxa* sp.2 |
|  | 17) *Actinocyclus* sp. |
| Goniadorididae | 18) *Ceratodoris* sp.1 |
|  | 19) *Ceratodoris* sp.2 |
|  | 20) *Ceratodoris* sp.3 |
| Onchidorididae | 21) *Atalodoris* sp. |
| Dendrorididae | 22) *Doriopsilla* sp.1 |
|  | 23) *Doriopsilla* sp.2 |
| Dorididae | 24) *Doris* sp. |
| Polyceridae | 25) *Polycera* sp. |
|  | 26) *Thecacera* sp.1 |
|  | 27) *Thecacera* sp.2 |
| Ovulidae | 28) *Cuspidolva* sp. |
|  | 29) *Primovula* sp. |
| Velutinidae | 30) *Lamellaria* sp. |
| Vermetidae | 31) *Thylacodes* sp.1 |
|  | 32) *Thylacodes* sp.2 |
|  | 33) *Thylacodes* sp.3 |
| Muricidae | 34) *Reishia* cf. *bitubercularis* |

| New Records of Family from MMR |  |
| --- | --- |
| Limapontiidae | 1) Unidentified |

The subclass Heterobranchia is the most diverse group represented by 79 species, and subclass Patellogastropoda being the least diverse with three species. Family Muricidae has the highest representation with 14 records. Five species from the order Nudibranchia were rediscovered 78 years after their original description, while one species each from Neogastropoda, *Lataxiena bombayana,* and Siphonariida, *Siphonaria bassienesis* were recorded from their type locality after 131 and 31 years respectively (Table 6). Table 3 presents shelled gastropods categorised by observation type: live individuals or empty shells.

**Table 6:** Rediscoveries of species originally described from Mumbai. (Source: Authors).

| Family | Species |
| --- | --- |
| Chromodorididae | 1) *Glossodoris poecila* |
|  | 2) *Glossodoris semeion* |
|  | 3) *Goniobranchus bombayanus* |
|  | 4) *Goniobranchus trimarginatus* |
|  | 5) *Goniodoris kolabana* |
| Muricidae | 6) *Lataxiena bombayana* |
| Siphonariidae | 7) *Siphonaria basseinensis* |

### Family: Aplysiidae

1. *Aplysia* cf. *rudmani* (Bebbington, 1974) Plate 1

Original name: *Aplysia rudmani*

Description: Dark greenish brown in colour, with mottling of black, pale white and brown and bears opaque white spots crowded at some places as well as distantly placed. The parapodia is flap-like which helps them to swim and bears white crowded spots at its edge.

2. *Bursatella leachii* (Blainville, 1817) Plate 1

Original name: *Bursatella leachii*

Description: The animal is dark brown in colour and vibrant blue ocelli scattered on the whole body. Small black patches may be present. The body is covered with several elongated papillae that are branching from the sides as well as at the tip.

### Family: Haminoeidae

3. *Bakawan rotundata* (A. Adams, 1850) Plate 1

Original name: *Bulla (Haminea) rotundata*

Description: The animal is reddish orange and speckled with indistinct microscopic black spots. Two black eye spots are clearly visible. The shell is rounded - oval and translucent, having a whitish tinge. Very fine longitudinal striations can be seen on the shell. A greyish green morph is also found in the region. The green morph has also been observed mating with the red one.

4. *Haloa pemphis* (R. A. Philippi, 1847) Plate 1

Original name: *Bulla pemphis*

Description: The general body colour is translucent and pale cream in colour. White spots are scattered all over. The anterior side shows mottling of brown and orange colours. The shell is transparent and shows a reddish white tinge. Two black eye spots can be seen at the anterior end of the head.

5. *Haloa wallisii* (Gray, 1825) Plate 1

Original name: *Bulla wallisii*

Description: The animal is translucent pale white in colour with brownish black mottlings. The body is speckled with white, black, and orange spots of varying sizes. The shell is narrow, oval, and transparent with fine longitudinal lines. Eggs are laid in a super coiled thread like manner stuffed inside a short and transparent cylindrical tube.

6. *Smaragdinella sieboldi* (A. Adams, 1864) Plate 1

Original name: *Smaragdinella sieboldi*

Description: The base colour of the animal is translucent pale white, with light and dark brown and green mottlings present on the body. Patches of closely placed opaque white spots are seen on the body. The shell is oblong, oval, and transparent with fine longitudinal striations. It lays its eggs in a super coiled, spiral and inside an arched and transparent cylindrical tube.

### Family: Cylichnidae

7. *Cylichna cylindracea* (Pennant, 1777) Plate 11

Original name: *Bulla cylindracea*

Description: It is cylindrical in shape, rounded at both ends. Spire is turned inside like a scroll. Shell is translucent white in colour with extremely fine spiral lines.

### Family: Ellobiidae

**Subfamily: Pythiinae**

8. *Cassidula aurisfelis* (Brugiere, 1789) Plate 11

Original name: *Bulimus aurisfelis*

Description: The shell is ovate, maroonish brown in colour. The spire is elevated, plano-convex whorls. The whorls may show a cream band near the suture. The edge is thickened and flattens on the ventral side. The aperture is irregular and elongated. The outer lip also has an irregular crenation and inner lip bears three pointed teeth placed distantly from each other.

9. *Cassidula nucleus* (Gmelin, 1791) Plate 11

Original name: *Helix nucleus*

Description: The shell characteristics are like that of *C. aurisfelis,* the body whorl shows several cream colour bands of varying thickness.

10. *Ellobium* sp. (Roding, 1798) Plate 11

Original name: N/A

Description: The shell is fusiform, off white in colour. Sutures are simple, elevated spire, whorls are slightly convex. The aperture is elongated drop-like. The inner lip is thick and smooth, the outer lip is thick and rounded and shows a tooth towards the upper end.

**Subfamily: Melampodinae**

11. *Melampus sincaporensis* (L. Pfeiffer, 1855) Plate 11

Original name: *Melampus sincaporensis*

Description: The shell is oval and fusiform in shape and bears longitudinal striations. The body whorl is 2/3^rd^ of the spire. The whorls are rounded. It is pale yellow in colour and a transverse thick brown band is present near the sutures. The body whorl shows four transverse brown bands of varying thickness. The aperture is narrow and sub vertical. The inner lip shows three parietal folds. The outer lip shows 3-4 denticulations on the inner side.

### Family: Onchidiidae

12. *Paromoionchis penangensis* (Dayrat & Goulding, 2109) Plate 1

Original name: *Paromoionchis penangensis*

Description: The dorsal surface will always be covered in mud and colour is light grey to dark grey. The dorsal surface appears leathery and warty with some large tubercles in between. The pedicels are pale orange in colour.

13. *Peronia verruculata* (Cuvier, 1830) Plate 1

Original name: *Onchidium verruculatum*

Description: The general body colour is dark grey. The mantle is quite thick, rough, and completely covered with conical tubercular elevations of irregular sizes placed randomly in a compact manner.

14. *Platevindex tigrinus* (Stolickza, 1869) Plate 1

Original name: *Onchidium tigrinum*

Description: The dorsum appears thick and leathery and is light pale brown in colour. Large black spots are scattered randomly on the mantle. Small, rounded tubercles cover the whole dorsal surface with some larger ones in between.

### Family: Plakobranchidae

15. *Elysia bangtawaensis* (Swennen, 1998) Plate 2

Original name: *Elysia bangtawaensis*

Description: The body is broad in front and tapering towards the end. General body colour is dark green which fades to light green towards the head. Opaque white and light green small spots of varying sizes are scattered all over the body including parapodia. The parapodia is thin, large and has small orange spots placed distantly at the edge. The head also shows the same specifications except the orange spots.

16. *Elysia* cf. *hirasei* (Baba, 1955) Plate 2

Original name: *Elysia hirasei*

Description: The body is translucent light green in colour and has a leaflike shape, rear end is tapering, and conical and front end appears to have head notched in the centre. The head bears a transverse black band in the front which divides into two longitudinal lines and median to these two lines is a pinkish red patch, which seems to be present only in the individuals found in the geographical region. Two longitudinal lines also run from the front end of the lateral sides of the head up to the parapodia. There is a white circular patch with several distinct, opaque, and small white spots present just behind the head. The ramifying digestive glands resembling veins on a leaf are spread all over the body and parapodia. The parapodia has opaque distinct and irregularly shaped white spots placed almost equidistantly on the edge. In some individuals, the parapodia is seen speckled with small and opaque white spots. The edge also shows the presence of a brownish cream band made up of small spots densely packed at the edge. The head is pale and translucent green in colour with light pink coloured spots that are very closely placed almost smudging into each other.

17. *Elysia tomentosa* (K. R. Jensen, 1997) Plate 2

Original name: *Elysia tomentosa*

Description: The body is uniformly bright green in colour having a pair of large parapodia of irregular margin. The parapodial margin is lined with black on either side and slight creamish pigmentation in the centre. The body surface bears numerous pale white papillae and small, opaque black spots. The egg ribbon is laid in a spiral manner, and eggs are yellow in colour arranged in tight coils.

18. *Elysia* sp. (Risso, 1818) Plate 2

Original name: N/A

Description: The body shape is the same as *E. hirasei.* It is uniformly light green in colour and speckled with small white spots and some larger orange spots. The parapodial edges show larger white spots that are distantly placed. The rhinophores show densely packed tiny light orange spots at the tip. Eye spots can be seen at the base of the rhinophores.

### Family: Limapontiidae

19. *Ercolania* sp.1 (Trinchese, 1872) Plate 2

Original name: N/A

Description: The body is translucent light brown in colour with minute white and dark brown specks all over. The rhinophores are simple and cylindrical. Rhinophores bear brown lines that extend up to their head. Cerata are flat and leaf-like having the same colouration as the body. Digestive glands can be seen through the translucent membrane.

20. *Ercolania* sp.2 (Trinchese, 1872) Plate 2

Original name: N/A

Description: It is translucent with dark grey to black pigmentation. The head shows a black band running from in between the rhinophores and black long markings in front of the head. The eye spots can be seen laterally just behind the rhinophores on a translucent background. The rhinophores are translucent with white markings at its base and tip, and black in the middle. The cerata are flat, leaf-like and black in colour with scattered white spots and white tips.

21. *Ercolania* sp.3 (Trinchese, 1872) Plate 2

Original name: N/A

Description: It is translucent white in colour with black pigmentation. The rhinophores are simple, cylindrical and have black pigmentation, white at the tips. Cerata are narrow, have black pigmentation all over and specks of white in the middle and at the tip.

22. Unidentified Plate 2

Original name: Limapontiidae (Gray, 1847)

Description: The body is translucent, pale cream in colour and bears smudged markings of pale green colour all over. The cerata are long, narrow and cylindrical having similar colouration as the body. The rhinophores are simple, cylindrical and are speckled with white and pale green spots.

### Family: Costasiellidae

23. *Costasiella* sp. (Pruvot-fol, 1951) Plate 3

Original name: N/A

Description: The body is translucent, pale greyish cream in colour and is speckled with white and black colour spots. The rhinophores have a similar colour to the body with a dark cream band seen inside it. The cerata are bulbous, slightly arched and heavily speckled, crowded white specks at the tips.

### Family: Creseidae

24. *Creseis sp.* (Rang, 1828) Plate 10

Original name: N/A

Description: It is a transparent, needle-like conical shell, a few mm in size.

### Family: Siphonariidae

25. *Siphonaria basseinensis* (Melvill, 1893) Plate 13

Original name: *Siphonaria basseinensis*

Description: The shell is sufficiently elevated and conical. It is oval in shape. It is pale cream to white in colour with biradiate brown stripes.

### Family: Pleurobranchidae

26. *Berthella* sp. (Blainville, 1824) Plate 3

Original name: N/A

Description: The body is round and bulbous. Uniformly translucent white in colour with small white opaque and irregular spots. The texture of the body resembles that of a sponge.

### Family: Facelinidae

27. *Cratena lineata* (Eliot, 1905) Plate 3

Original name: *Hervia lineata*

Description: The body is translucent pale white in colour. Distinct white longitudinal lines are present on the notum and cerata. The cerata are dull and light orange coloured and tipped with white. There is a bright orange colour patch between the rhinophores and tentacles. The upper half of the rhinophores are pale white while the lower half shows slight orange colouration.

28. *Cratena pawarshindeorum* (Bharate, Padula, Apte & Shimpi, 2020) Plate 3

Original name: *Cratena pawarshindeorum*

Description: Translucent pale white body with cerata translucent and dark reddish digestive glands with a white tip can be seen in the centre. Rhinophores are translucent pale white at the base and at the tip, mid region is bright orange coloured. Oral tentacles are of the same colour as the body with little white pigmentation near the tip. Two large bright orange patches are present between the oral tentacles and rhinophores, almost smudging into each other. The egg ribbon appears white in colour and is wrapped in tight coils on the branches of the hydroid it feeds on.

29. *Cratena* sp.1 (Bergh, 1864) - Plate 3

Original name: N/A

Description: The morphological characters of this *Cratena* sp. are very similar in appearance to *C. pawarshindeorum*. The differences occur in the colouration of the rhinophores, which are light orange in colour and the orange patches present in between the oral tentacles and rhinophores are placed apart at a distance from each other dorsally on top of the head as compared to its former counterpart.

30. *Cratena* sp. 2 ((Bergh, 1864) Plate 3

Original name: N/A

Description: The body is translucent white in colour. The rhinophores and oral tentacles are translucent at the base and white towards the tip. The cerata are translucent with white irregular marks scattered throughout and dark red on the inside. The patches between the oral tentacles and rhinophores are small and faded orange in colour.

31. *Herviella yatsui* (Baba, 1930) Plate 4

Original name: *Cratena yatsui*

Description: The colour of the body is pale cream with brownish grey patch running throughout the notum. The base colour of cerata is also pale cream with brownish grey pigmentation and tipped with white. The oral tentacles and rhinophores show three bands of colouration - basal pale cream, mid dark brownish grey and tip is white.

32. *Phidiana* sp. (Gray, 1850) Plate 4

Original name: N/A

Description: The body is translucent white in colour. The oral tentacles bear a white tip followed by orange colouration that continues a line in front of the head. The rhinophores also show a similar pattern of colouration. The cerata are cylindrical with pointed orange tips and a white line running in the centre with black digestive glands visible inside.

33. *Noumeaella* sp. (Risbec, 1937) Plate 4

Original name: N/A

Description: It is dark grey to black in colour with pale white blotches all over the body. The white blotches are crowded in the oral tentacles and at tips of cerata. The cerata are cylindrical with pointed and curved tips. The rhinophores have dark grey to black pigmentation with white markings at the tips.

### Family: Flabellinidae

34. *Coryphellina* sp. 1 (O’Donoghue, 1929) Plate 4

Original name: N/A

Description: The body is translucent with a broken vibrant bluish-purple line running in the centre dorsally. The oral tentacles bear a white tip, bluish purple in the centre and translucent at the base. The rhinophores are translucent at the base and bluish purple at the tip. The cerata show the same pattern of colouration as that of oral tentacles with distinct, orange-coloured digestive glands inside. The tip of the tail too shows bluish purple pigmentation.

35. *Coryphellina* sp. 2 (O’Donoghue, 1929) Plate 4

Original name: N/A

Description: The body is translucent with a small sky blue-ish iridescent line present in between the base of the rhinophores. The rhinophores are translucent at the base and have a light purplish-blue colouration at the tip. The oral tentacles are translucent at the base, sky blue-ish iridescence in the middle and white at the tip. The cerata too are white at the tip, sky blue-ish iridescent sub-apically and translucent at the remaining length with light orange digestive glands visible in the centre.

### Family: Bornellidae

36. *Bornella stellifera* (A. Adams & Reeve, 1848) Plate 4

Original name: *Dendronotus stellifer*

Description: The body is pale orangish brown in colour and bears opaque and large white distinct patches that appear as cracks on a surface. The cracks in between the white patches appear orange in colour. Lateral appendages and rhinophore sheaths bear a white tip, followed by orange colour and a darker orange band. The rhinophores are present on a long stalk with a ring of long pointed papillae. The oral tentacles are modified into star-like sensory organs.

### Family: Dotidae

37. *Doto* sp.1 (Oken, 1815) Plate 4

Original name: N/A

Description: Translucent white coloured body rounded anterior and tapering posterior foot corners. Brown coloured branching patterns can be seen all over the body. The rhinophoral sheaths are translucent white in colour may or may not show the brown branching pattern and shows presence of small distinct small white spots. The rhinophores are brown in colour bearing white tip. Cerata are irregular with several conical tubercles, transparent sheath. The egg mass is laid in the form of ribbons that are compactly coiled on the hydroid branches and is pale yellow in colour. Individuals with dark brown pigmentation and a non-pigmented body were also observed.

38. *Doto* sp.2 (Oken, 1815) Plate 4

Original name: N/A

Description: The body is translucent, cream in colour, speckled with irregular spots of pale yellow, white and black colour towards the head. Cerata brown in colour, have conical tubercles with white tips. Rhinophores are translucent and speckled with white, more at the tip. Rhinophoral sheaths are also translucent.

### Family: Eubranchidae

39. *Eubranchus* sp. (Forbes, 1838) Plate 5

Original name: N/A

Description: General body is translucent white with distinct black spots scattered all over the body except the foot. The foot’s tentacular processes, oral tentacles, ceratal tips, and rhinophores have a subapical orange band, and their tips appear to have shining iridescent pigmentation. The cerata are arranged laterally in vertical rows and each cerata has three circlets of conical tubercles. Yellowish orange stem of the digestive gland is visible inside the cerata.

### Family: Cuthonidae

40. *Cuthona annandalei* (Eliot, 1910) Plate 5

Original name: *Cuthona annandalei*

Description: The body is translucent off white to white in colour. The foot is rounded and disc-like and produces two tentacular processes anteriorly. The body surface shows white shining patches present all over and light grey patches can also be observed in some individuals on rhinophores or on the back of head. The rhinophores and foot processes are translucent, covered with shining white pigmentation. The cerata are arranged on either side of the body with a gap seen after the first two pairs. Ceratal tips become narrower towards, giving the appearance of bulging sub-apically, the degree of which seems to vary from individual to individual. Light to dark grey digestive glands is visible through the translucent membrane of cerata. Their tips may show bands of shining pigmentation or may be completely smudged with it.

### Family: Trinchesiidae

41. *Tenellia* sp.1 (Lhering, 1879) Plate 5

Original name: N/A

Description: The body and cerata are translucent and bears shining iridescent white to peach pigmentation, which extends as a band in between the rhinophores and oral tentacles. The rhinophores and oral tentacles have orange pigmentation that fades towards the tip. The cerata are cylindrical and slightly bulbous.

42. *Tenellia* sp.2 (A. Costa, 1866) - Plate 5

Original name: N/A

Description: The body is translucent, viscera being clearly visible in yellow. Rhinophores are simple and translucent. The cerata are cylindrical, with a subapical bulging. It has white and orangish brown irregular markings. The branching digestive glands, brown in colour, are distinctly seen in the centre of the body.

### Family: Tritoniidae

43. *Marionia pambanensis* (O’ Donoghue, 1932) Plate 5

Original name: *Marionia pambanensis*

Description: The body is oval and tapering towards the hind end. The colour of the body is translucent cream with internal organs visible. The oral veil is present at the anterior end, in front of the rhinophores and is bifurcated. The dorsal surface and lateral sides are covered by small conical pustules. The mantle and the oral veil bear a symmetrical and reticulated pattern of dark brown and orange, brown colour, running in between the pustules.

### Family: Janolidae

44. *Janolus mirabilis* (Baba & Abe, 1970) Plate 6

Original name: *Janolus mirabilis*

Description: The colour of the body, cerata and rhinophores is pale grey with irregular marks of brown and pale white. Rhinophores are closely situated to each other. The cerata have pointed tips and several conical tubercles and similar extensions on the anterior side. The cerata are arranged in two groups, one is placed anteriorly and other posteriorly.

### Family: Actinocyclidae

45. *Hallaxa* sp. 1 (Eliot, 1909) Plate 6

Original name: N/A

Description: Shape of the body is elongated and ovoid. Body colour is translucent pale white to having brownish tinge with internal organs visible. White spots are scattered unevenly on the notum. The rhinophores are translucent pale white with brown pigments at the top. The branchial plume is smoky dark grey in colour. The egg mass is creamish white and laid in a spiral and ruffled ribbon form.

46. *Hallaxa* sp. 2 (Eliot, 1909) Plate 6

Original name: N/A

Description: The body is elongated and oval in shape. It is translucent with internal organs visible. The entire notum is speckled with faint brownish grey spots along with irregular and raised white spots placed distantly. The branchial plume is also similar in appearance with white spots at extreme ends. The tail has a pointed tip, is translucent with brownish grey tinge and absentia of any white spots.

47. *Actinocyclus* sp. (C.G. Ehrenberg, 1837) Plate 6

Original name: N/A

Description: The body is oval in shape, translucent and heavily pigmented with cream colour. Notum is decorated with round, elevated tubercles, brown in colour which are depressed in the centre with black pigments. The tubercles are more in number on the sides and surrounding the gills. The gills are reddish pink in colour.

### Family: Chromodorididae

**Subfamily: Chromodoridinae**

48. *Glossodoris poecila* (Winckworth, 1946) Plate 6

Original name: *Glossodoris poecila*

Description: The base colour of the body is pale white to pale blue in colour with irregular yellow spots of varying sizes present all over the notum. The notum also bears a network of red lines and patches which seem to differ in different individuals. The mantle’s edge has yellow and light purple colour spots with outlines of bright blue, which becomes red on joining the central network of red lines and patches. The gills are pale white with reddish orange outlines. The rhinophores also are pale white in colour with orange pigmentation, faded from the base and darkened towards the tip. The tail also has a similar pattern of colouration.

49. *Glossodoris semeion* (Winckworth, 1946) Plate 6

Original name: *Glossodoris semeion*

Description: The notum is pale white in colour with irregular purple spots of varying sizes present all over the body. The gills and rhinophores are white in colour with orangish yellow pigmentation towards their tips. The edge of the notum has a sub marginal yellow band.

50. *Goniobranchus bombayanus* (Winckworth, 1946) Plate 6

Original name: *Glossodoris bombayana*

Description: General body colour is white. Mantle margin has a deep orange colour with a row of dark purple spots. Small white spots are densely populated and concentrated in the centre of the dorsal surface. Large dark purple spots are scattered irregularly. The foot is short and white in colour. The foot, rhinophores and gills are translucent and pale white, speckled with silver spots.

51. *Goniobranchus trimarginatus* (Winckworth, 1946) Plate 7

Original name: *Glossodoris trimarginata*

Description: Body is translucent, pale white to creamish in colour. The notum is decorated with red spots of varying sizes and closely placed. The edge of the mantle is trimarginated. The outermost band is white in colour with a slight blue tinge, middle band is deep red and innermost band is orange red in colour. It also shows the presence of highly branched mantle glands which are white in colour. The gills and rhinophores are pale white and translucent with white edges.

**Subfamily: Miamirinae**

52. *Hypselodoris zebrina* (Alder & Hancock, 1864) Plate 7

Original name: *Chromodoris zebrina*

Description: Body is elongated, rounded in front, and tapering behind. The general body colour is translucent pale white. The notum has yellowish - orange spots and a red coloured line running in the centre from the front end to 1/3rd of the body. The edges of the mantle skirt have spots of similar colour to that of the body along with purplish blue spots. The gills and rhinophores have orange pigmentation.

### Family: Discodorididae

53. *Atagema osseosa* (Kelaart, 1858) Plate 7

Original name: *Doris osseosa*

Description: The shape of the body is elliptic and oblong. The general body colour is uniformly grey or creamish white. The notum is covered with compound stellate clusters of several caryophyllida. The rhinophores and gills are of the same colour as the body, in two of the individuals observed, the rhinophores were lightly pigmented with brown on the outlines.

54. *Atagema tristis* (Alder & Hancock, 1864) Plate 7

Original name: *Doris tristis*

Description: The body is slightly raised and is ovate and oblong in shape. The body is pale to bright cream or pale grey in colour. The notum has angular ridges which show dark grey pigmentation at the sides filled with large tubercular elevations. Few minute tubercles with light brown pigmentation are also present towards the mantle edge. The rhinophores are pigmented with grey at the tip and pale at the base. In one of the observations, a brown indistinct line is seen in the centre of the notum, behind the rhinophores that breaks in the middle and then reaches up to the gill branches.

55. *Atagema* sp. (Gray, 1850) Plate 7

Original name: N/A

Description: The shape and features of the body are similar to that of *A. osseosa.* The colour of the body, gills and rhinophores is smoky black.

56. *Carminodoris* cf. *grandiflora* (Pease, 1860) Plate 7

Original name: N/A

Description: The body shape is oval and oblong, slightly raised in the centre having a slight orangish tinge. The basal body is creamish to grey in colour with dark grey wide striations from edge of the mantle to the central part of the dorsal surface. The notum is covered by prominent rounded tubercles with varying sizes and becomes smaller in size towards the mantle edge and most of them have white circling ring at the base. The rhinophores have brownish black pigmentation and are tipped with white. The gills have a deep grey colour with white outline. The egg masses are in a form of flat, ruffled ribbon laid in a spiral manner.

57. *Carminodoris* sp. 1 (Bergh, 1889) Plate 7

Original name: N/A

Description: Body is oval and pale brown in colour with random black patches. Notum is filled with rounded tubercles of the same colour as the body with a bit darker pigmentation at their tips and a white ring at the base. The gill branches are six or seven in number and brown in colour with black pigmentation at the border. Rhinophores have pale brown colour and darker pigmentation of brown at the tip.

58. *Carminodoris* sp. 2 (Bergh, 1889) Plate 7

Original name: N/A

Description: The shape of the body is similar to its counterparts. The colour of the notum is translucent grey with teal green pigmentation. It also has grey band-like patches scattered all over. The surface is completely covered with rounded tubercles that are shortest towards the edge, increasing in size towards the centre, and several longest ones concentrated in the centre. The tubercles also show teal green to white coloured rings at their base. The rhinophores and the gill branches are translucent grey towards the base, getting darker grey towards the tip and bear white pigmentation and outline.

### Family: Dorididae

59. *Doris immonda* (Risbec, 1928) Plate 8

Original name: *Platydoris immonda*

Description: The mantle is pale yellowish cream in colour and bears several rounded tubercles of varying sizes with black pigments. The centre of the mantle has a slight long and rounded elevation, the middle portion of which is pale white in colour. The rhinophores have black pigmentation at their edges.

60. *Doris* sp. (Linnaeus, 1758) Plate 8

Original name: N/A

Description: The mantle, rhinophores and gills are lemon yellow in colour. The mantle appears textured with short and pointed tubercles present all over the surface of the notum.

61. *Sebadoris fragilis* (Alder & Hancock, 1864) Plate 8

Original name: *Doris fragilis*

Description: The base colour of the body is cream which gets paler at the margin and profusely spotted with brown spots. The centre of the notum is marked by a brown band and has white pigmentation inner to it. The notum also bears randomly placed brown to black blotches. Gills are dark brown in colour. The rhinophores are also brown in colour with a darkened outline and have slight white pigmentation towards the tip.

### Family: Goniodorididae

62. *Pelagella kolabana* (Winckworth, 1946) Plate 8

Original name: *Goniodoris kolabana*

Description: The body is translucent, pale cream to pale white in colour and is minutely speckled with brown spots, hence giving a general appearance of pale brown colour. The inner organs are visible. The edge of the mantle is rounded and rolled downwards with small conical tubercles seen on the lateral side and under the mantle fold. It meets behind the gills and is continued into a dorsal median crest. The rhinophores are pale and pigmented with brown at the base and white or cream at the tip. Gills have scarce to deep pigmentation of brown, base colour being pale white or cream. The head bears two lateral tentacles.

63. *Pelagella modesta* (Alder & Hancock, 1864) - Plate 8

Original name: *Goniodoris modesta*

Description: The body is ovoid in shape. Notum, gills and rhinophores appear as a mottling of white and brown colours. The mantle’s edge has large yellow and brown spots. The gill branches are pale white in colour with brown pigmentation all over. The rhinophores bear the same pattern as the notum.

64. *Murphydoris puncticulata* (Paz-Sedano, Smirnoff, Candás, Gosliner & Pola, 2022) Plate 8

Original name: *Murphydoris puncticulata*

Description: Pale white and translucent body with densely packed cream patches present all over the notum, rhinophores and gills. Dark brown, irregularly shaped, small spots are present behind the rhinophores forming a triangular shape and few are found scattered on the mantle margin and notum.

65. *Trapania japonica* (Baba, 1935) Plate 8

Original name: *Drepania japonica*

Description: The general body colour is pale white and translucent and internal organs are visible. Dark brown patches of irregular shape are seen on the body everywhere including the rhinophores. The gill branches too have dark brown colouration appearing like a venation. Extra-branchial and extra-rhinophoral appendages are yellowish orange in colour.

66. Ceratodoris sp.1 (Menke, 1830) Plate 8

Original name: N/A

Description: The mantle is translucent white in colour with finger-like extensions. The mantle, rhinophores and gills are speckled with yellow and reddish spots of varying sizes. The egg mass is laid in a tight and flat spiral, and it is white in colour.

67. *Ceratodoris* sp. 2 (Menke, 1830) Plate 8

Original name: N/A

Description: The body is translucent, orange in colour with finger-like processes present at the edges of the mantle that are white in colour. Irregularly shaped spots of dark brown colour are present all over the body, mantle extensions and gills. The rhinophores and gills are orange in colour with brown specks at their tips. One finger-like process is also present just before the gills.

68. *Ceratodoris* sp.3 Plate 8

Original name: N/A

Description: This *Ceratodoris sp.* has similar morphological characteristics as the one mentioned above, except it is entirely translucent white in colour with absence of irregular brown spots.

### Family: Onchidorididae

69. *Atalodoris* sp. (Iredale & Donoghue, 1923) Plate 9

Original name: N/A

Description: It is oval in shape with a translucent white body. The body texture resembles exactly like the bryozoan it feeds on. The rhinophores have a slight dense light orange to brown pigmentation and gills are translucent white in colour.

### Family: Dendrodorididae

70. *Dendrodoris fumata* (Ruppell & Leuckart, 1830) Plate 9

Original name: *Doris fumata*

Description: The body is ovoid, elongated and rounded at both ends. The colour can vary from orange, pink, smoky grey, brown, or black. Black or grey blotches are present on the dorsal surface. Rhinophores and gills have a similar colour like that of the notum. An individual with bushy and smoky grey coloured gills has also been observed.

71. *Doriopsilla miniata* (Alder & Hancock, 1864) Plate 9

Original name: *Doridopsis miniata*

Description: It has a highly decorated notum with deep reddish orange colour which appears to fade towards the mantle’s edge, in some specimens the mantle edge has creamish white colour. Tubercles are rounded and pale orange in colour are present all over the dorsal surface. Distinct small white spots can be seen more populated surrounding the tubercles and the mantle margin. Gills and rhinophores are pale and light orange in colour and in some are deeply pigmented. The colouration differs slightly from the specimens observed elsewhere. The eggs are laid in a spiral and flat ribbon form, yellow in colour.

72. *Doriopsilla* sp. (Bergh, 1880) Plate 9

Original name: N/A

Description: The colour of the mantle is orangish brown, which is lighter in the centre and darker towards the edge. It is completely covered with rounded tubercles of varying sizes. Some of the tubercles have a white ring at their base. The rhinophores and gills are orangish brown at the base, dark brown sub-apically and white pigmentation at their tips.

## Family: Polyceridae

**Subfamily: Triophinae**

73. *Plocamopherus ceylonicus* (Kelaart, 1858) Plate 9

Original name: *Polycera zeylanica*

Description: The general body colour is pale white and brown blotches form patterns on the dorsum. The body is speckled with small brown spots and large orangish yellow spots. The orange - yellow spots form a band at the anterior most margin of head. The head is veiled with a fimbriated membrane. The lateral sides of the dorsum possess three pairs of papillae that end in pink to brown rounded knobs and emit light when disturbed. The foot is translucent pale white in colour with bright orange spots.

**Subfamily: Polycerinae**

74. *Polycera* sp. (Cuvier, 1816) - Plate 9

Original name: N/A

Description: The body is rounded anteriorly and tapering posteriorly. It is pale white in colour with grey colouration, bearing yellow, black and orange irregular specks. Narrow, conical and yellow-coloured extensions are seen scattered all over the body. The gills and rhinophores have blackish grey pigmentation and have yellowish orange pigmentation at the tip.

75. *Thecacera pennigera* (Montagu, 1813) Plate 9

Original name: *Doris pennigera*

Description: The body is translucent and speckled with black and orange irregular spots or varying sizes, including the rhinophores and gills. Extra-rhinophoral and extra-branchial appendages have a white tip.

76. *Thecacera* sp.1 (J. Fleming, 1828) Plate 10

Original name: N/A

Description: The body, rhinophores and gills are translucent and dull coloured. They bear black spots densely placed almost covering all the surfaces, in between are yellow- and orange-coloured irregular spots.

77. *Thecacera* sp.2 (J. Fleming, 1828) Plate 10

Original name: N/A

Description: The body is translucent and white in colour and has long black patches present all over the body. Body is also speckled with orange and pale-yellow spots of different sizes. Similar pattern is observed on rhinophores and gills. Extra-rhinophoral appendages have a creamish tip, black towards the tip and an orange band in the centre. Extra-branchial appendages have a similar colour pattern as that of the body.

### Family: Neritidae

**Subfamily: Neritininae**

78. *Neripteron violaceum* (Gmelin, 1791) Plate 11

Original name: *Nerita violacea*

Description: The general colour of the shell is cream with an orangish tinge and black patches and wavy bands forming latitudinal patterns which vary from individual to individual. The shell is thick, solid, and globose. The spire is flat and strongly keeled posteriorly. The aperture is half-moon shaped and callous is thick and appears deep violet, maroon, light or dark brown coloured. The outer lip is thin and smooth. The inner lip appears crenated.

79. *Nerita albicilla* (Linnaeus, 1758) Plate 11

Original name: *Nerita albicilla*

Description: The shell is thick, solid, and globose. The spire is not elevated, the apex appears a bit depressed and strongly keeled posteriorly. The aperture is half-moon shaped. The columella is thick, creamy, and calloused. The outer lip is thin, sharp, crenulated on the inside and at the edge has light brown spots. The inner margin bears 4 weak teeth and several granulations. The shell’s colour is variegated by black and white.

80. *Nerita chamaeleon* (Linnaeus, 1758) Plate 11

Original name: *Nerita chamaeleon*

Description: The shell is thick, solid, and globose. The spire is not elevated, the apex is pointed. The whorls are convex and are sculptured with spiral ridges. Aperture is small, half-moon shaped and deep. The columella is creamy, thick and the outer margin is thin, sharp, and traversed by two to three longitudinal ridges and granulations on the outer and inner side. The outer margin has black spots or a thin black band. The inner lip is denticulated, with the first two superior ones being knob shaped and the inferior one is the strongest. The spiral ridges are wavy, and its colour can vary from greenish black, greyish, reddish orange with blotched oblique bands of a different colour.

81. *Nerita oryzarum* (Récluz, 1841) Plate 11

Original name: *Nerita oryzarum*

Description: The shell is thick, solid, and globose. The spire is flat. The whorls are convex and have spiral ridges. Aperture is half-moon shaped and deep. The outer lip’s inner margin is crenulated and the outer margin thin and sharp. The inner lip is denticulated with three pointed small teeth. The callous has few longitudinal ridges and granulations. The base colour of the shell is cream and bears irregular spiral greyish black or reddish orange bands.

82. *Clithon colombarium* (Recluz, 1846) Plate 11

Original name: *Nerita (Clithon) colombaria*

Description: The shell is comparatively smaller and thinner than *Nerita* sp. Shape is oval, the spire is compressed. Surface appears smooth and glossy. The colour and patterns of the shell highly vary. The whorls have been observed to have chipped surfaces or damaged surfaces. The aperture is half-moon shaped. Columella is slightly calloused and bears three fine teeth in the depression.

### Family: Chilodontoidae

83. *Euchelus asper* (Gmelin, 1791) Plate 12

Original name: *Trochus asper*

Description: The shell is quite thick, and the spire is elevated. The shell appears dark brownish grey in colour. The whorls are inflated and have a beaded appearance because of grainy spiral ridges. The aperture is round. The columella has a conspicuous tooth at the base. The dorsal ridges extend up to the margin of the outer lip thereby giving it a toothed appearance.

### Family: Fissurellidae

**Subfamily: Diodorinae**

84. *Diodora lima* (G. B. Sowerby II, 1862) Plate 12

Original name: *Fissurella lima*

Description: The shell margin is oval, conical, concave and inclined anteriorly. Prominent spiral and radial ridges seen.

85. *Diodora ruppellii* (G. B. Sowerby I, 1835) Plate 12

Original name: *Fissurella ruppellii*

Description: The shell is oval in shape and slightly elevated. Shell surface bears fine radial ribs and riblets. It is pale orange in colour with eight radial black bands. The aperture is elongated.

**Subfamily: Emarginulinae**

86. *Scutus unguis* (Linnaeus, 1758) Plate 12

Original name: *Patella unguis*

Description: The shell is flat, oblong, shield-like and reddish brown in colour. It is posteriorly inclined. The mantle extends out of the shell and almost covers the shell completely. The colour of the mantle is translucent white and black reticulated blotches all over. The muscular impression of the foot is horseshoe shaped.

### Family: Trochidae

**Subfamily: Trochinae**

87. *Trochus radiatus* (Gmelin, 1791) Plate 13

Original name: *Trochus radiatus*

Description: Shell is conical with the whorls straight, and its base is circular and flat. The whorls have three spirally arranged rows of tubercles, in which the top row has larger tubercles than the two below it. The base also has spirally arranged ridges. The base colour is cream with a pinkish or green tinge and wavy oblique bands of reddish-brown colour are present on it.

88. *Clanculus depictus* (A. Adams, 1855) Plate 13

Original name: *Clanculus depictus*

Description: Conical shell, with flat whorls and a slightly convex base. Smaller than and very similar to *T. radiatus*, hence often confused as its juvenile form. It is generally cream to light yellow in colour, sometimes uniformly grey, with black to brownish black stripes.

**Subfamily: Umboniinae**

89. *Umbonium vestiarium* (Linnaeus, 1758) Plate 13

Original name: *Trochus vestiarius*

Description: The shell is solid, disc shaped and compressed. The surface of the shell is smooth and glossy. The colour and patterns on the shell highly vary. The operculum is very small, thin, and circular.

### Family: Turbinidae

**Subfamily: Turbininae**

90. *Turbo intercostalis* (Menke, 1846) Plate 13

Original name: *Turbo intercostalis*

Description: The shell is thick and heavy. Few prominent and thick spiral ridges encircle the whorl. A denticle is formed by extension of the largest middle ridge on the margin of the outer lip. Umbilicus is deep and narrow. The shell generally appears uniform grey in colour or has a chequered pattern of black or dull green.

91. *Astralium semicostatum* (Keiner, 1850) Plate 13

Original name: *Trochus semicostatus*

Description: The shell is chalky to creamy white in colour. Conical in shape, elevated spire, whorls are concave with its upper and lower margins scalloped and slightly extended outside the edges. Base is circular and flat with numerous lirae. Aperture is semioval and white in colour. Outer and lip edges are smooth.

### Family: Nacellidae

92. *Cellana radiata* (Born, 1778) Plate 13

Original name: *Patella radiata*

Description: The shell is moderately conical and roundly ovate in shape. The base colour is pale cream, and rays have brownish black pigmentation.

93. *Cellana rota* (Gmelin, 1791) Plate 13

Original name: *Patella rota*

Description: The shell is moderately elevated and conical. The shape is more or less circular. The shell is slightly sculptured with fine riblets. It bears greyish black stripes in alternation with cream stripes with orange streaks on them.

94. Patelloida lentiginosa (Reeve, 1855) Plate 13

Original name: Patella lentiginosa

Description: The shell is moderately elevated and almost circular in shape. It bears greyish black stripes alternated with cream stripes. In some individuals the striped pattern may appear smudged. Diagnostic orange zig-zag streaks are clearly visible on the cream stripes.

### Family: Planaxidae

**Subfamily: Planaxinae**

95. *Planaxis sulcatus* (Born, 1778) Plate 14

Original name: *Buccinum sulcatum*

Description: Conical shell and is dark purplish or maroon in colour, white or cream colour spots arranged spirally on spiral striae. The aperture is wide, oval, and dark black in colour and sometimes can have dark purple grooves. The outer lip appears wavy because of the rounded ridges. Inner lip is pale white and shows light purple colouration.

### Family: Potamididae

96. *Pirenella cingulata* (Gmelin, 1791) Plate 14

Original name: *Murex cingulatus*

Description: Elongated and conical shell. The spire is structured with spirally arranged granulation. The shell is black in colour with transverse light brown lines present near the suture and several similar lines of varying thickness is seen on the body whorl. The aperture is half-moon shaped. The outer lip appears crenated and shows purple bands and the inner lip is smooth.

97. *Telescopium telescopium* (Linnaeus, 1758) Plate 14

Original name: *Trochus telescopium*

Description: The shell is thick and conical. Whorls are 14-16 in number, and sides are straight. It is brownish black in colour. The surface bears several flat and uneven spiral stripes. The aperture is small, twisted. Columella is canalled. The outer lip is thick and flares anteriorly over the siphonal canal.

### Family: Cerithiidae

98. *Clypeomorus batillariaeformis* (Habe & Kosuge, 1966) Plate 14

Original name: *Clypeomorus batillariaeformis*

Description: The shell is black in colour. Elongated, conical and the spire is elevated. The whorls are slightly convex and decorated with spirallly beaded ribs. Its egg ribbon is transparent with microscopic creamish-yellow eggs, long and loosely folded in a random manner, on a rock.

### Family: Thiaridae

Subfamily: Thiarinae

99. *Melanoides tuberculata* (O. F. Muller, 1774) Plate 14

Original name: *Nerita tuberculata*

Description: This species is known to have several shell colours and various patterns. Although it is a freshwater species, it is also reportedly found in almost every aquatic habitat provided the water is well oxygenated. The shell is conical, elongated and the spire is elevated. The whorls are rounded. The base colour is pale cream, ornamented with reddish orange flame like spots present longitudinally on each whorl. The surface of the shell is structured with spirally arranged ridges. The aperture is oval. The outer and inner lips are smooth.

### Family: Turritellidae

100. *Turritella duplicata* (Linnaeus, 1758) Plate 15

Original name: *Turbo duplicatus*

Description: The shell is elongate, conical and the spire is elevated and pointed. It is light brown in colour. The whorls are convex and bear 2 spirally placed sharp ridges in the centre. Sutures are slightly impressed and tight. The aperture is oval, both lips are smooth.

101. *Turritella* sp. (Lamarck, 1799) Plate 15

Original name: N/A

Description: the shell characteristics are similar as that of *T. duplicata.* Except the whorls show 3 spirally placed sharp ridges in their centre.

### Family: Babyloniidae

102. *Babylonia spirata* (Linnaeus. 1758) Plate 15

Original name: *Buccinum spiratum*

Description: The shell is smooth and heavy and the whorls are inflated. The spire is high, elongated with shoulders prominent and sutures are deep and channelled. The aperture is large and ovate with the outer lip sharp and strongly turned at the top and the interior is smooth and thick. The colour is pale white to cream with light brown blotches and spots arranged obliquely.

Family: Muricidae

103. Unidentified Plate 15

Original name: Muricidae (Rafinesque, 1815)

Description: The shell is spindle shaped and quite thick. It is reddish maroon in colour with cream colour bands of different thicknesses. The spire is elevated, and the whorls are flat and bear ridges with slightly raised tubercles. The sutures appear concave. The body whorl shows three rounded and slightly raised ridges. The outer lip is thick and bears 5-6 denticulations on its inside. The inner lip is thin and smooth.

**Subfamily: Muricinae**

104. *Chicoreus brunneus* (Link, 1807) Plate 15

Original name: *Purpura brunnea*

Description: The shell appears rhomboid in shape. Colour of the shell is burnt black. Elevated spire and body whorl is 2/3rd of the spire. The spiral cords are thick with very fine tubercles, varices are frilled and bear spines. The spines on the shoulder are elongate and foliate.

105. *Chicoreus torrefactus* (G. B. Sowerby II, 1841) Plate 15

Original name: *Murex torrefactus*

Description: The shell is spindle shaped, tapering at both ends. The surface bears numerous latitudinal reddish orange lines of varying thickness. The base colour can vary from individual to individual. The surface also bears very fine granulation. Radial margins of the whorls are extended and frilled. The aperture is round, creamish white. Outer lip is wavy, and the inner lip is smooth.

106. *Murex* sp. (Linnaeus, 1758) Plate 15

Original name: N/A

Description: The shell is off white in colour, conical in shape and with elevated spire. The whorls are convex, shoulders are rounded. The whorls show latitudinal thick and thin alternating ribs. The thick ribs have small bands of brown colour. The siphonal canal is long, recurved anteriorly and bears 6 primary spines decreasing in size towards the anterior side. The body whorl has 3 primary spines, with the shorter spine in between the 2 longer spines.

**Subfamily: Rapaninae**

107. *Indothais lacera* (Born, 1778) Plate 15

Original name: *Murex lacerus*

Description: The shell is oval and fusiform in shape. It can be orangish brown, peach to grey in colour. Shoulders are prominent, angulated and bear pointed, conical projections. Aperture is cream in colour. Outer lip is denticulate, and the inner lip is smooth. The egg cases are laid in clusters. Its apex is pointed and curved.

108. *Indothais sacellum* (Gmelin, 1791) Plate 16

Original name: *Murex sacellum*

Description: The shell is oval and fusiform in shape. The colour can be yellowish tan, grey with brown patches or completely grey when covered in mud. The whorls have distinct projecting knobs on shoulders and squamose spiral cords that are separated by intervals as broad as cords. The body whorl has larger and thicker spiral cords which are closely placed. The aperture is white in colour, the outer lip bears denticulations that extend far inside the aperture. The egg cases are laid in clusters, rectangular in shape with a longitudinal depression running in the centre.

109. *Indothais blanfordi* (Melvill, 1893) Plate 16

Original name: *Purpura (Stramonita) blanfordi*

Description: The shell appears fusiform in shape and blackish grey in colour. Spire is elevated, apex is pointed. Whorls are decorated with large, rounded elevations in the centre. The body whorl has wide, spiral ribs distanced with riblets in between them. The aperture is oval with inner lip smooth. Outer lip has a waved margin and bear six longitudinal denticulations in pairs.

110. *Purpura bufo* (Lamarck, 1822) Plate 16

Original name: *Purpura bufo*

Description: The shell characteristics are similar to that of *P. persica*. Except that the spire is slightly low, and the body whorl shows large, rounded nodules. The egg cases are laid in clusters. They are cylindrical with a blunt, flat tip of the apex.

111. *Purpura persica* (Linnaeus, 1758) Plate 16

Original name: *Buccinum persicum*

Description: The shell is fusiform, spire is elevated. It is blackish grey in colour. The spire whorls are slightly convex while the body whorl is convex and narrowing anteriorly. The body whorl bears spirally arranged raised ribs with black and white colour bands alternating each other. This rib is spaced with 5 – 6 impressed lines that are black in colour and bear a row of rounded tubercles. The aperture is white in colour with a slight orange tinge and oval. The outer lip has a crenulated margin, and its inside are smooth. The egg cases are laid in clusters, anchored to rocks and they are somewhat bilaterally flat, arch shaped with a slight depression in the centre at the apex, appearing like a tooth.

112. *Semiricinula tissoti* (Petit de la Saussaye, 1852) Plate 16

Original name: *Purpura tissoti*

Description: It is often confused with *I. blanfordi* because visually it’s very similar looking. It has transverse bisulcate grooves and spiral raised ridges of equidistant rounded nodules. Egg cases are laid in clusters and anchored to a rock. Each case is cylindrical with a rounded tip at the apex, like a finger.

113. *Reishia* cf*. bitubercularis* (Kuroda & Habe, 1971) Plate 16

Original name: *N/A*

Description: The shell is fusiform, black in colour with white interrupted bands alternated by 2 to 3 flat ridges on the whorls. The spire is elevated, whorls appear concave with nodular shoulders. Body whorl shows two spiral rows of large, rounded nodules. The aperture is oval in shape. The outer lip shows black colouration towards the inside edges and margin appears wavy, denticulations are absent, while the inner lip is smooth and creamish orange in colour.

**Subfamily: Ergalataxinae**

114. *Ergalatax contracta* (Reeve, 1846) Plate 16

Original name: *Buccinum contractum*

Description: The shell is uniformly dark reddish brown in colour. It is fusiform in shape; the spire is elevated with pointed apex. The whorls are convex and bear longitudinal, raised and rounded ridges. It is also structured with spirally arranged flat ribs. The aperture is oval, elongated and white in colour. Inner lip is smooth while the outer lip bears denticulations on the inside.

115. *Lataxiena bombayana* (Melvill, 1893) Plate 16

Original name: *Murex (Ocinebra) bombayana*

Description: The shell is fusiform in shape, elevated spire and creamish to off white in colour. It has longitudinally raised ridges and spirally raised prominent ribs or fasciae which are brown in colour. The aperture is oval. The outer lip is thick and bears denticulations on the inside.

116. *Arakawania granulata* (Duclos, 1832) Plate 17

Original name: N/A

Description: The shell is uniformly black in colour and fusiform in shape. The spire is elevated. The whorls are decorated with spirally arranged rounded nodules that are closely placed.

### Family: Clavatulidae

117. *Turricula javana* (Linnaeus, 1767) Plate 17

Original name: *Murex javanus*

Description: The shell is deep brown in colour. The spire is tall and pointed. Body whorl is longer than the spire and slightly inflated. Small nodules are obliquely arranged in the middle region of the body whorl, forming a spiral ridge. The aperture is wide, and shoulders are angular. Large nodules on the spire are lighter in colour.

### Family: Conidae

118. *Conus achatinus* (Gmelin, 1791) Plate 17

Original name: *Conus achatinus*

Description: The shell is conical and the last whorl is convex. The spire is elevated. The aperture widens at the base. The colour patterns of the shell vary from individual to individual. Blotches of white, brown, black, or orange form distinct patterns on the last whorl.

119. *Conus biliosus* (Roding, 1798) Plate 17

Original name: *Cucullus biliosus*

Description: Shell is elongated and conical with a slightly elevated spire. The last whorl outline is straight and tuberculate. The shoulder is angulated. The periostracum is reddish brown in colour, white colour longitudinal striations and red spots placed equidistant covering the last whorl.

120. *Conus hyaena* (Hwass, 1792) Plate 17

Original name: *Conus hyaena*

Description: Elongated, conical shell with spire slightly elevated and convex last whorl. The shoulder is slightly convex, or sub angulated. The periostracum is orangish brown in colour and has white longitudinal striations.

### Family: Turridae

121. *Annulaturris amicta* (E. A. Smith, 1877) Plate 17

Original name: *Pleurotoma amicta*

Description: The shell is conical in shape; elevated spire and the whorls are straight. It is greyish white in colour. The whorls are decorated with spiral raised lines with spaces in between. The aperture is white in colour, elongated and oval. The inner lip is smooth while the outer lip appears wavy because of the lines of the body whorl. The outer lip also shows a small fold posteriorly.

122. Unidentified Plate 17

Original name: Turridae (H. Adams & A. Adams, 1853)

Description: The shell is chalky white in colour. It is conical in shape and the spire is elevated. The whorls are convex and bear longitudinal, raised and rounded ridges. They are also structured with latitudinal flat ribs. The aperture is also white coloured, elongated and oval. The inner lip is smooth and upturned. The outer lip is smooth too.

### Family: Terebridae

123. *Duplicaria spectabilis* (Hinds, 1844) Plate 18

Original name: *Terebra spectabilis*

Description: The shell is conical in shape and the spire is elevated. The whorls are straight, sutures are slightly impressed. It is reddish brown in colour with a cream colour band placed just above and below the suture. The whorls are structured with longitudinal raised lines in two distinct sets, one is short, placed sub suturally while the longer ones are placed beneath the shorter ones. The aperture is elongated, and both the lips are smooth.

### Family: Columbellidae

124. *Anachis terpsichore* (G. B. Sowerby I, 1822) Plate 18

Original name: *Columbella terpsichore*

Description: Shell is oval acuminate and pointed at apex. Body whorl is more than 2/3rd of the spire. Prominent axial ridges are seen. The outer lip is thick. The aperture is wide and is finely toothed at its interior margin at the base. Periostracum is white to cream in colour with orangish brown streaks, making distinct patterns.

125. Unidentified Plate 18

Original name: Columbellidae (Swainson, 1840)

Description: The shell is fusiform in shape, elevated spire and convex whorls. The sutures are slightly deep. The shell displays an intricate pattern of black, brown, orange and cream colours.

### Family: Melongenidae

126. *Volegalea cochlidium* (Linnaeus, 1758) Plate 18

Original name: *Murex cochlidium*

Description: The shell is reddish brown in colour. The spire is elevated, body whorl is 2/3rd of the spire. The body whorl is inflated. Shoulder bears prominent pointed turbercles. Aperture is oval and wide. Outer lip is finely ridged, and the inner lip is smooth. The egg cases measure about an inch in length, are laterally flat, concave at the top and bulging tips at the sides.

### Family: Nassariidae

**Subfamily: Nassariinae**

127. *Nassarius stolatus* (Gmelin, 1791) Plate 18

Original name: *Buccinum stolatum*

Description: Shell is conical, spire is elevated and whorls are inflated. It is creamy white in colour with wide brown bands present at sub-sutural positions and one in the centre of the body whorl. Each whorl bears thick longitudinal ridges with spaces in between them. These ridges become almost obsolete near the outer lip on the dorsal side. The aperture is oval, the outer lip is thickened and its inside shows denticulations. The inside of the aperture shows the three spiral brown bands.

128. *Nassarius jacksonianus* (Quoy & Gaimard, 1833) Plate 18

Original name: *Buccinum jacksonianum*

Description: It is quite like *N. stolatus* with respect to its shell characteristics and colour pattern. *N. stolatus* is bigger than *N. jacksonianus*.

129. Nassarius foveolatus (Dunker, 1847) Plate 18

Original name: *Buccinum foveolatum*

Description: The shell is narrow, somewhat oval and elongate. It is greyish purple in colour with a dark band in the middle of the body whorl. Spire is elevated. Axial ribs are present with unequal interspaces in between them. Sutures appear cream in colour and nodulose on the spire.

130. *Nassarius nodiferus* (Powys, 1835) Plate 19

Original name: *Nassa nodifera*

Description: The shell is pale or ashy white in colour with bands of cream and dark red brown, conical and ovate in shape. Spire is elevated and the whorls are convex. It is longitudinally ribbed; ribs being placed closely and have enamel like shine. The suture line appears to have knobs aligning with the ridges. The aperture is oval. The inner lip is thin, and columella is extended as a thin layer. The outer lip is thick and plain with numerous ridges on its inside.

**Subfamily: Bullinae**

131. *Bullia* sp. (Gray, 1833) Plate 19

Original name: N/A

Description: The shell is conical in shape; spire is elevated and whorls are convex. It is grey in colour. The whorls are smooth and show a line sub suturally and several spiral lines towards the aperture.

### Family: Pisaniidae

132. *Cantharus spiralis* (Gray, 1839) Plate 19

Original name: *Cantharus spiralis*

Description: Shell is rhomboid in shape. Colour is pale creamish with brown longitudinal blotches. Spire is elevated and the apex is pointed. The whorls are structured with widely spaced prominent, thickened grooves and in between them are spiral threads. Shoulders are sunken. Aperture is oval and white in colour. Outer margin and inner margin are denticulated. The egg cases are convex and have a round margin.

### Family: Cancellariidae

133. *Scalptia scalarina* (Lamarck, 1822) Plate 19

Original name: *Cancellaria scalarina*

Description: The shell is dark reddish brown in colour. It is conical in shape with an elevated spire. The sutures are depressed, and shoulders are angulated. The whorls are concave and are structured with longitudinal, raised lines and latitudinal flat lines. The aperture is oval. Both inner lip and outer lips are thickened.

### Family: Mitridae

134. *Pseudonebularia proscissa* (Reeve, 1844) Plate 19

Original name: *Mitra proscissa*

Description: The shell is fusiform in shape, elevated spire and convex whorls. It is orange in colour and the whorls are structured with raised, spiral ribs. The aperture is elongated, and both the lips are smooth. The egg cases are small, cylindrical, tapering towards apex, and resemble grains of rice.

### Family: Olividae

**Subfamily: Agaroniinae**

135. *Agaronia gibbosa* (Born, 1778) Plate 20

Original name: *Voluta gibbosa*

Description: The shell is spindle shaped, with tapering ends. The spire is elevated, and its whorls are flat. The body whorl is convex. The colour of the shell is cream with longitudinal brown wavy lines. The aperture is slit-like and becomes wider towards the anterior end. The outer lip is smooth, while the inner lip shows several spiral raised lines.

### Family: Cypraeidae

**Subfamily: Cypraeinae**

**Tribe: Mauritiini**

136. *Mauritia arabica* (Linnaeus, 1758) Plate 20

Original name: *Cypraea arabica*

Description: The shell is oval, the spire is convex, smooth, and glossy. The base colour is greyish cream, and the network formed with dark brown spots and lines. The edges are thickened with brown bands placed randomly. The base is flat, greyish cream. The aperture is slit-like and wide. The outer lip and inner lip bear numerous teeth and are yellow to brown in colour. The juvenile stage looks like olive shells, with the spire sticking out, convex whorls and a distorted colour pattern. The egg cases are triangular in shape and have pink to orange tinge.

**Subfamily: Erosariinae**

137. *Monetaria moneta* (Linnaeus, 1758) Plate 20

Original name: *Cypraea moneta*

Description: Shell is oval, wider at the anterior end and narrow towards the hind end. It is smooth, glossy, and yellowish white in colour. The spire is convex and elevated. A greyish blotch is present in the centre of the elevated spire. The edges are thickened. The aperture is slit-like, wide and both the lips bear teeth.

138. *Naria turdus* (Lamarck, 1810) Plate 20

Original name: *Cypraea turdus*

Description: The shell is oval and uniformly white in colour, speckled with yellowish brown spots of irregular sizes. The spire is elevated and convex. The edges are thickened. The base is white, and the aperture is wider at the anterior end and narrower towards the hind end. Both the outer and inner lips have numerous teeth that extend up to the columella and are callous.

**Subfamily: Erroneinae**

139. *Erronea pallida* (J. E. Gray, 1824) Plate 20

Original name: *Cypraea pallida*

Description: Shell is ovate with the front end slightly wide and hind end is rounded. The spire is convex, smooth, and glossy. It is pale cream or greyish in colour and speckled with numerous brown spots. The edge is thickened, pale cream in colour with large purplish brown spots. The aperture is slit-like and wide. The inner lip bears teeth that are blunt and extend up to the columella. The outer lip has smaller teeth and is pointed.

**Tribe: Bistolidini**

140. *Palmadusta lentiginosa* (J. E. Gray, 1825) Plate 20

Original name: *Cypraea lentiginosa*

Description: The shell is oval; the spire is elevated and convex. It is white in colour and speckled with brown spots. Three brownish obscure wavy bands span the width of the spire. The edges are thickened. The base and edges also have distinct brown spots scattered. The aperture is wide. The inner and outer lips appear ribbed. The juvenile has a white coloured shell with orange spots placed spirally.

### Family: Ovulidae

**Subfamily: Prionovolvinae**

141. *Cuspivolva* sp. (C. N. cate, 1973) Plate 21

Original name: N/A

Description: The shell is ovoid, white in colour. Very fine transversal lines can be seen on the shell. The mantle is translucent and speckled with small and distinct black spots.

142. *Primovula* sp. (Thiele, 1925) Plate 21

Original name: N/A

Description: Ovoid shell, orangish red in colour. The transversal lines are distinctly visible on the shell. The mantle is pale orangish red in colour and speckled with black spots and white raised spots.

143. *Crenavolva striatula* (G. B. Sowerby I, 1828) Plate 21

Original name: *Ovulum striatulum*

Description: The shell is ovoid and pale orangish red with a cream-coloured band. The mantle is orangish pink in colour with raised dark pink and white spots.

### Family: Velutinidae

**Subfamily: Lamellariinae**

144. *Lamellaria* sp. (Montagu. 1816) Plate 21

Original name: N/A

Description: The snail is oval in shape; the surface shows black spots on a translucent circular patch and the rest is covered by specks of irregular pale yellow and white. The circular shell is visible in the centre and is translucent white in colour.

### Family: Littorinidae

145. *Littoraria sp. 1* (Gray, 1833) Plate 21

Original name: N/A

Description: Conical shell, spire elevated and long. The whorls are slightly impressed at the suture, convex, and rounded. The colour is black with irregular zig zag or oblique off-white lines.

146. *Littoraria* sp. 2 (Gray, 1833) Plate 21

Original name: N/A

Description: The shell is conical, spire is elevated, apex pointed. Whorls are straight, are cream in colour with longitudinal marks of red and purple in an irregular spiral.

147. *Littoraria* sp. 3 (Gray, 1833) Plate 21

Original name: N/A

Description: The shell is light greyish white in colour, spire is elevated, conical in shape, whorls slightly inflated and bears brown longitudinal marks which appear to be spirally arranged.

148. *Littoraria* sp. 4 (Gray, 1833) Plate 21

Original name: N/A

Description: The shell is conical and pale cream in colour. The whorls are flat, sutures are tight and impressed. The whorls show spiral and flat bands which show orange coloured longitudinally arranged spots. The aperture is circular and the outer and inner lip is smooth.

149. *Echinolittorina malaccana* (R. A. Philippi, 1847) Plate 21

Original name: *Littorina malaccana*

Description: The shell is very small, conical and is deep grey in colour. The spire is elevated, the body whorl is convex and bears two rows of rounded nodules.

### Family: Naticidae

**Subfamily: Naticinae**

150. *Paratectonatica tigrina* (Roding, 1798) Plate 22

Original name: *Cochlis tigrina*

Description: Shell characteristics are similar to *T. lineata.* Shell is pale white in colour with light to dark brown spots. The egg case is flat and collar like dark grey in colour speckled with tiny white microscopic eggs.

151. *Tanea lineata* (Roding, 1798) Plate 22

Original name: *Cochlis lineata*

Description: The shell is globose; whorls are inflated, and the spire is slightly raised. Colour is pale white with closely placed transverse wavy pale orange colour lines. The umbilical region is white in colour. The surface is glossy. The aperture is large, both the lips are smooth. A short funicle spirally enters into the umbilical depression. A U-shaped space separates it from parietal callus.

152. *Neverita didyma* (Roding, 1798) Plate 22

Original name: *Albula didyma*

Description: Shell characteristics are like *T. lineata.* It is brownish in colouration and bears whitish grey transverse lines. Surface is smooth and glossy. The interior of the aperture and callus has purplish brown colouration.

### Family: Rostellaridae

153. *Tibia curta* (G. B. Sowerby II, 1842) Plate 22

Original name: *Tibia curta*

Description: The shell is elongated. Spire is conical, pointed, and elevated. It is pale cream to light brown in colouration which gets darkened at the sutures. Aperture is oval and tapering at both ends and white in colour. The outer lip is thick and bears 3 - 4 short and rounded finger-like projections. The inner lip is also thick, white and is turned outwards.

### Family: Bursidae

154. *Bufonaria echinata* (Link, 1807) Plate 22

Original name: *Gyrineum echinatum*

Description: The shell is elongated and is light brown in colour and dark brown markings can be found occasionally. The spire is elevated and bears granular spiral ribs that are closely placed and has peripheral rows of small nodules. Adjacent to the interior canal is a strong recurved spine. The outer lip’s inner margin is flared and bears irregular shaped teeth on the outer margin.

### Family: Cymatiidae

155. *Gyrineum natator* (Roding, 1798) Plate 23

Original name: *Tritonium natator*

Description: The shell is dorso-ventrally compressed and is conical in shape. The spire is elevated, and the apex is pointed. The body whorl is structured by granulation and lines in spiral cords. The spire whorls too have spiral cords. The aperture is round, white and its interior is toothed. The outer lip is varixed, strong and thick. The shell displays bands of grey, orange, pale black with reddish tinge and cream on the lateral thickened and curved sides. The egg cases are spherical and laid in a tight spiral.

### Family: Assimineidae

156. *Assiminea marginata* (Leith, 1853) Plate 23

Original name: *Optediceros marginatum*

Description: The shell is uniformly pale light brown in colour, convex whorls and tight sutures. The snail is red in colour.

157. *Assiminea subconica* (Leith, 1853) Plate 23

Original name: *Optediceros subconicum*

Description: The shell features are the same as *Assiminea marginata,* except that the body whorl is yellow in colour. The snail is white in colour.

### Family: Iravadidae

158. *Iravadia bombayana* (Stoliczka, 1868) Plate 23

Original name: *Fairbankia bombayana*

Description: The shell is pale cream in colour, whorls are convex, and the spire is elevated. The snail’s tentacles show alternate bands of white and black.

### Family: Vermetidae

159. *Thylacodes* sp.1 (Guettard, 1770) Plate 23

Original name: N/A

Description: The shell spirals loosen and almost straightens towards the aperture, pale cream in colour with slight pinkish, orange, and brown tinge. Faint ribs are visible on the shell. The snail is purplish black in colour. Oval, concave and black operculum can be seen.

160. *Thylacodes sp. 2* (Guettard, 1770) Plate 23

Original name: N/A

Description: The shell spirals appear somewhat flattened, and it is concentrically arranged. Ribs are visible on the surface and several longitudinal ridges are also seen in the centre of the shell. No operculum is visible.

161. *Thylacodes* sp. 3 (Guettard, 1770) Plate 23

Original name: N/A

Description: The snail has a blackish maroon colour, speckled with white. Operculum is black in colour, showing a spiral pattern.

### Family: Calyptraeidae

162. *Ergaea walshi* (Reeve, 1859) Plate 23

Original name: *Crepidula walshi*

Description: These shells were usually observed inside the dead shells. They are chalky white in colour and convex in shape. The spire is keeled posteriorly. Very fine concentric lines can be seen on the shell.

### Family: Epitoniidae

163. *Janthina* sp. (Roding, 1798) Plate 24

Original name: N/A

Description: The shell is smooth, purple in colour and the spire is flat with rounded whorl.

## Discussion

The present study brings to light multiple species records from the Mumbai Metropolitan Region (MMR), that collectively underscore the need for renewed taxonomic works, arising from uncertainties in species identity, historical revisions, and the limitations of original descriptions that predate modern anatomical and molecular approaches.

The study reports the first records of *Pelagella modesta* (Alder & Hancock, 1864), originally described from Visakhapatnam, Andhra Pradesh; *Cuthona annandalei* (Eliot, 1910) from Port Canning, West Bengal; and *Aplysia* cf. *rudmani* (Bebbington, 1974) from Bayt Island, Gujarat (present-day Beyt Dwarka). Additionally, five other individuals of *Aplysia sp.* were documented, the identification of which as *Aplysia* cf. *rudmani* remains tentative, as identification based solely on external morphology is inconclusive without anatomical and molecular examination.

The rediscoveries of 7 species, with the oldest one documented after more than a century are indicative of the robustness of the data collected from this effort. (Winckworth, 1946a, b; Melvill, 1893). *Glossodoris poecila*, one of those 7 rediscoveries and *Assiminea subconica* are classified as a taxon inquirendum (WoRMS, 2025), highlighting their unresolved taxonomic status.

According to Nudibranch Systematic Indices published online by Mc Donald (2006, 2009, 2021), *Glossodoris semeion* had been synonymised with *Glossodoris diardii* by Pruvot-Fol (1951a, b) and subsequently classified as *Hypselodoris semeion* by Rudman (1983). It was later regarded as a synonym of *Hypselodoris infucata* by Bhave & Apte (2013), citing data presented in Nudibranch systematic indices. Since the current study relies on taxonomical information from WoRMS, the species is listed as *Glossodoris semeion* in the checklist.

Multiple nudibranch species of *Atagema, Carminodoris* and *Cratena* are recorded as new species records. Although these genera have been previously reported from the Mumbai Metropolitan Region (MMR), the species encountered in the present study differ morphologically from those described earlier and thus require further taxonomic assessment.

In addition to the gastropods listed in this study, empty shells of *Monetaria annulus* and *Conasprella dictator* were reported from the Carter Road site. The pelagic marine gastropod *Janthina sp.* has also been included in the checklist, having been discovered washed ashore at Girgaon beach. Furthermore, some shelled gastropods from the presented checklist namely *Ellobium* sp., *Physella* sp., *Murex* sp., *Turritella* sp., *Bullia* sp., *Littoraria* sp.1, *Littoraria* sp.2, *Littoraria* sp.3, *Littoraria* sp.4, *Thylacodes* sp.1, *Thylacodes* sp.2, *Thylacodes* sp.3; and unidentified specimens belonging to families Muricidae, Turridae, Columbellidae remain unidentified to the species level due to the paucity of published literature from the region.

On assessing the conservation status of species listed in the present study, it was found that seven species - *Cassidula aurisfelis*, *Melampus sincaporensis*, *Telescopium telescopium*, *Melanoides tuberculata*, *Conus achatinus*, *Conus biliosus*, *Conus hyaena* are classified as least concern. The remaining have not been evaluated in the IUCN Red List of Threatened Species. Furthermore, none of the species included in the present study are listed under any Schedules of the Wild Life Protection Act, 2022. The presence of numerous new species records from one of the most diverse taxonomic groups in the intertidal highlights a significant knowledge gap regarding intertidal diversity in the region. The present checklist provides gastropod distribution patterns observed in intertidal habitats of the Mumbai Metropolitan Region during data collection (Table 4). A sustained long-term effort is essential to systematically assess the abundance of various groups in the region.

The coastline of the Mumbai Metropolitan Region is a highly altered one, continuously reshaped to accommodate the demands of its population and industry. The intertidal habitats are heavily impacted by anthropogenic activities such as coastal construction, untreated sewage disposal, oil spills, and land reclamations, rendering the nearshore waters one of the most degraded in the country (Bhojane et al., 2025). The presence of 163 species from the region indicates a thriving gastropod community but also highlights the lack of ecological studies from such an anthropogenically altered habitat. This checklist draws attention to one of the largest yet highly overlooked groups from the intertidal region, which serve as crucial indicators of their habitats (Kulkarni, 2017). The authors recommend the need for fine-scale ecological studies to better understand species-habitat relationships in the region. Such studies are vital to elucidate the role of gastropods in maintaining ecological stability and to assess resilience thresholds these and other species possess amid ongoing habitat alterations.

Analysis of the dataset taken for this study has revealed several new records and rediscoveries for the region, underscoring the significance of long-term citizen science efforts in uncovering biodiversity. Voluntary initiatives such as these, especially across previously unrecorded habitats, remain among the most cost-effective approaches to addressing data gaps and establishing essential baselines. The outcomes of this research highlight the immense value of such findings and further emphasise the necessity for finer-scale, taxonomic and ecological studies in a complex and dynamic urban coastal ecosystem like Mumbai.

## Acknowledgements

The authors are grateful to Harshal Karve and Mahi Mankeshwar for providing their invaluable insights in drafting the manuscript.

We also express our gratitude to D Barclay, Erwin Koehler, Giorgio Strano, Hsini Lin, Javier, Joe Rowlett, Joᾶo Pedro Silva, Juan Manuel de Roux, Junn Kitt Foon, Leslie Harris, Dr. Matt Nimbs, R. Lucine, Saryu Mae, Sayali Nerurkar, Scott Loarie, Susan J. Hewitt, Terence Gosliner, Tricia Goulding, Trond R. Oskars, Usha Parameswaran, Valentin de Mazancourt, Vie Panyarachun, Vishal Bhave, Yingyod Lapwong and Yloza Ula for helping with species identification on iNaturalist.org.

An extended thanks to Abhishek Jamalabad, Ashoi Dantra, Fatema Hirkani, Gaurav Patil, Harshal Karve, Jessica Luis, Nikhil Sathe, Pooja Pednekar, Sayee Girdhari, Sejal Mehta, Simran Mangale, Sudarshan Pawar and Xavier D’souza for their contributions to the dataset. The authors also thank Abhishek Jamalabad, Dutta Pednekar, Gaurav Patil and Xavier D’souza for their photographic contributions.

## Appendix

### Tables

Table 1: Study sites divided into Zones (Source: Authors)

Table 2: List of Historical Molluscan publications from the city (Source: Authors).

Table 3: Record of shelled Gastropods if they were observed live. (Source: Authors)

Table 4: Zone Wise distribution of Gastropods. (Source: Authors. This dataset was generated through analysis of observations from the Marine Life of Mumbai project on iNaturalist - https://www.inaturalist.org/projects/marine-life-of-mumbai).

Table 5: New Records of Gastropods from MMR. (Source: Authors)

Table 6: Rediscoveries of species originally described from Mumbai. (Source: Authors)

### Figures

Figure 1: Map of MMR region showing the Study sites. (Source: Authors)

---

**PLATE 1.**
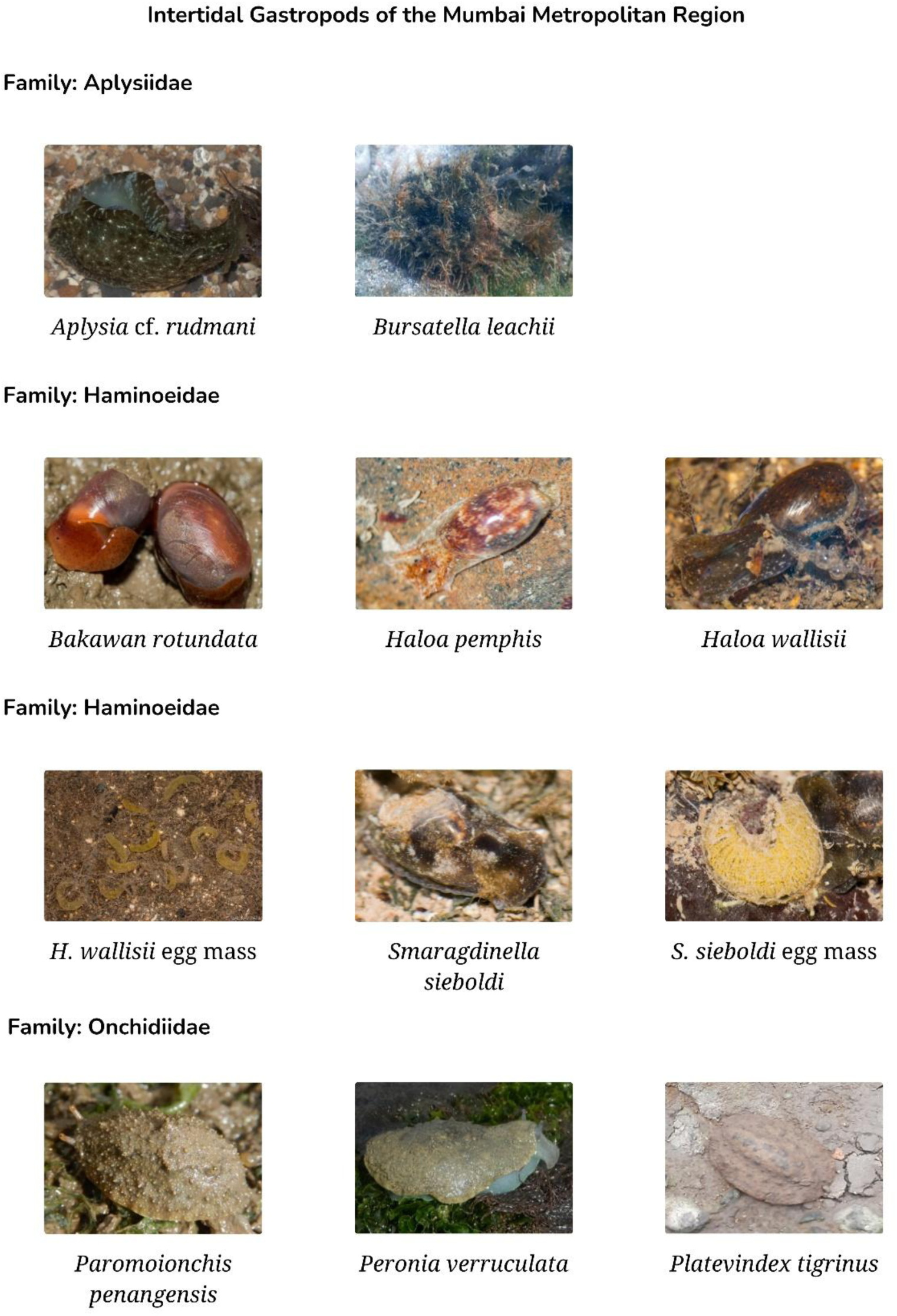

**PLATE 2.**
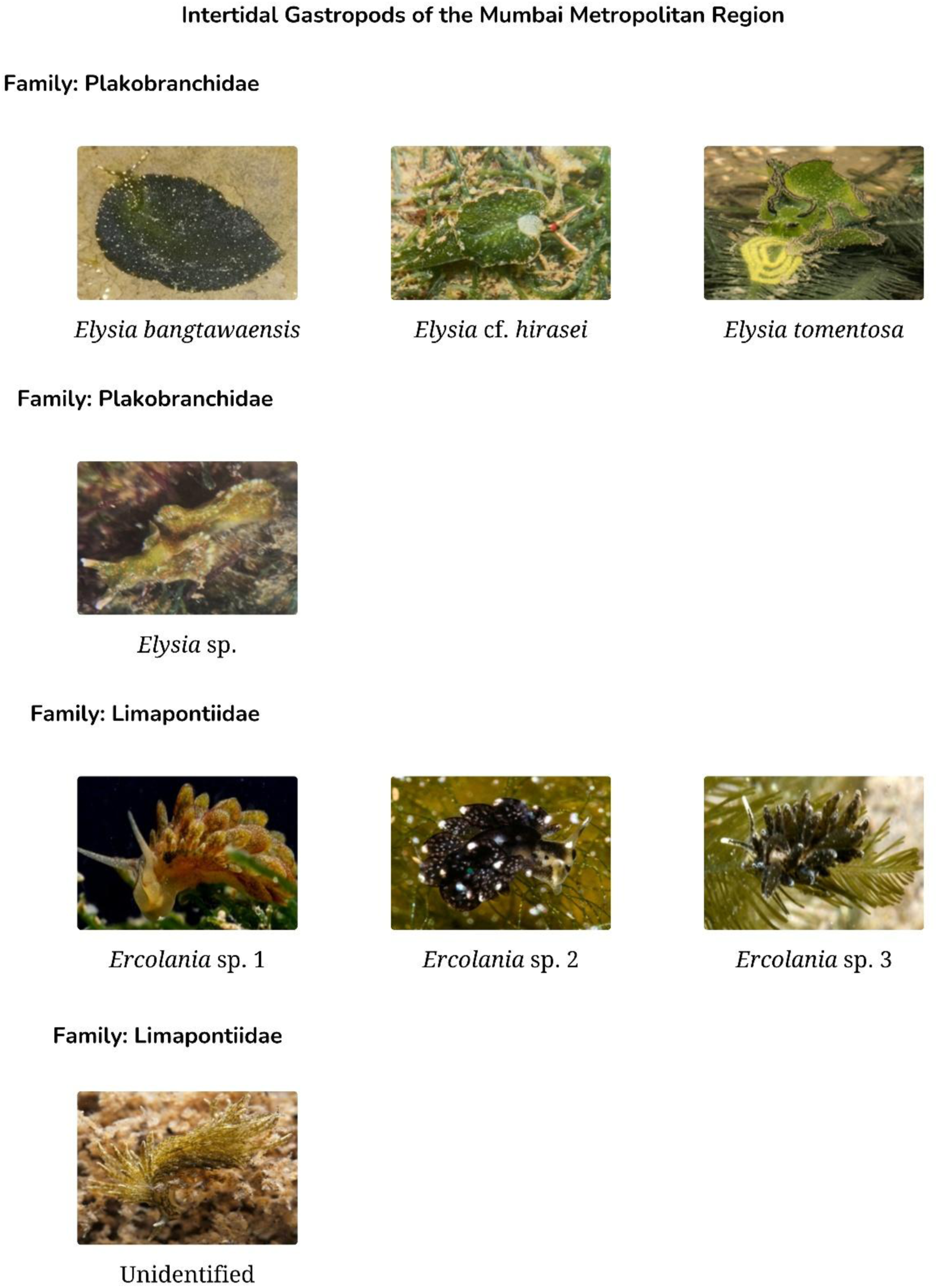

**PLATE 3.**
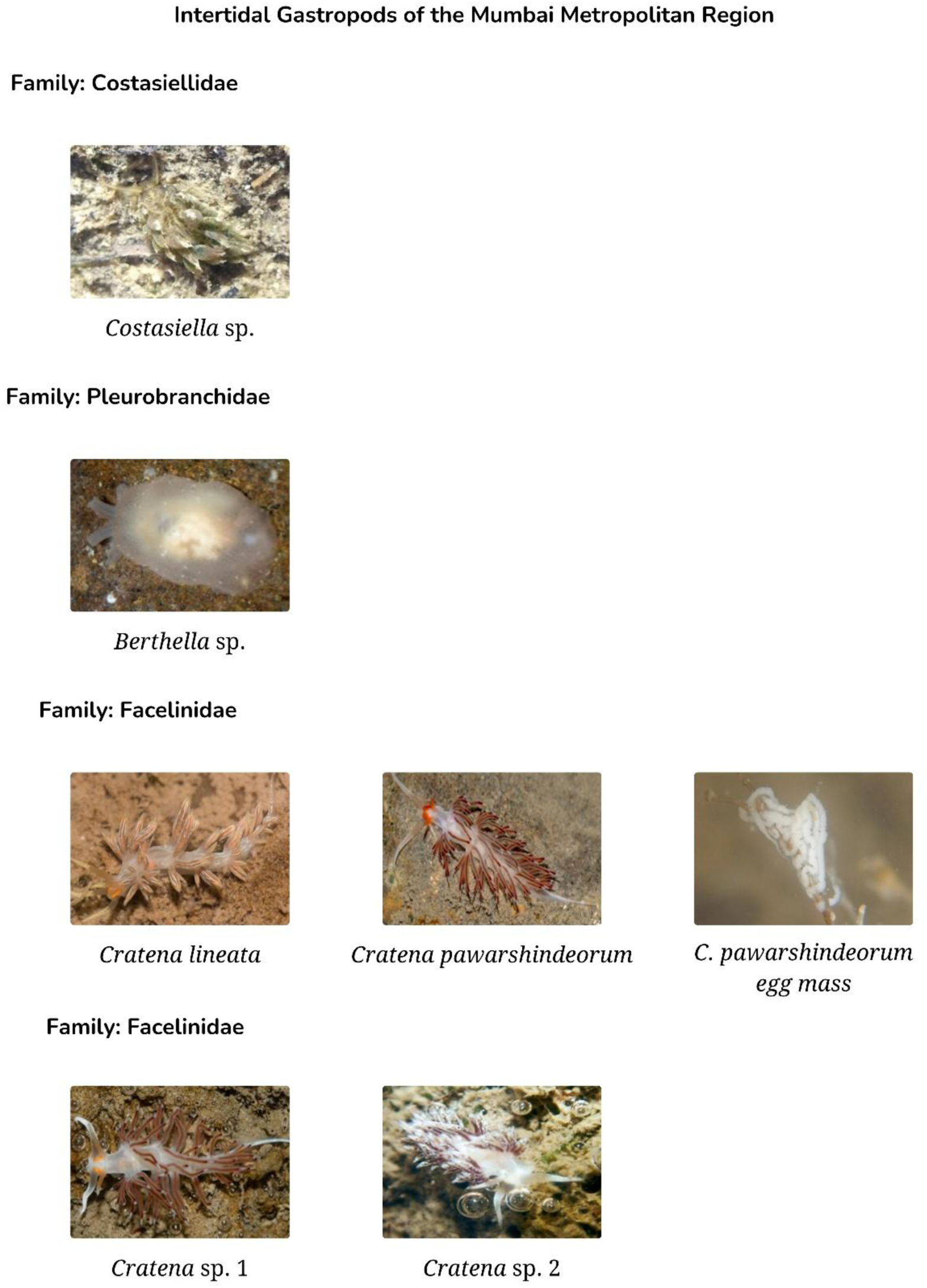

**PLATE 4.**
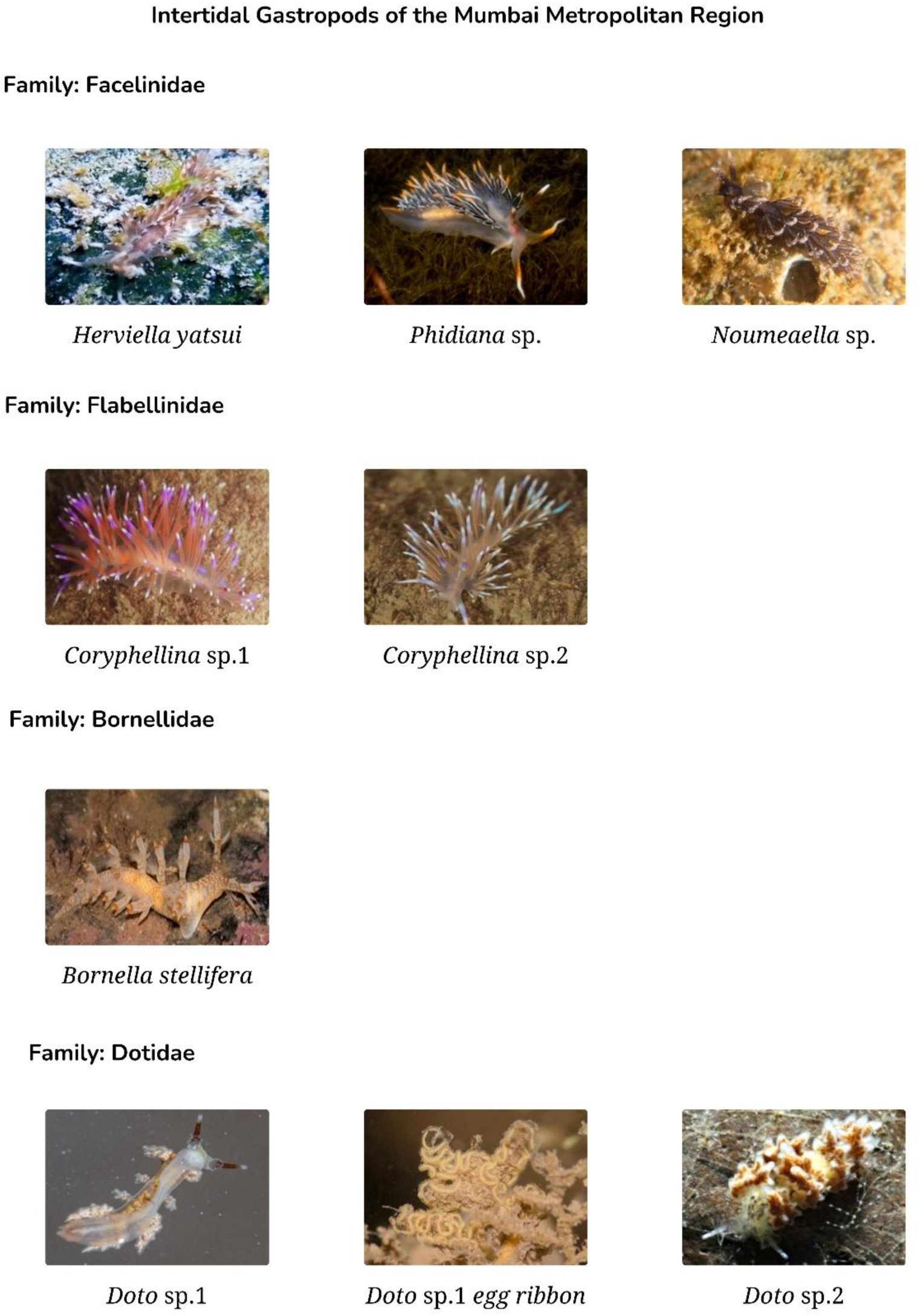

**PLATE 5.**
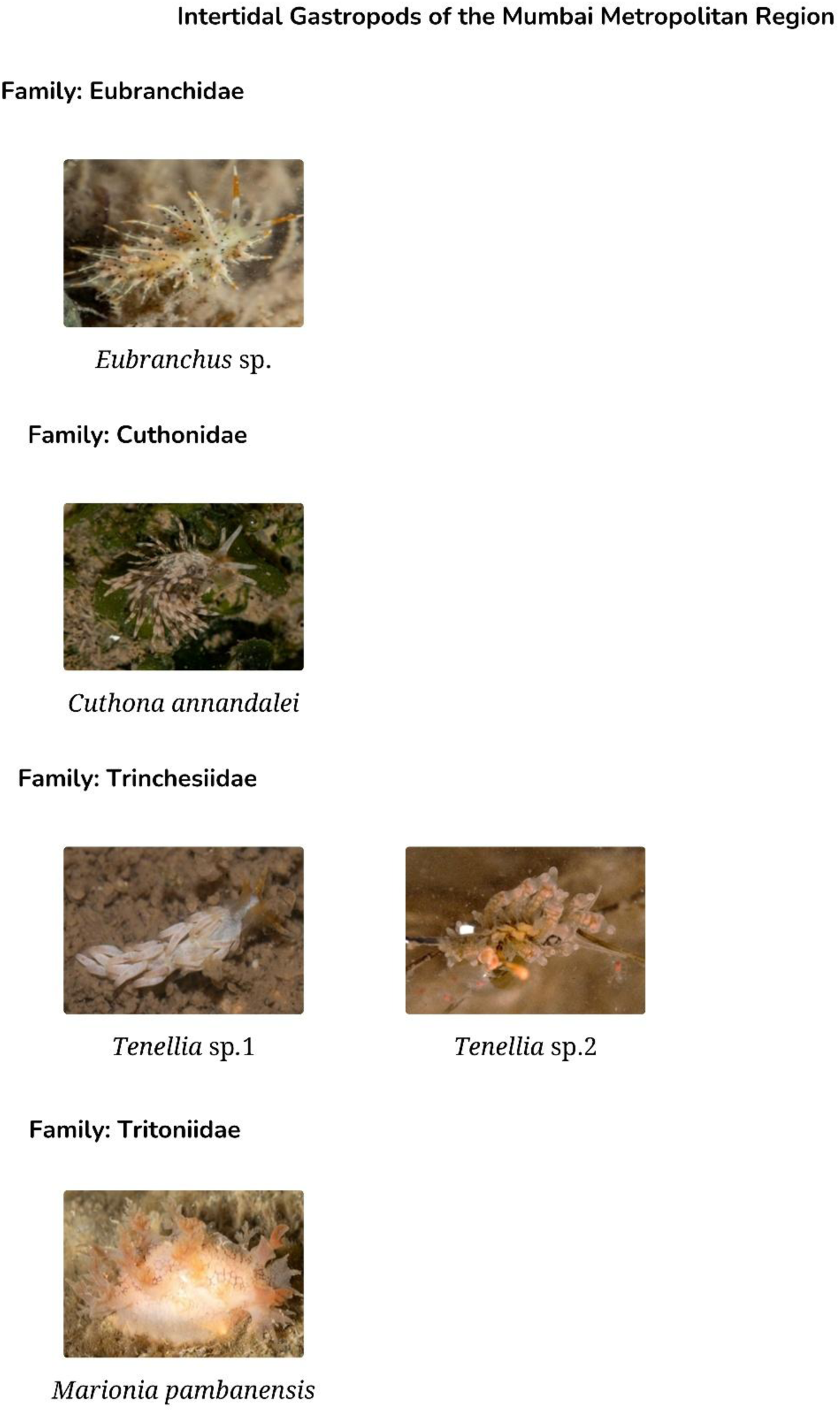

**PLATE 6.**
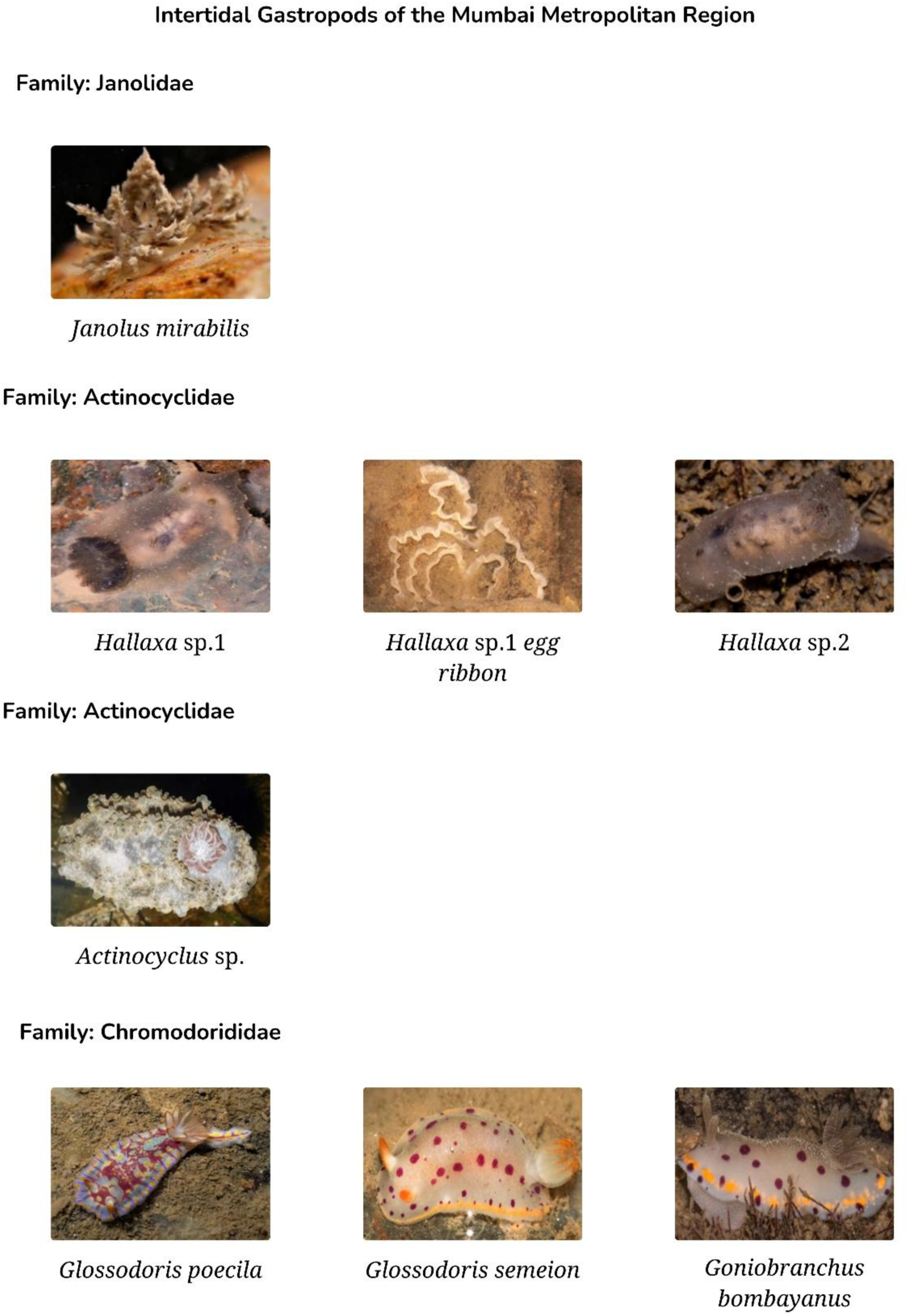

**PLATE 7.**
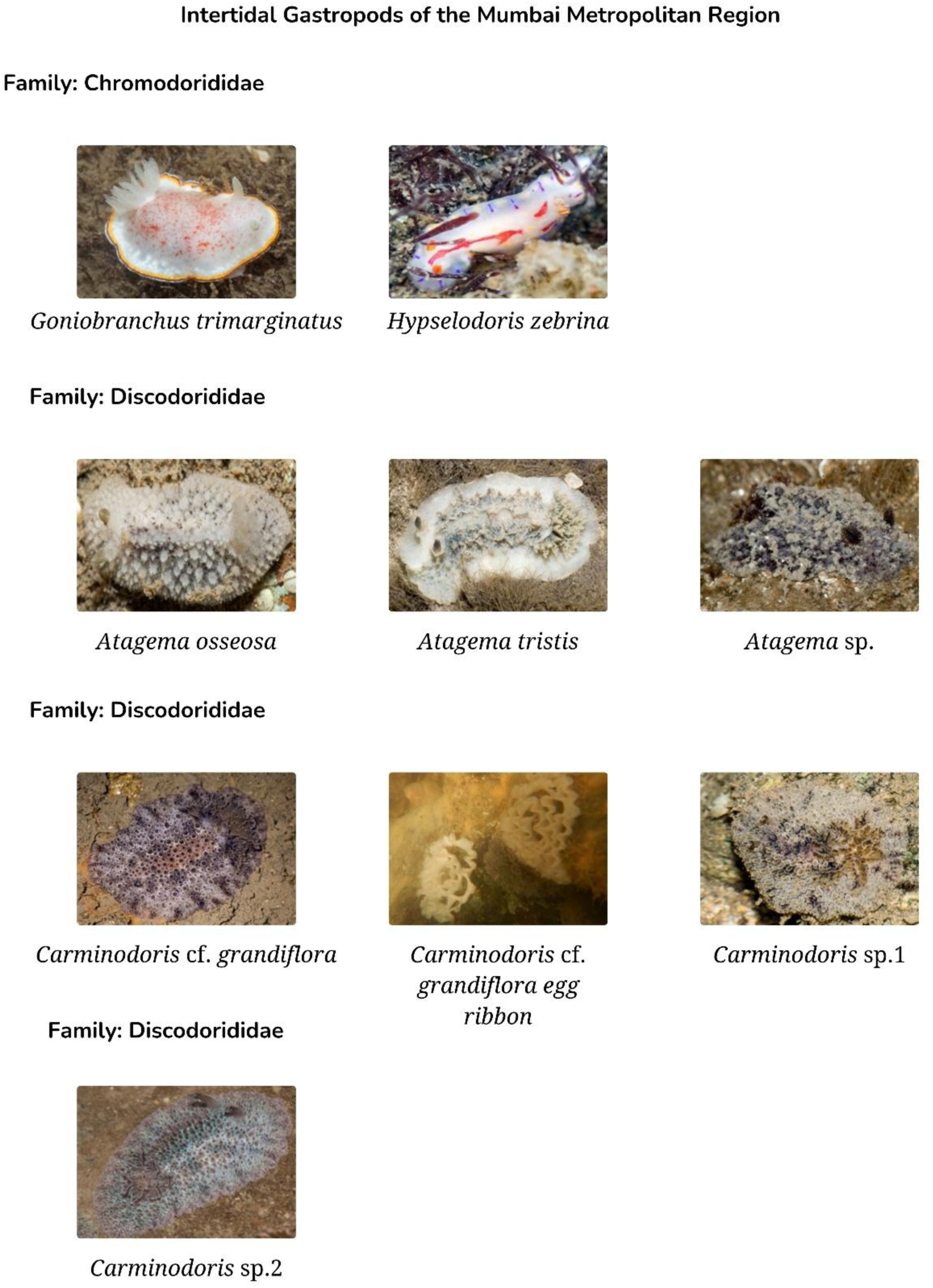

**PLATE 8.**
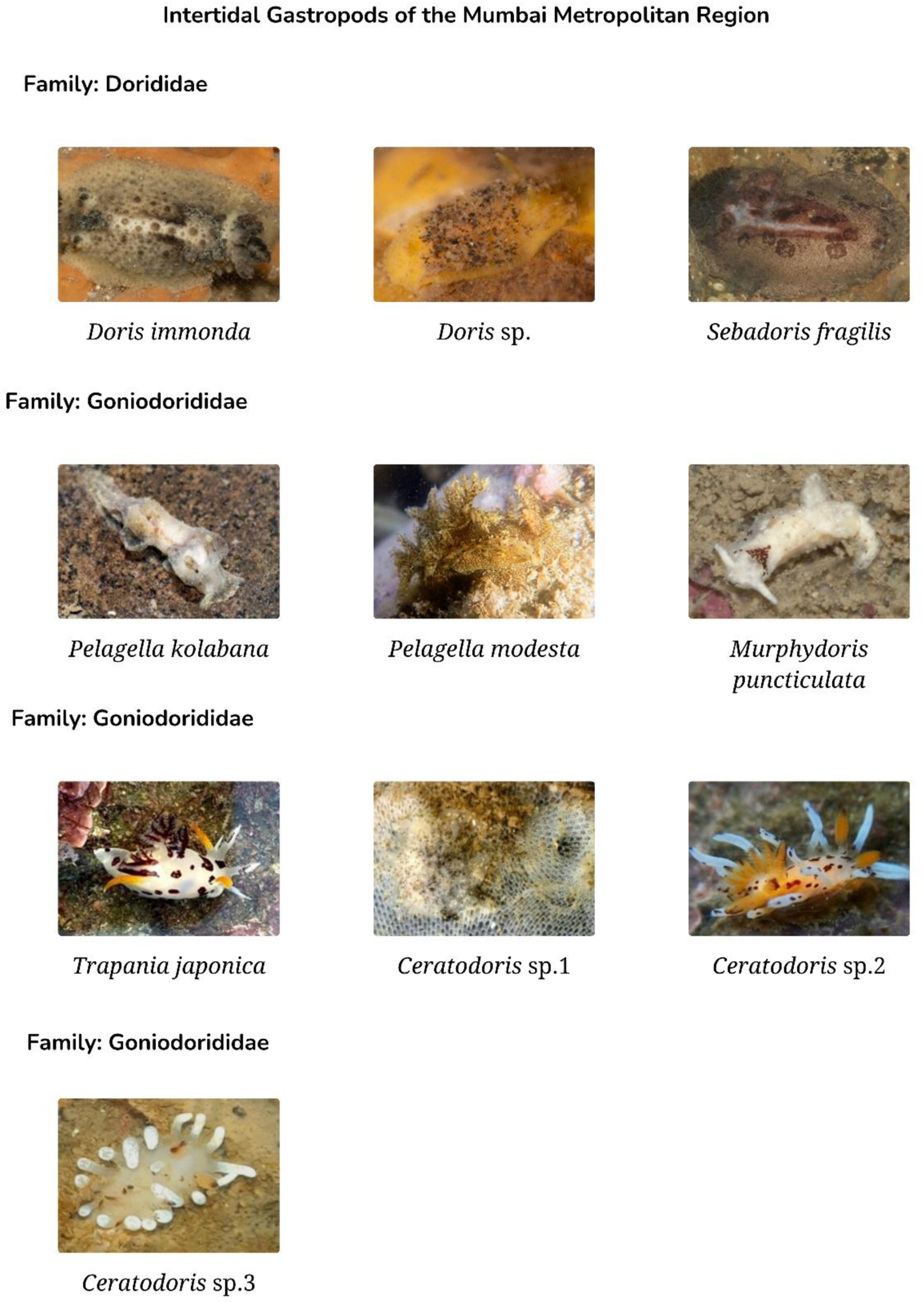

**PLATE 9.**
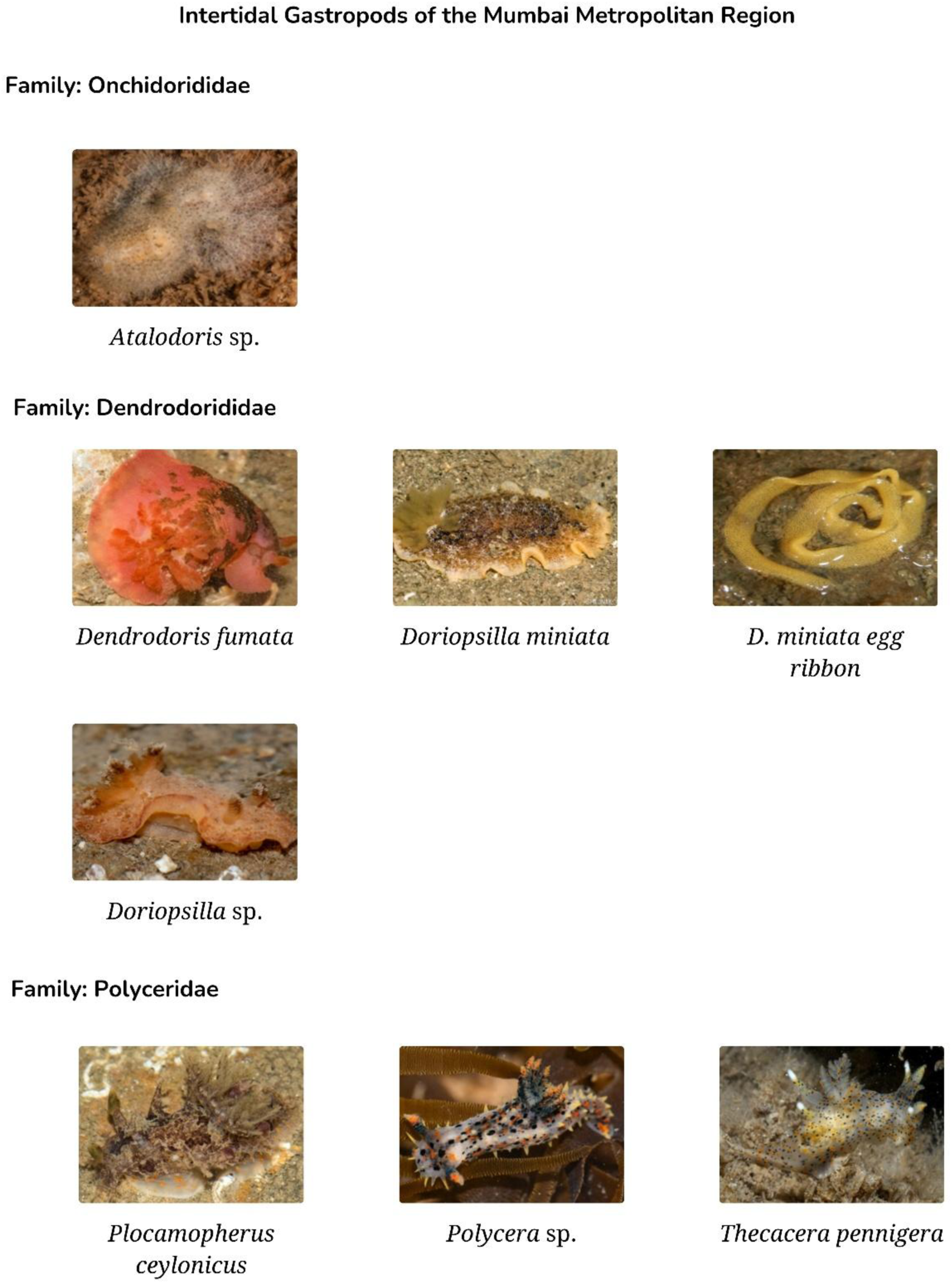

**PLATE 10.**
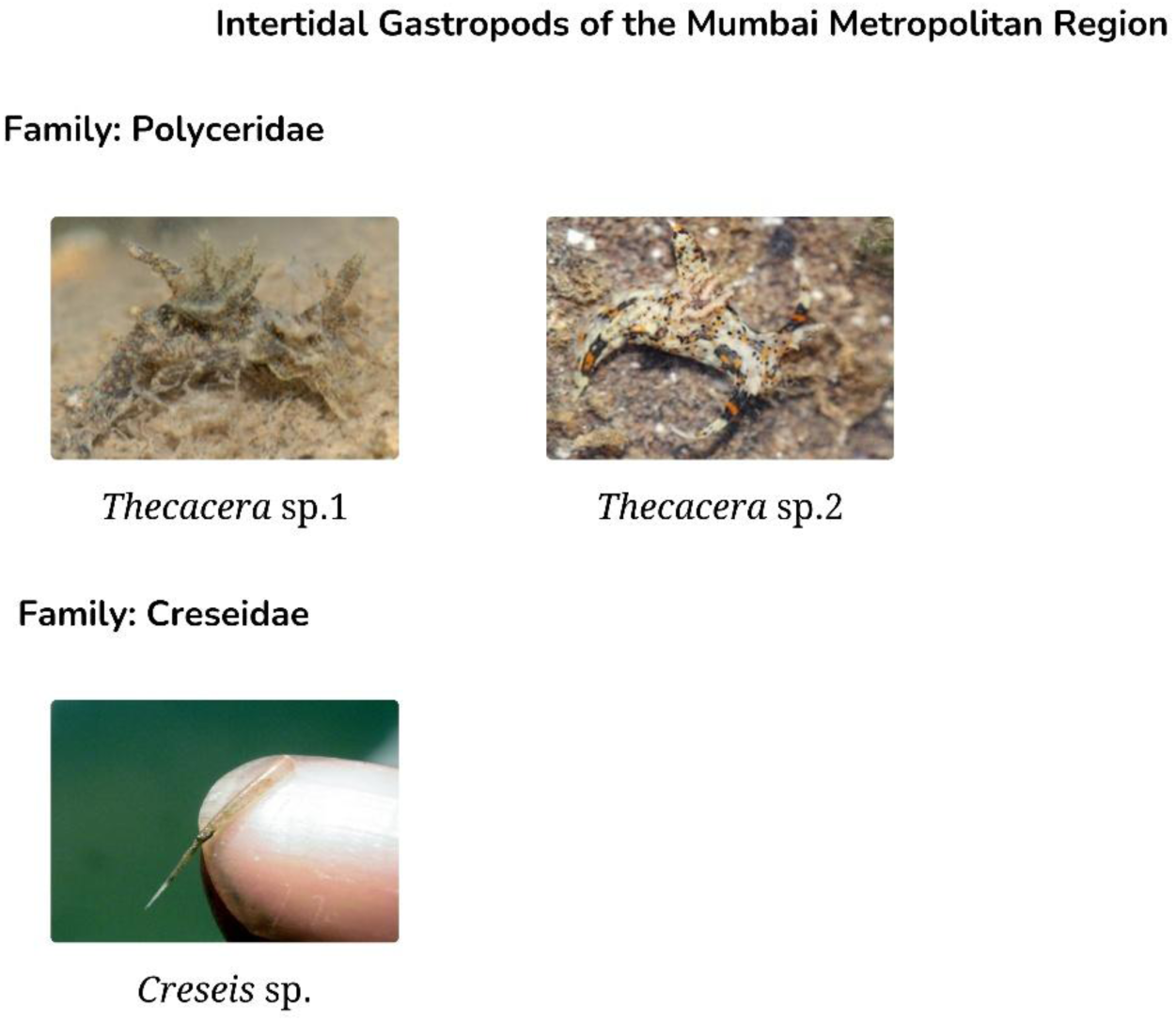

**PLATE 11.**
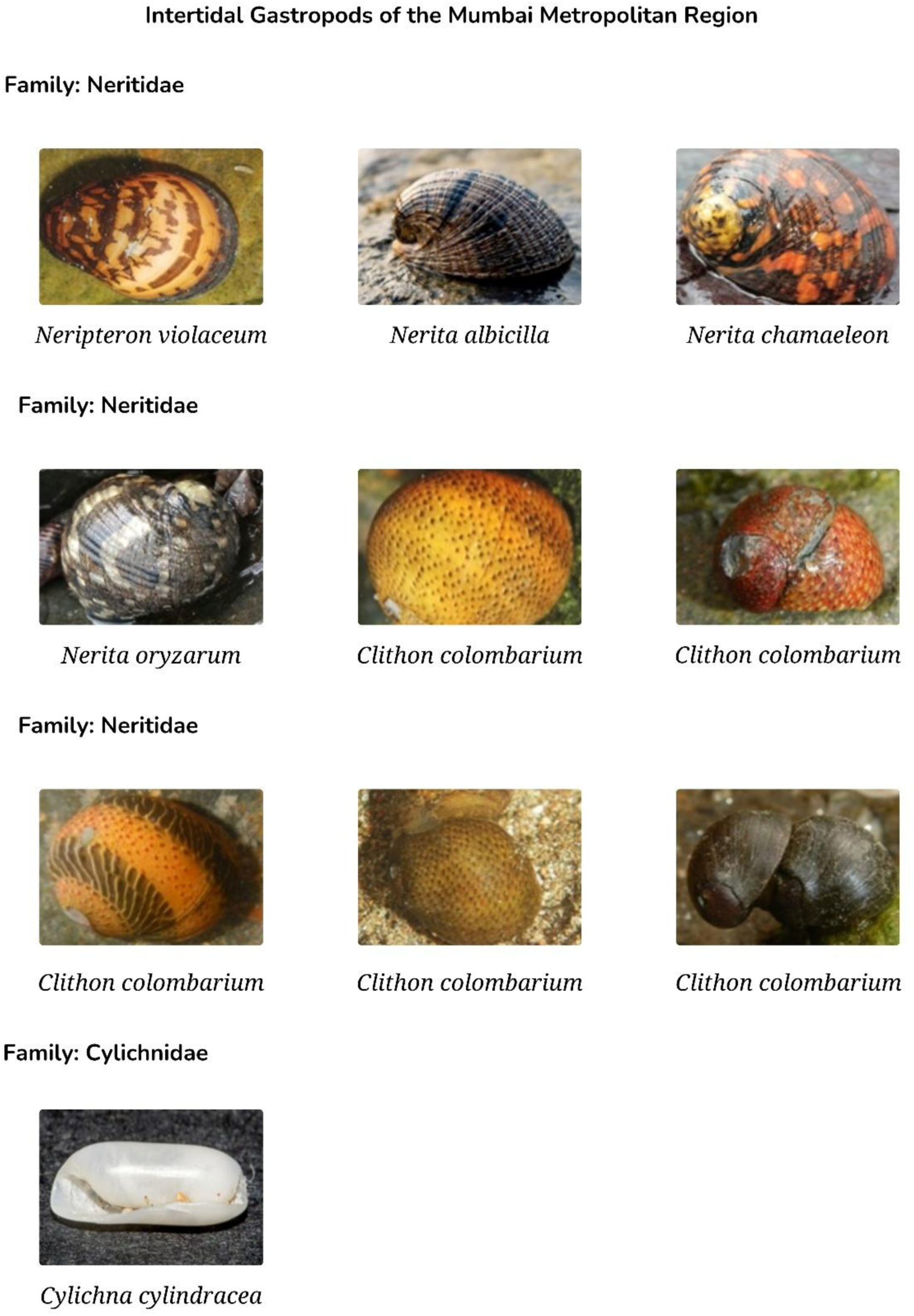

**PLATE 12.**
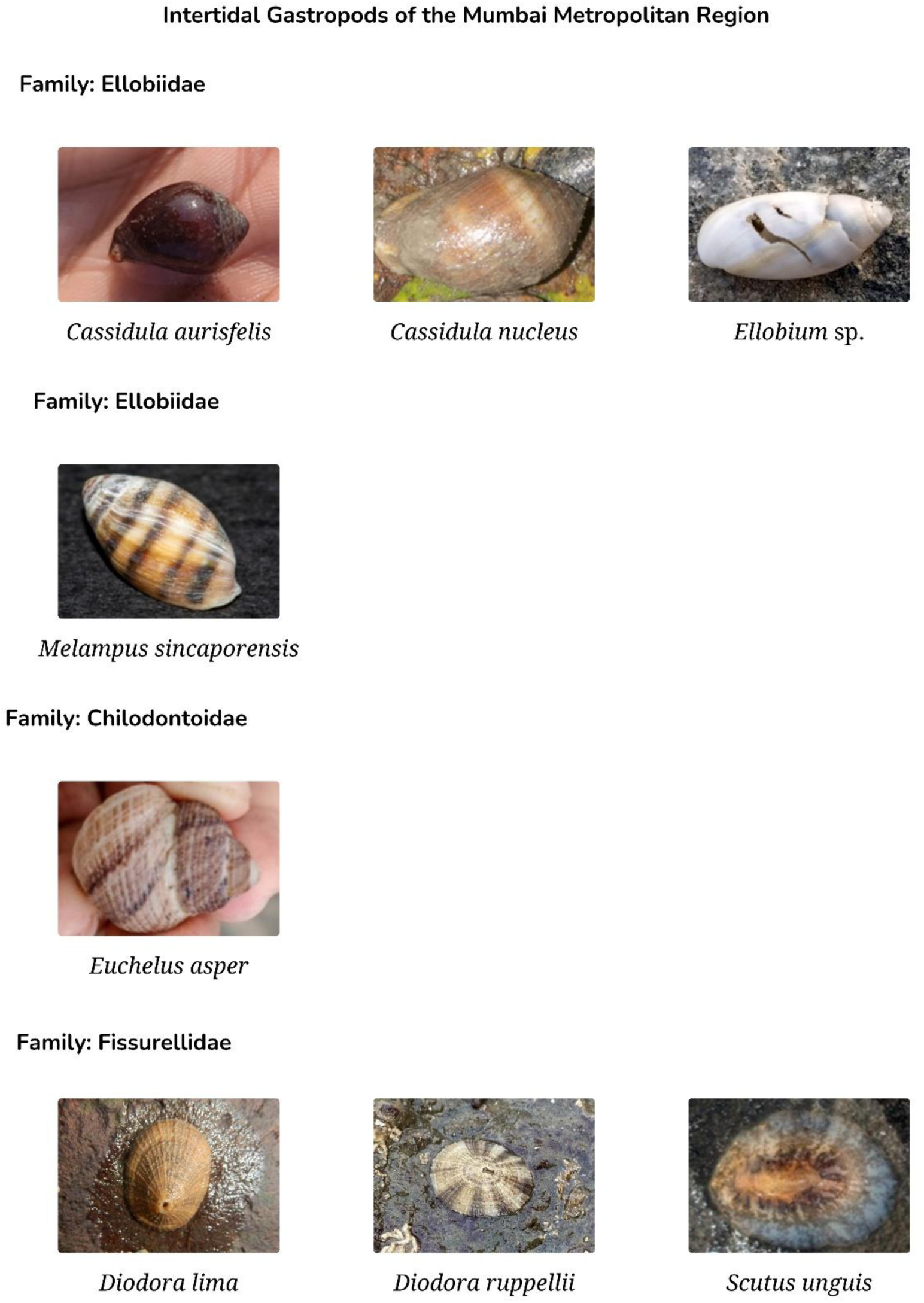

**PLATE 13.**
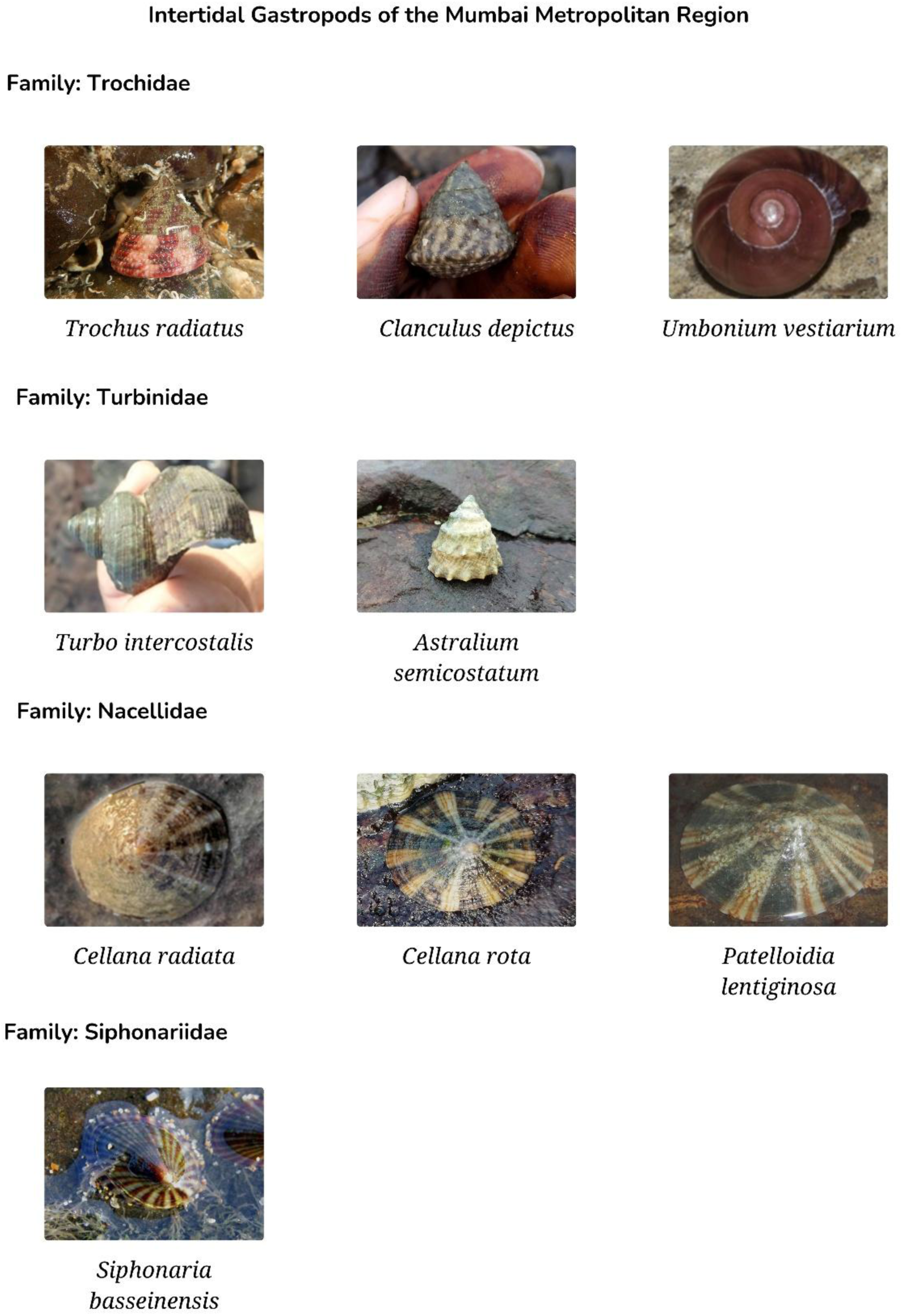

**PLATE 14.**
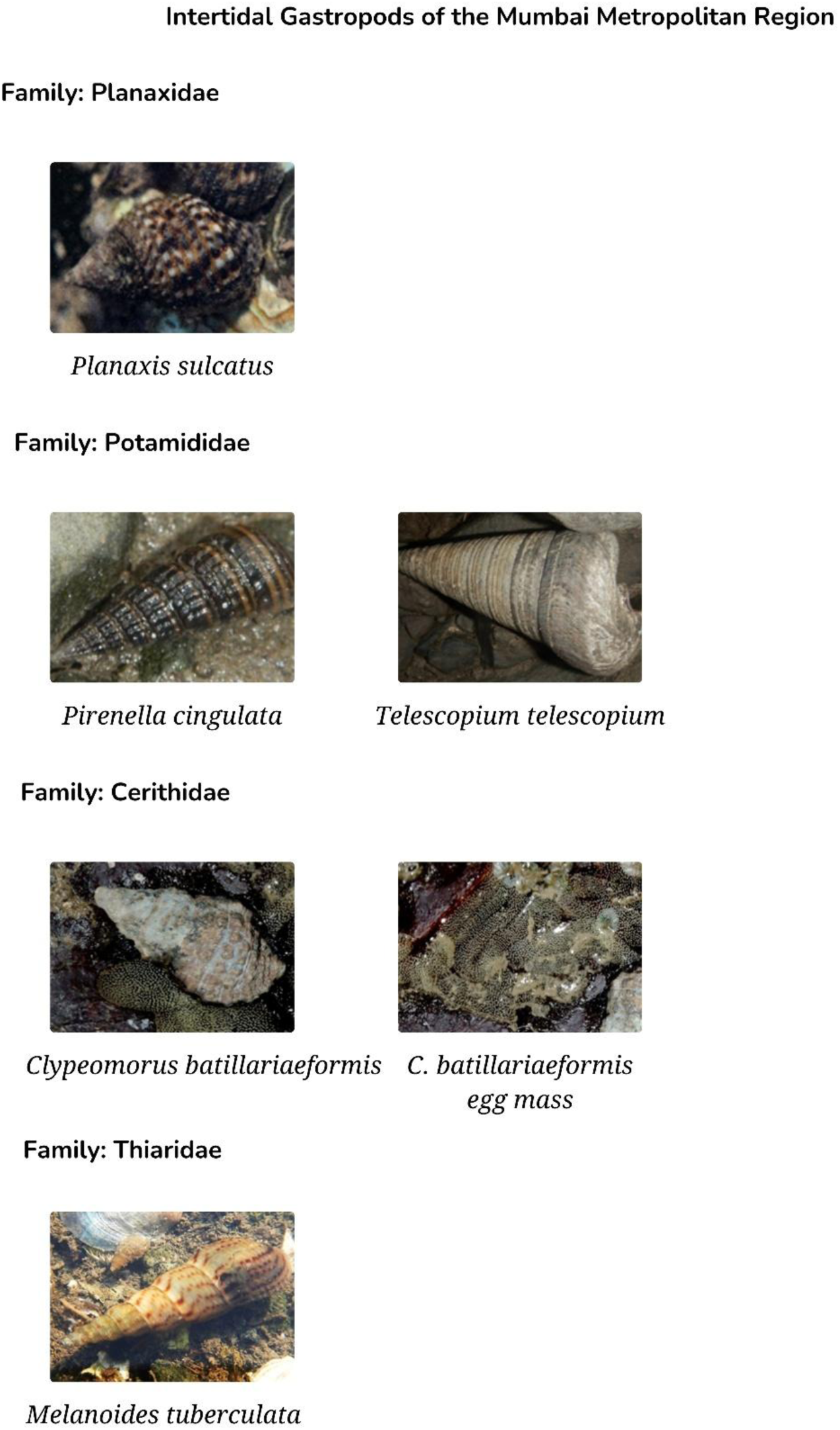

**PLATE 15.**
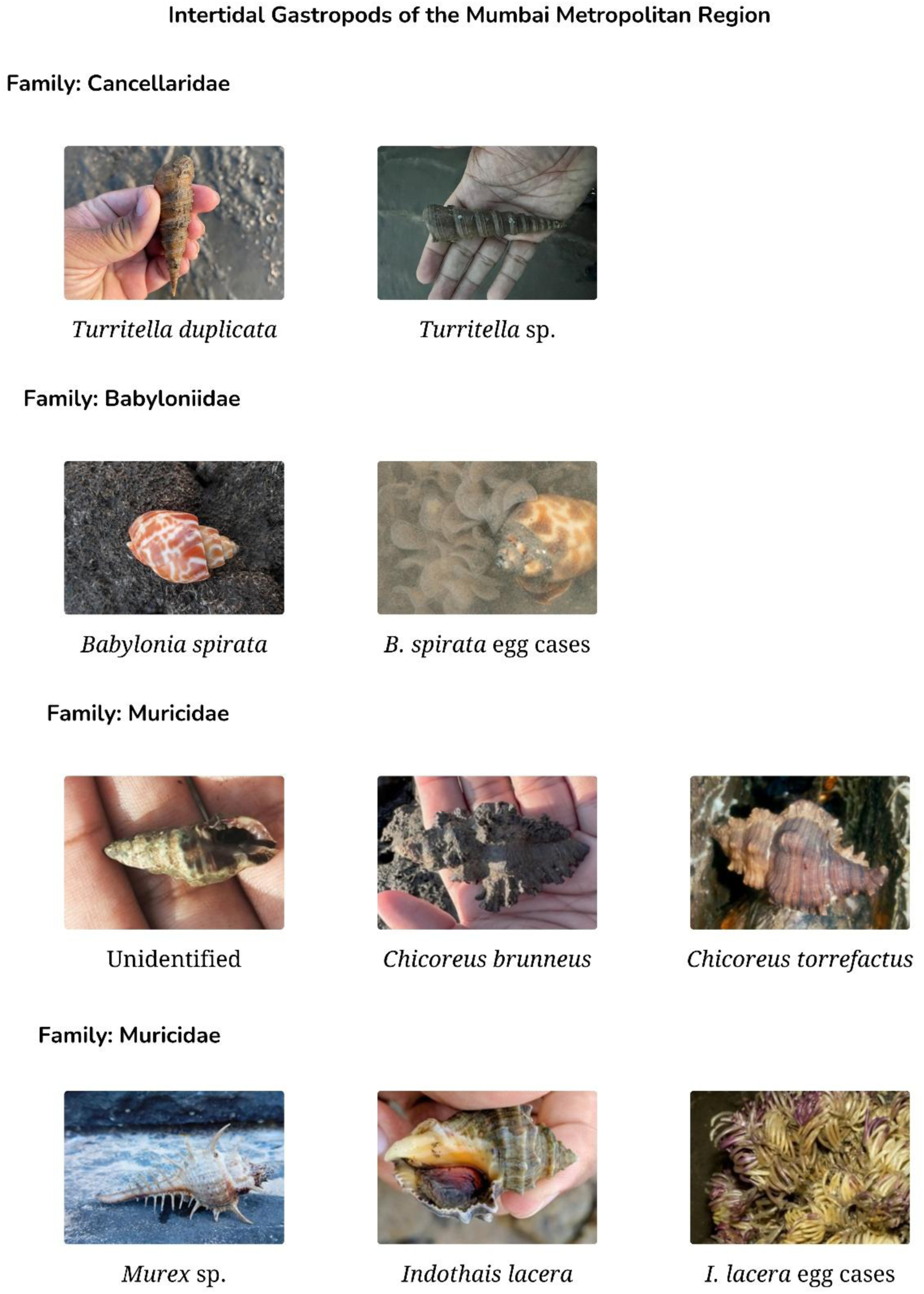

**PLATE 16.**
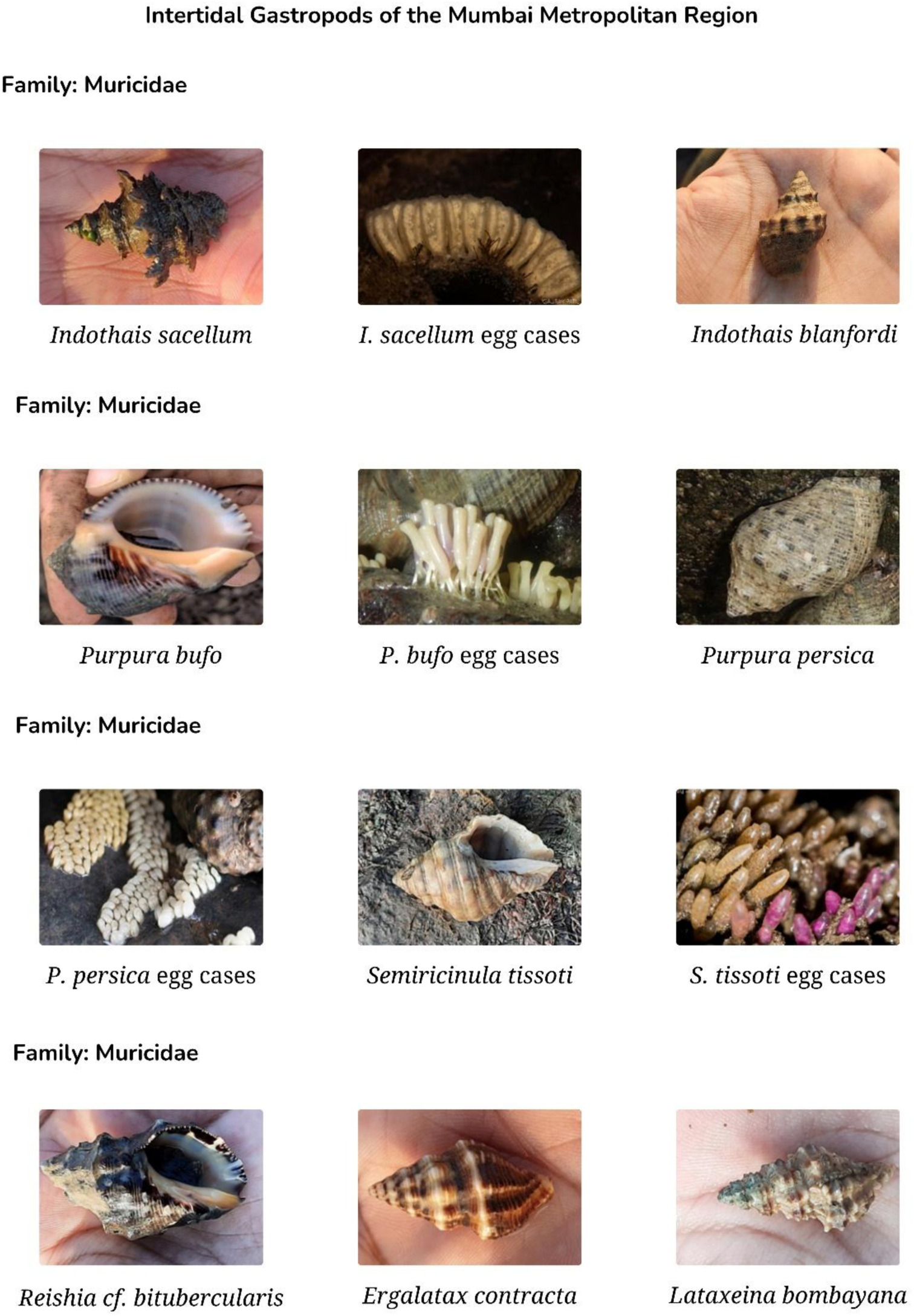

**PLATE 17.**
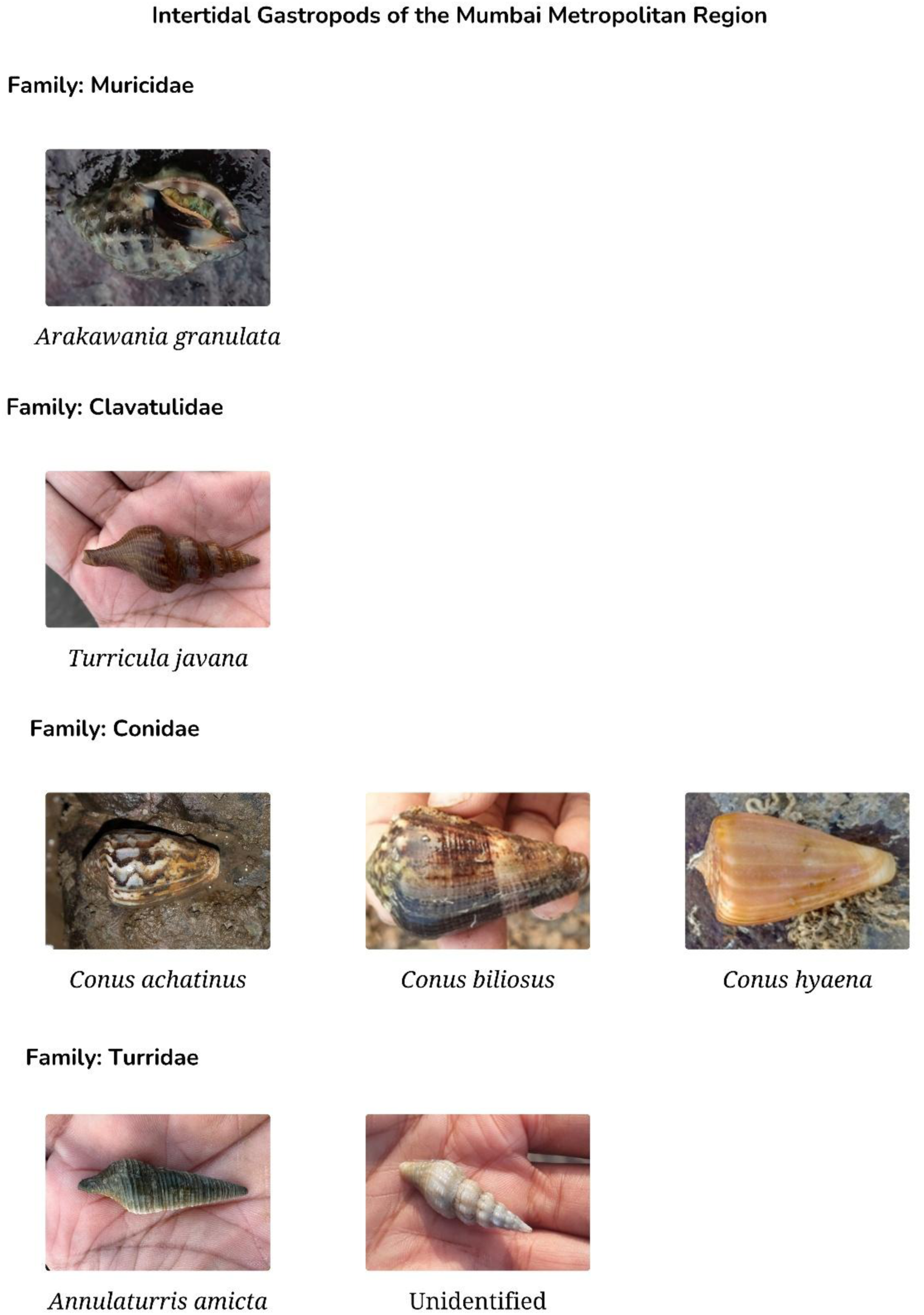

**PLATE 18.**
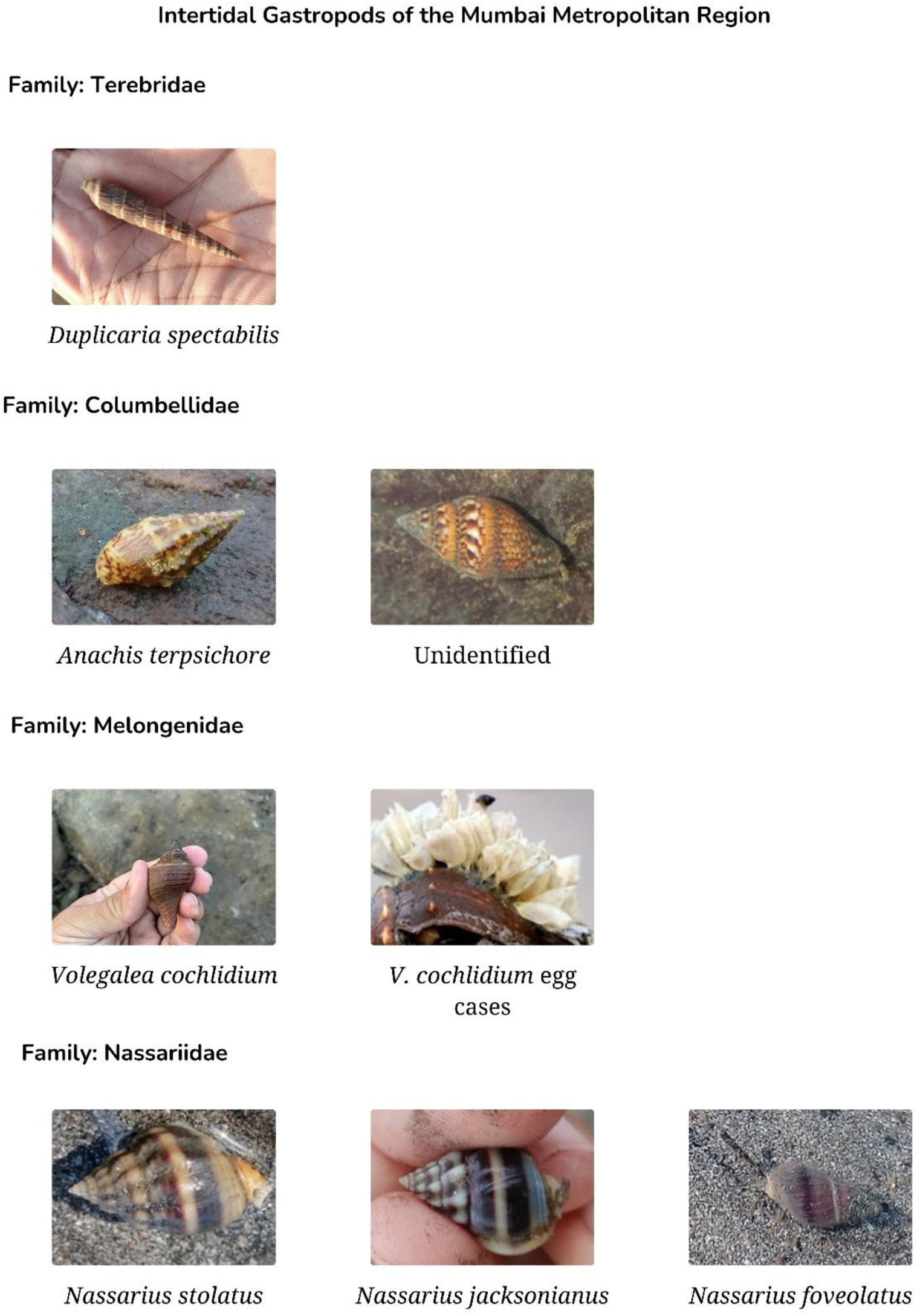

**PLATE 19.**
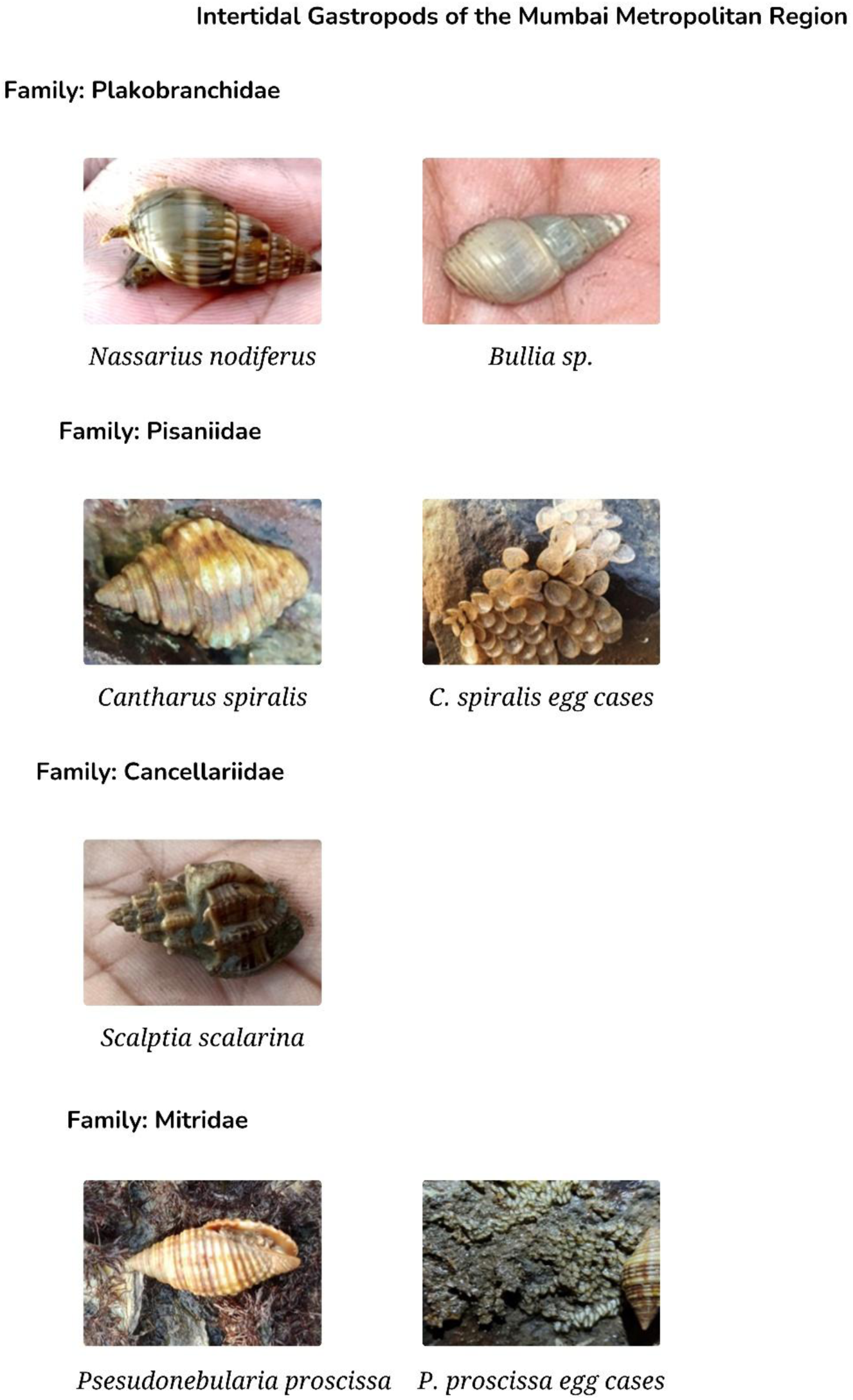

**PLATE 20.**
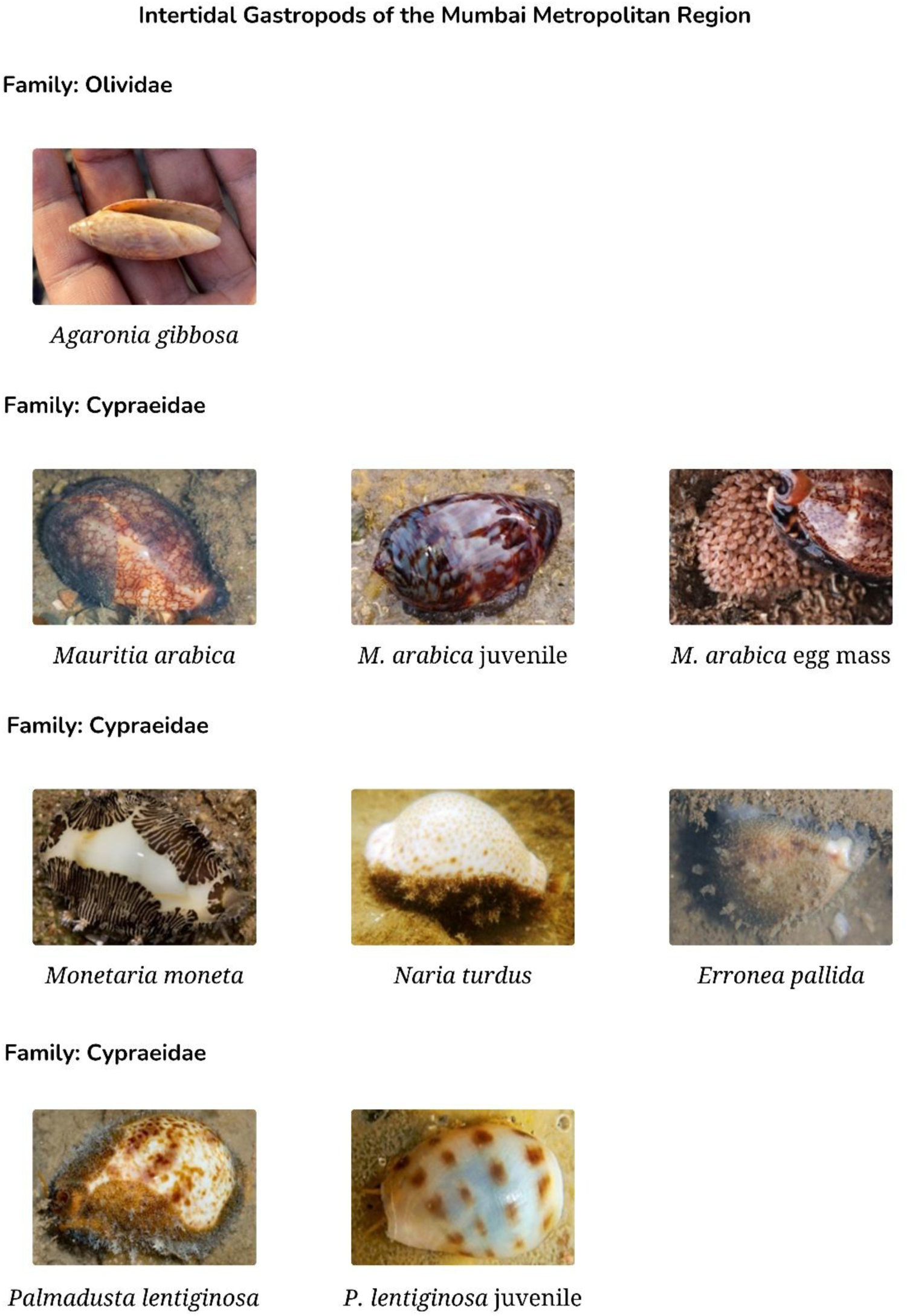

**PLATE 21.**
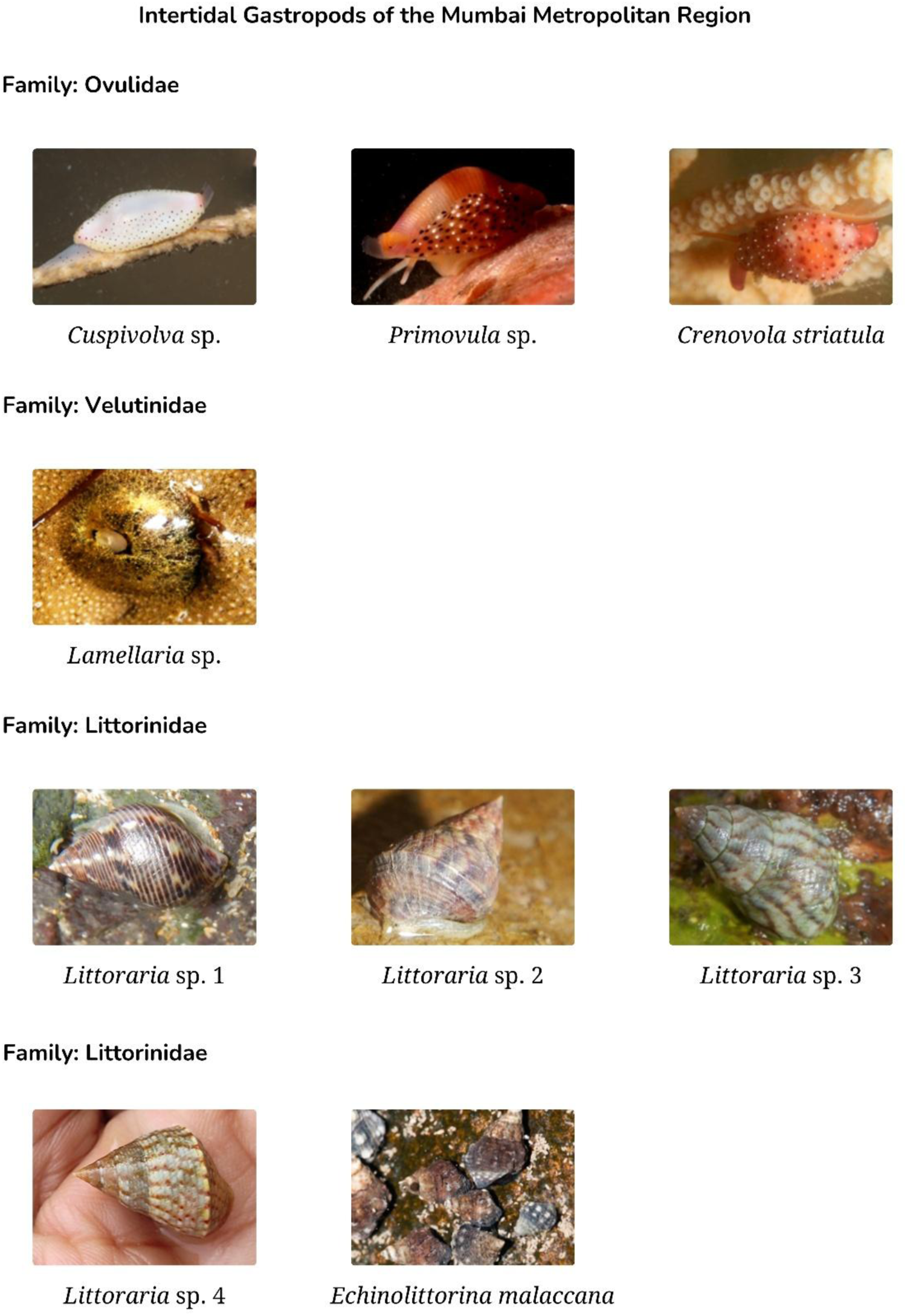

**PLATE 22.**
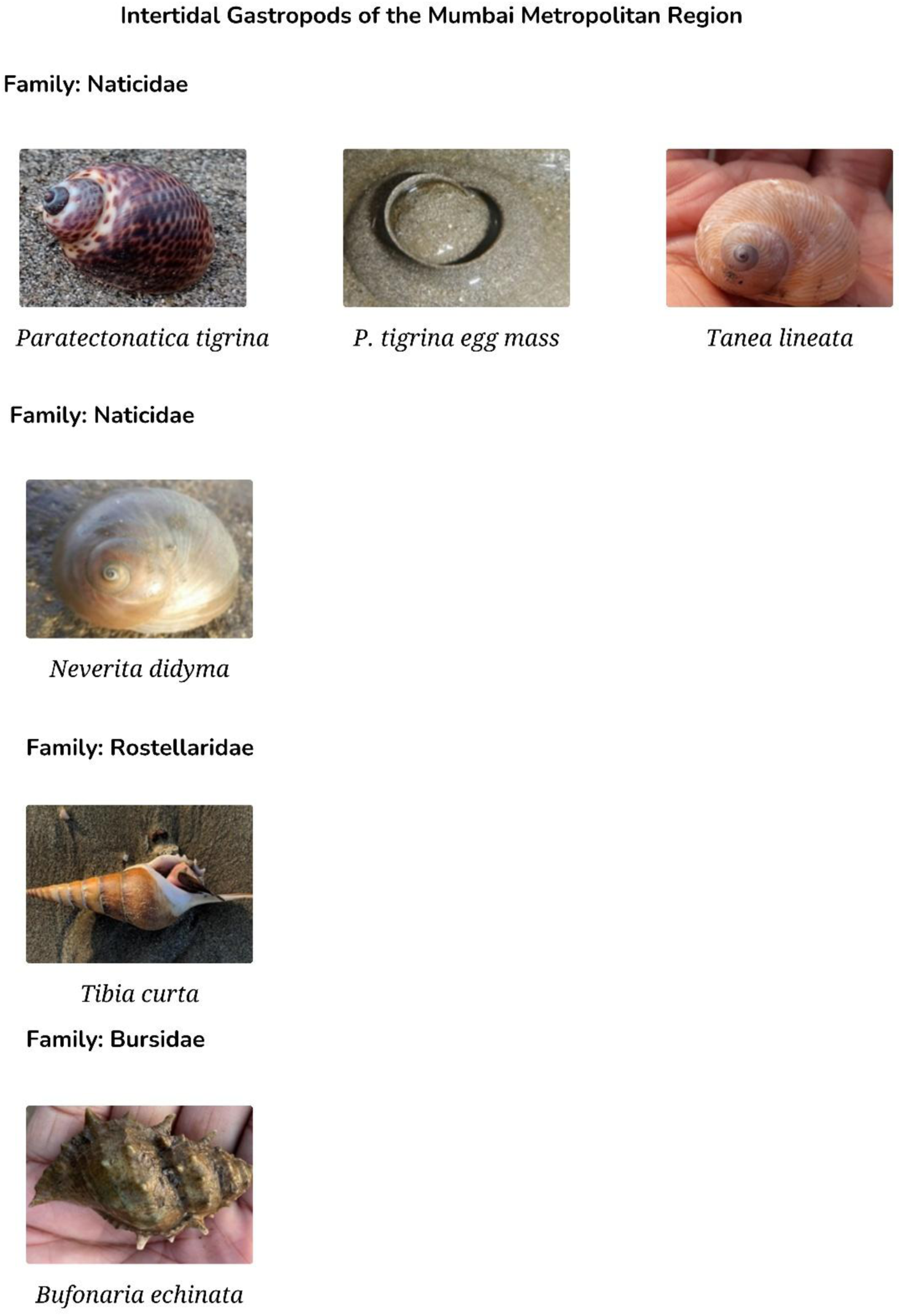

**PLATE 23.**
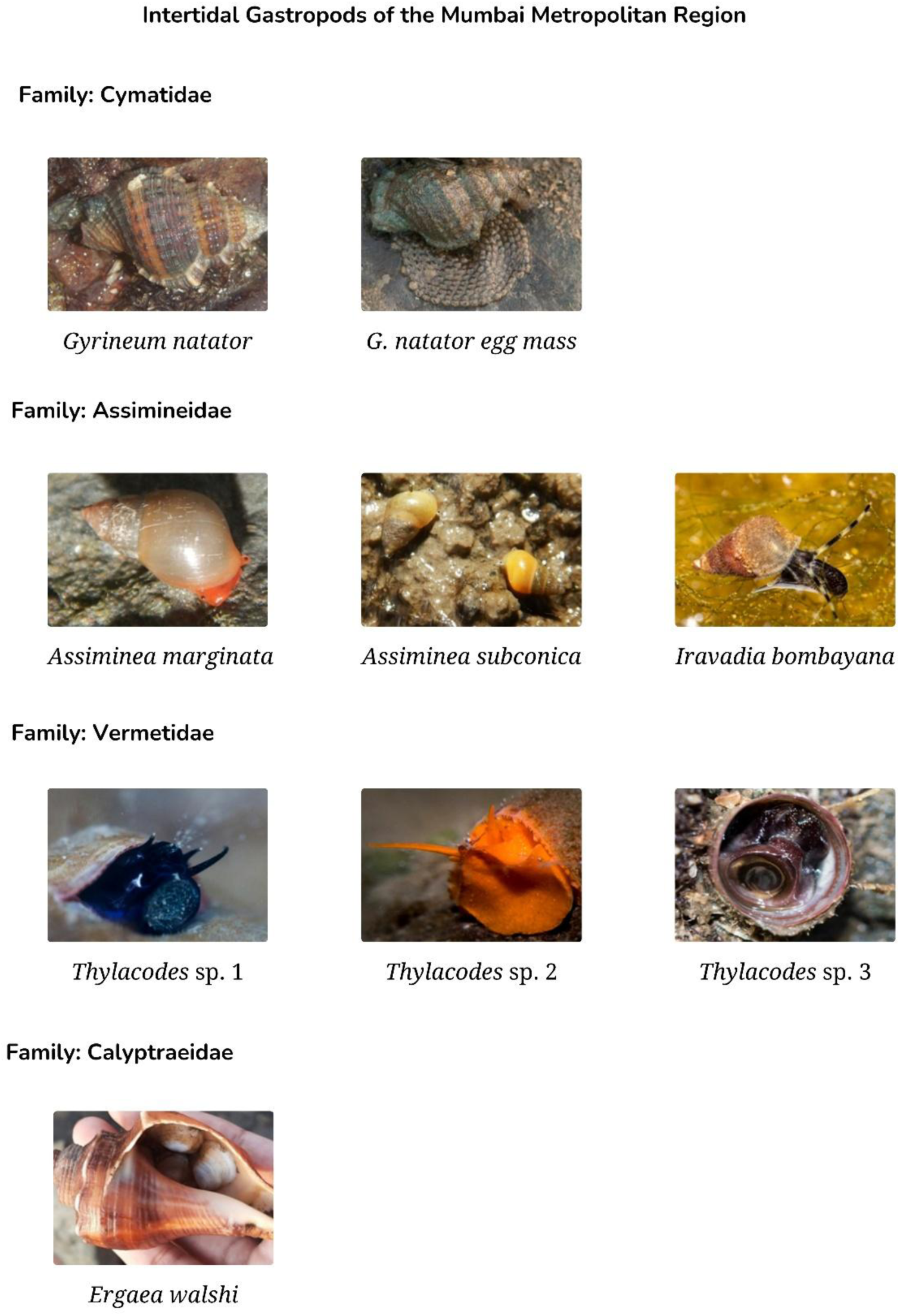

**PLATE 24.**
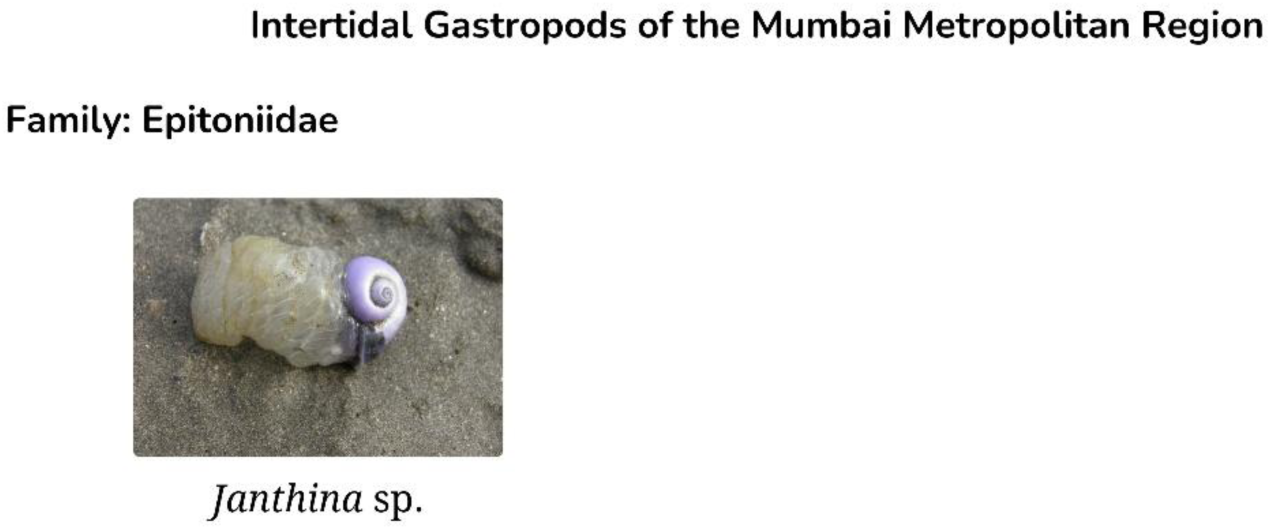

